# Arabidopsis uses a molecular grounding mechanism and a biophysical circuit breaker to limit floral abscission signaling

**DOI:** 10.1101/2022.07.14.500021

**Authors:** Isaiah W. Taylor, O. Rahul Patharkar, Medhavinee Mijar, Che-Wei Hsu, John Baer, Chad E. Niederhuth, Uwe Ohler, Philip N. Benfey, John C. Walker

## Abstract

Abscission is the programmed separation of plant organs. It is widespread in the plant kingdom with important functions in development and environmental response. In Arabidopsis, abscission of floral organs (sepals, petals, and stamens) is controlled by two receptor-like protein kinases HAESA (HAE) and HAESA LIKE-2 (HSL2), which orchestrate the programmed dissolution of the abscission zone connecting floral organs to the developing fruit. In this work, we use single-cell RNA-Sequencing to characterize the core *HAE/HSL2* abscission gene expression program. We identify the *MAP KINASE PHOSPHATASE-1/MKP1* gene as a negative regulator of this pathway. MKP1 acts prior to activation of HAE/HSL2 signaling to establish a signaling threshold required for the initiation of abscission. By analogy to electrical circuit control, we liken MKP1 to a molecular grounding mechanism that dissipates errant pathway activation absent HAE/HSL2 signaling. Furthermore, we use single-cell data to identify genes expressed in two sub-populations of abscission zone cells: those proximal and those distal to the plane of separation. We identify *INFLORESCENCE DEFICIENT IN ABSCISSION/IDA*, encoding the activating ligand of HAE/HSL2, as one of the mRNAs most highly enriched in distal abscission zone cells at the base of the abscising organs. We show how this expression pattern forms a biophysical circuit breaker whereby, when the organ is shed, the source of the IDA peptide is removed, leading to cessation of HAE/HSL2 signaling. Overall, this work provides insight into the multiple control mechanisms acting on the abscission-signaling pathway.

## Introduction

Abscission is a ubiquitous process occurring at multiple stages of the plant life cycle. Shedding of leaves in the autumn and fruit after ripening are common examples, as is shedding of organs after damage or infection^1–3^. Historically, selection for grass varieties with defects in seed abscission (also known as shattering) dramatically increased the efficiency of harvest and is considered one of the critical steps in domestication^4^. Controlling abscission today is still of agricultural and horticultural relevance in a wide range of species^5–8^.

In the genomics era, abscission of Arabidopsis floral organs is among the best studied systems^9^. In particular, Arabidopsis floral abscission has been shown to be regulated by ethylene, auxin, and their interaction^10–12^, with additional contributions from jasmonic acid and abscisic acid signaling^13,14^. In Arabidopsis, the site of floral abscission, known as the abscission zone (AZ), is lignified on the distal margin to create a well-defined separation plane^15^. During abscission, the middle lamella (the pectin-rich substance adhering adjacent cells) is degraded enzymatically before the organs detach^16^.

Abscission in Arabidopsis is regulated by two related leucine-rich repeat receptor-like protein kinases (LRR-RLKs) HAESA/HAE and HAESA LIKE-2/HSL2. The double mutant *hae hsl2* fails to shed its floral organs^17–19^. Several components of the *HAE/HSL2* signaling pathway regulating abscission have been discovered. In particular, HAE and HSL2 cooperate with members of the SOMATIC EMBRYOGENSIS RECEPTOR KINASE/SERK family of co-receptor LRR-RLKs to bind the proteolytic-processing derived INFLORESCENCE DEFICIENT IN ABSCISSION/IDA peptide, which activates a MAP kinase cascade^18,20–22^. MAPK signaling triggers expression of downstream abscission genes involved in pectin degradation, cell wall remodeling, and, ultimately, separation of the abscising organs^18,23–27^. A number of transcription factors have been implicated in regulating AZ development and abscission signaling ^14,28–31^. It has been shown that *Nicotiana* lines with silenced *HAE* and *IDA* orthologs exhibit reduced perianth abscission^32^, and expression of both citrus ad litchi orthologs of *IDA* can complement the abscission-deficient *ida* mutant phenotype in Arabidopsis^33,34^. We have also demonstrated abscission of Arabidopsis cauline leaves upon water stress or pathogen infection is under similar genetic control as floral abscission^35,36^. These results indicate the knowledge of the specific components of the *HAE/HSL2* pathway will inform regulation of abscission in multiple systems across multiple species.

While knowledge of positive regulators of Arabidopsis abscission signaling has been increasing in recent years, we know comparatively little about negative regulation of this process. This is a notable gap because plants must tightly control abscission signaling to avoid negative fitness consequences such as premature shedding of stamens and developing fruit. Moreover, prolonged abscission signaling could lead to wound formation, posing the risk of infection and desiccation. Among the negative genetic regulators we know include AGL transcription factors (such as AGL15 that has been shown to bind and negatively regulate the *HAE* promoter, repression of which is a component of a positive feedback loop), as well as the pseudo-kinase BIR1 (implicated in negatively regulating a SERK-SOBIR1 mediated signaling cascade)^20,26,37–39^.

In this work, we provide insight into how the abscission pathway is maintained at a low basal activity prior to abscission activation. We also describe an abscission termination mechanism that highlights how plants have evolved an elegantly simple way to halt the abscission process. We began our work by hypothesizing that assessing the transcriptional output of the *HAE/HSL2* pathway at the single-cell level could help us answer questions surrounding the dynamics of how abscission is orchestrated, and how the cell types within the AZ coordinate in this orchestration. This line of investigation led to the characterization of a mutant in a phosphatase gene that we show negatively regulates basal signaling prior to abscission pathway activation. Second, we establish how a spatial gradient in *IDA* expression contributes to limiting HAE/HSL2 signaling to the time when organs are attached. In essence, we demonstrate that IDA is a key switch that the plant has localized to the AZ cells on the abscising organ, ensuring that the trigger for the abscission pathway is physically removed from the plant once abscission has occurred. This creates a virtually infallible way of terminating abscission signaling, thus eliminating the potential hazards of uncontrolled pathway activation.

## Results

### Single-cell RNA-Sequencing identifies floral abscission zone cells

The genes encoding HAE and HSL2 are required for floral organ abscission in Arabidopsis [Figure 1A]. In order to characterize transcriptional outputs of *HAE/HSL2* signaling, we developed a flower receptacle protoplasting method (Materials and Methods) and used it to perform scRNA-Seq on stage 15 floral receptacles of WT and *hae hsl2* (2 replicates each) [Fig. 1B]. Stage 15 is the time when abscission signaling has been initiated, but WT and *hae/hsl2* are still morphologically comparable. We also profiled 2 replicates each of FACS sorted cells derived from *hae hsl2* + *HAEpr::HAE-YFP* (complemented) and kinase dead *hae hsl2* + *HAEpr::HAE-K711E-YFP* (non-complemented) receptacles to further enrich for AZ cells from phenotypically wild-type and mutant plants, respectively [Figures 1C and 1D] (Materials and Methods). In total, we obtained data for 4 WT/complemented (hereafter “WT”) and 4 mutant/non-complemented (hereafter “*hae hsl2*”) replicate samples each, granting us robust statistical power to identify genes regulated by *HAE/HSL2* signaling (Supplemental Table 1 and Supplemental Figure 1 for sample sequencing metrics). After preprocessing and filtering low quality cells, we integrated all cells using Seurat^40^, excluding genes whose expression was altered in a separate receptacle protoplast bulk RNA-Seq experiment (Materials and Methods). We used Louvain clustering to identify transcriptionally similar groups of cells and embedded them in a 2-dimensional Uniform Manifold Approximation and Projection (UMAP) [Figure 2A, top panel]. Next, we plotted WT cells on the UMAP expressing known markers of the AZ: *HAE* and the pectin degrading enzyme-encoding genes *QUARTET2/QRT2* and *POLYGALACTURONASE AZ OF A. THALIANA/PGAZAT* [Figure 2A, bottom panel]. A compact group of cells was apparent in the UMAP expressing all three AZ markers (Figure 1A, lower panels). These cells, numbering 869 out of a total of 16169 profiled WT cells (5.4%), grouped together across a wide range of Louvain clustering resolutions [Figure 2A, pink border, Supplemental Figure 2]. Thus, this group of cells appears to represent the AZ, which is transcriptionally identifiable and highly distinct from other cell types in the receptacle.

**Figure 1:**
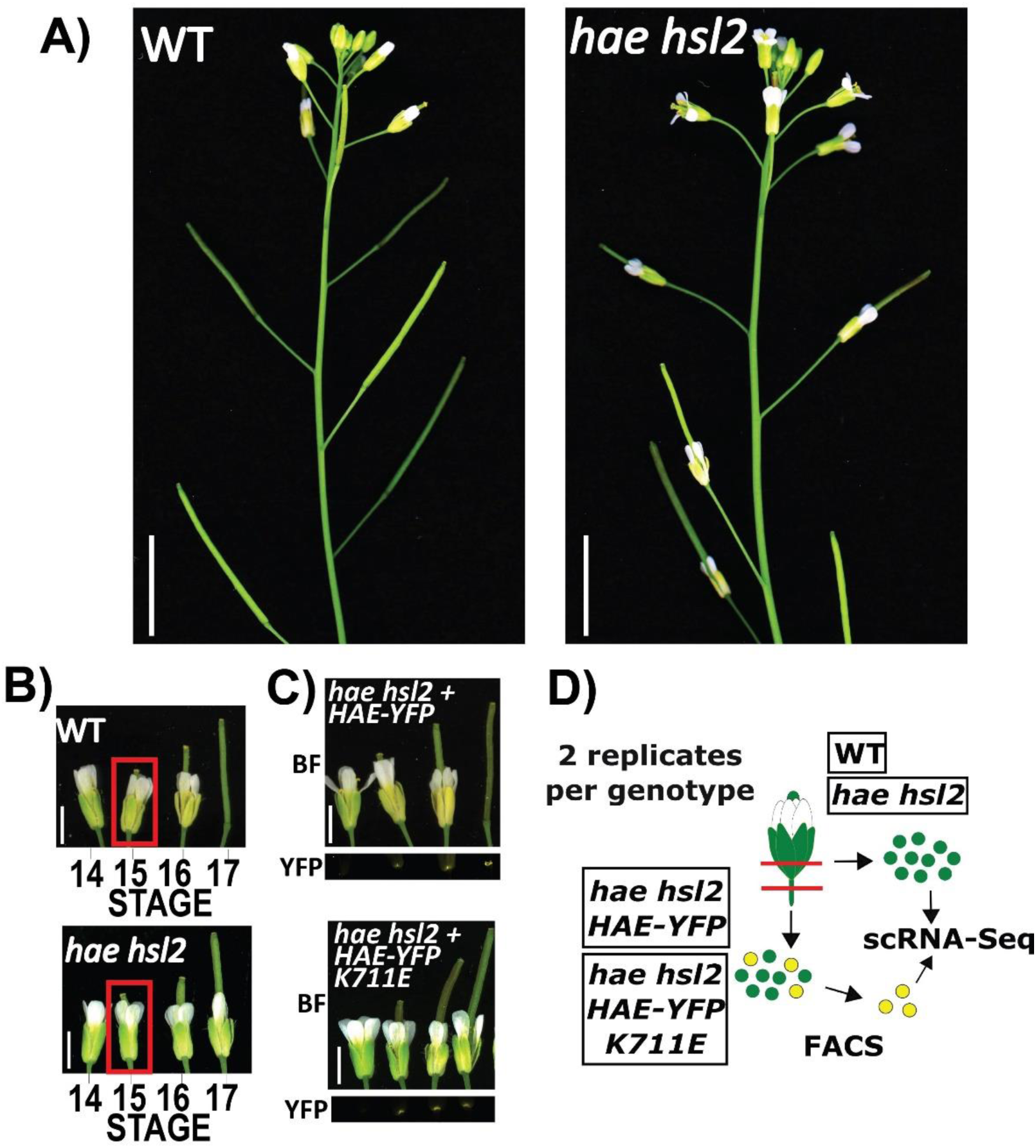
Background and experimental system. A) Floral abscission phenotype of WT and *hae hsl2* mutant. Scale bar = 10 mm. B) Flower stages in WT and *hae hsl2* mutant. Scale bar = 2 mm. C) Phenotype of transgenic *hae hsl2* expressing wild-type *HAEpr::HAE-YFP* and *HAEpr::HAE-YFP K711E* (bright-field/BF and YFP). Scale bar = 2 mm. D) Diagram of AZ single-cell isolation and experimental procedure.

**Figure 2:**
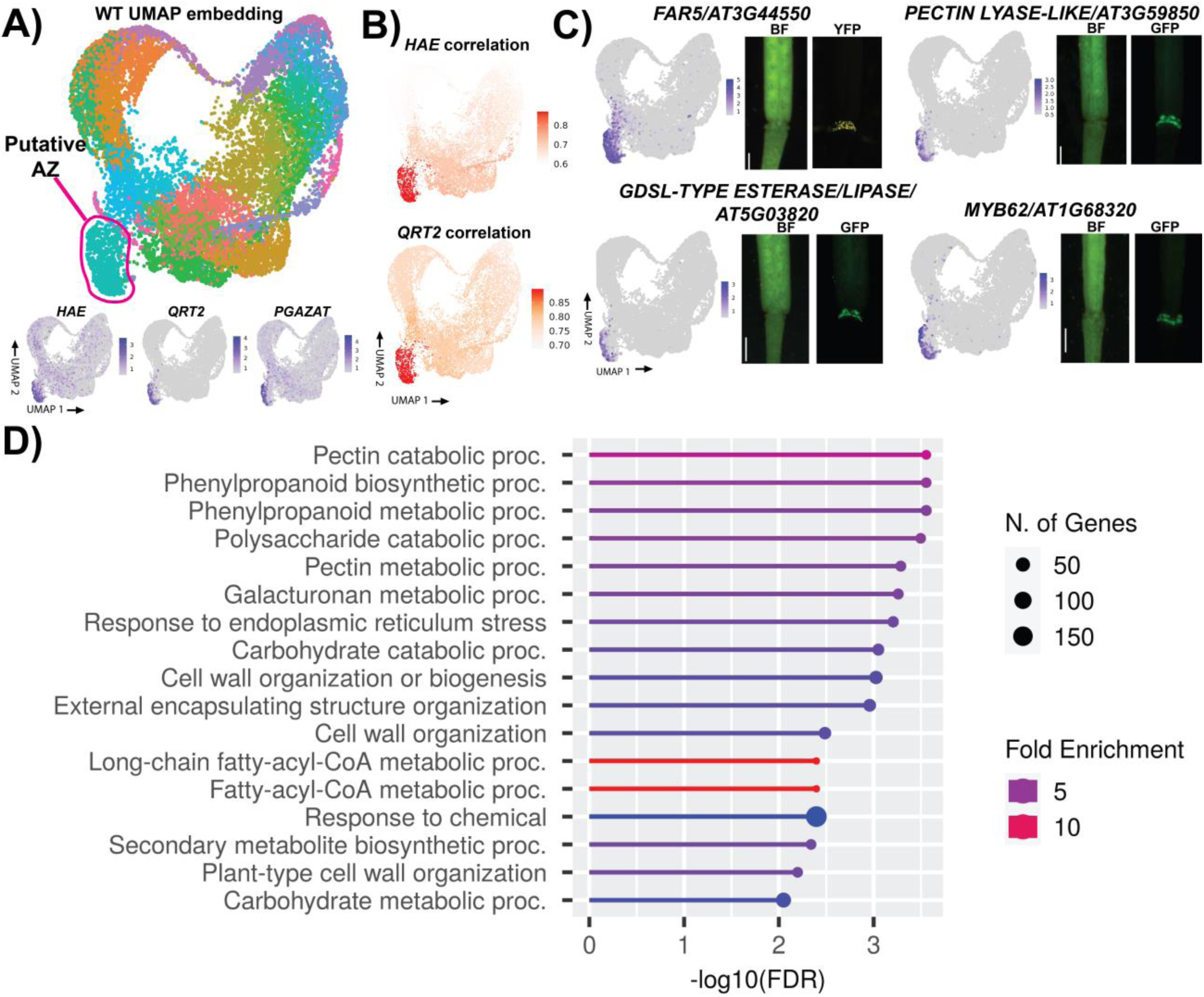
Identification and characterization of AZ cells by single-cell RNA-Sequencing. A) UMAP embedding of WT cells with putative AZ cluster circles in pink (top) and expression of AZ marker genes on UMAP (bottom). Expression is on Seurat SCT scale. B) Cluster-wise pseudo-bulk Spearman correlation with sorted bulk data of *HAEpr::HAE-YFP* (top) or *QRT2pr::GFP* (bottom). C) Expression of 4 putative AZ marker genes in single-cell UMAP embedding and in the young siliques of *promoter::fluorescent protein* expressing transgenic plants (bright-field/BF and YFP or GFP). Expression is on Seurat SCT scale. Scale bar = .7 mm. D) Gene Ontology Biological Process term enrichment of AZ specific genes.

To confirm the identity of the putative AZ, we took several approaches. First, we sorted *HAE-YFP* expressing cells and generated and sequenced bulk RNA-Seq libraries (Materials and Methods, Supplemental Table 2). We then calculated cluster-wise Spearman correlation between this dataset and pseudo-bulked data of each of the identified clusters from the single-cell experiment. The putative AZ cluster displayed the highest correlation, consistent with the hypothesis that it represents the AZ [Figure 2B, top panel]. Similarly, we used published bulk RNA-Seq data from FACS sorted *QRT2*-expressing stage 16 AZ cells to perform an identical correlation analysis and found that, again, this cluster displayed the highest correlation [Figure 2B, bottom panel, Supplemental Tables 3 and 4]^15^. Next, we identified genes enriched in the putative AZ cluster (not previously known to be specific to the AZ) and generated promoter::fluorescent reporter lines [Figure 2C]. All four reporters displayed highly specific expression in the AZ cells, indicating this cluster represents the AZ.

To characterize the biology of the putative AZ, we identified AZ enriched genes using Seurat (Materials and Methods, Supplemental Tables 5 and 6). Analysis of Gene Ontology (GO) terms found enrichment of genes associated with middle lamella degradation and cell wall remodeling (pectin catabolic process, polysaccharide catabolic process, galacturonan metabolic process, etc.) [Figure 2D]. Additionally, there were terms associated with extracellular barrier formation such as phenylpropanoid biosynthesis, the major pathway leading to lignin formation, which has been shown to play an important role in the delineation of the AZ in both Arabidopsis and rice^15,41^. Terms related to Fatty-acyl-CoA metabolism are associated with genes involved in cutin biosynthesis, a waxy substance deposited on the scar of newly differentiated epidermal cells after abscission has occurred^15^. Overall, these terms are highly consistent the AZ being a site of focused cell wall disassembly and barrier formation.

While not the focus of this study, we were also curious if we could identify other non-AZ cell types present in the receptacle. Indeed, many genes associated with specific cell types from Arabidopsis leaf scRNA-Seq data were present in restricted clusters in our data^42^, allowing us to make tentative assignments of epidermis, mesophyll, guard cells, etc [Supplemental Figure 3A-C]. This suggests that floral organs, as modified leaves, retain some of the gene expression patterns of their homologous organs. There were also clusters that did not express known leaf markers [Supplemental Figure 3B], likely representing less characterized, flower-specific cell types. For example, we identified a large group of cells expressing AT3G01420, an alpha dioxygenase encoding-gene previously shown to be highly enriched in bulk RNA-Seq of developing siliques^43^, suggesting that there are uncharacterized flower-specific cell types [Supplemental Figure 3D].

### The *hae hsl2* mutant exhibits a reduction in expression of genes associated with core abscission zone signatures

We next examined the scRNA-Seq data to characterize gene expression differences between *hae hsl2* and WT. Plotting mutant cells by UMAP indicates a similar spectrum of cell types as WT [Figure 3A], with a similar proportion of putative AZ cells numbering 964 out of 24497 profiled mutant cells (3.9%). To identify differentially expressed genes (DEGs), we performed pseudo-bulk analysis comparing WT to *hae hsl2* using edgeR^44^. This analysis revealed 302 genes with lower expression and 274 with higher expression in the double mutant as compared to WT [Supplemental Table 7].

GO term enrichment of genes with reduced expression in *hae hsl2* AZ was consistent with known biology of abscission, including terms related to pectin degradation and cell wall remodeling [Figure 3B, Materials and Methods]. Terms associated with suberin deposition are likely to reflect biosynthesis of cutin, a related wax compound with similar biosynthesis, which has been shown to accumulate in Arabidopsis floral AZs^15^. For visualization, we plotted the expression levels of a number of known and novel *HAE/HSL2* regulated genes [Figure 3C]. Overall, these results are consistent with the role of *HAE/HSL2* as central regulators of genes required for breakdown and remodeling of the cell wall during abscission. Interestingly, GO term analysis of genes with higher expression in *hae hsl2* did not reveal such clearly biologically interpretable signals and was enriched in terms associated with defense and hypoxia responses [Supplemental Figure 4]. This suggests there may be novel molecular pathways repressed by *HAE/HSL2* that can be interrogated in future work.

**Figure 3:**
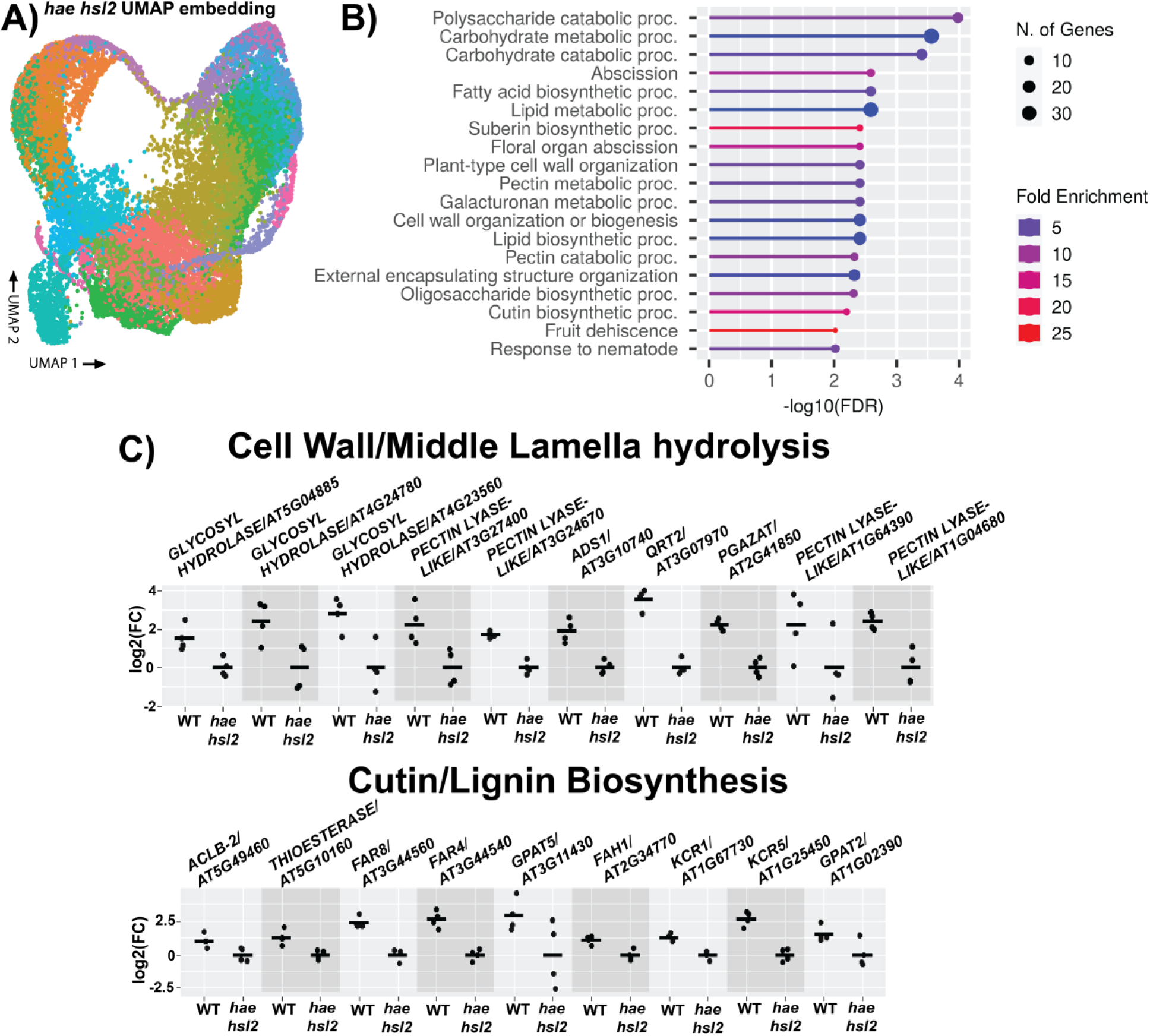
Analysis of differentially expressed genes in the *hae hsl2* mutant AZ. A) UMAP embedding of *hae hsl2* cells. B) Gene Ontology Biological Process term enrichment of DE genes higher in WT (FDR <.05, log2(FC) > 1). C) Expression of cell wall/middle lamella hydrolysis genes (top) and cutin/lignin biosynthesis genes (bottom). Scale is log2(FC) for each sample relative to the average of *hae hsl2*. All genes are lower in mutant with FDR < .05.

### *hae hsl2* suppressor mutants partially restore the abscission zone gene expression signature

Having identified a core set of genes regulated by *HAE/HSL2* specifically in the AZ, we set out to characterize this signaling pathway in two novel *hae hsl2* partial suppressor mutants isolated from a previously described *hae hsl2* T-DNA suppressor screen^26^. These mutants, which we named *fal-3* and *fal-7* for *facilitated abscission locus,* display a cool-temperature-enhanced suppression of the *hae/hsl2* phenotype when grown at 16° [Fig 4A]. This can be quantified by measuring the force required to remove petals of stage 16 flowers using a previously described petal break-strength assay [Fig 4B]^45^.

Based on the partial suppression phenotype, we hypothesized that gene expression changes in *hae hsl2* would be partially reversed in the *hae hsl2 fal-3* and *hae hsl2 fal-7* suppressors. To test this hypothesis, we performed stage 15 receptacle bulk RNA-seq of plants grown at 16° comparing WT, *hae hsl2*, *hae hsl2 fal-3* and *hae hsl2 fal-7* and examined the expression of 67 genes that were reduced both in *hae/hsl2* in the AZ in our single-cell data while also exhibiting a reduction in previously published receptacle bulk RNA-Seq experiments comparing WT and *hae/hsl2* [Supplemental Tables 8 and 9]^26,27^. Thus, these genes represent an AZ-specific, validated set of genes regulated by *HAE/HSL2* identifiable from bulk analysis [Supplemental Figure 5]. Indeed, in *hae hsl2 fal-3* and *hae hsl2 fal-7,* most of these genes show an intermediate level of expression between WT and *hae hsl2* [Figure 4C]. To perform a single composite statistical test, we used Parametric Analysis of Gene Expression/PAGE, which averages expression of a set of genes on a log scale and uses the resulting approximate normality to perform comparisons^46^ (Materials and Methods). Consistent with the hypothesis that the suppressors have partial reversion of the abscission gene expression program, the average log2(FC) levels for both *hae hsl2 fal-3* and *hae hsl2 fal-7* are intermediate between WT and *hae hsl2* [Figure4D]. These results confirm the partial suppression phenotype is due to partial *HAE/HSL2* pathway activation in these mutants.

**Figure 4:**
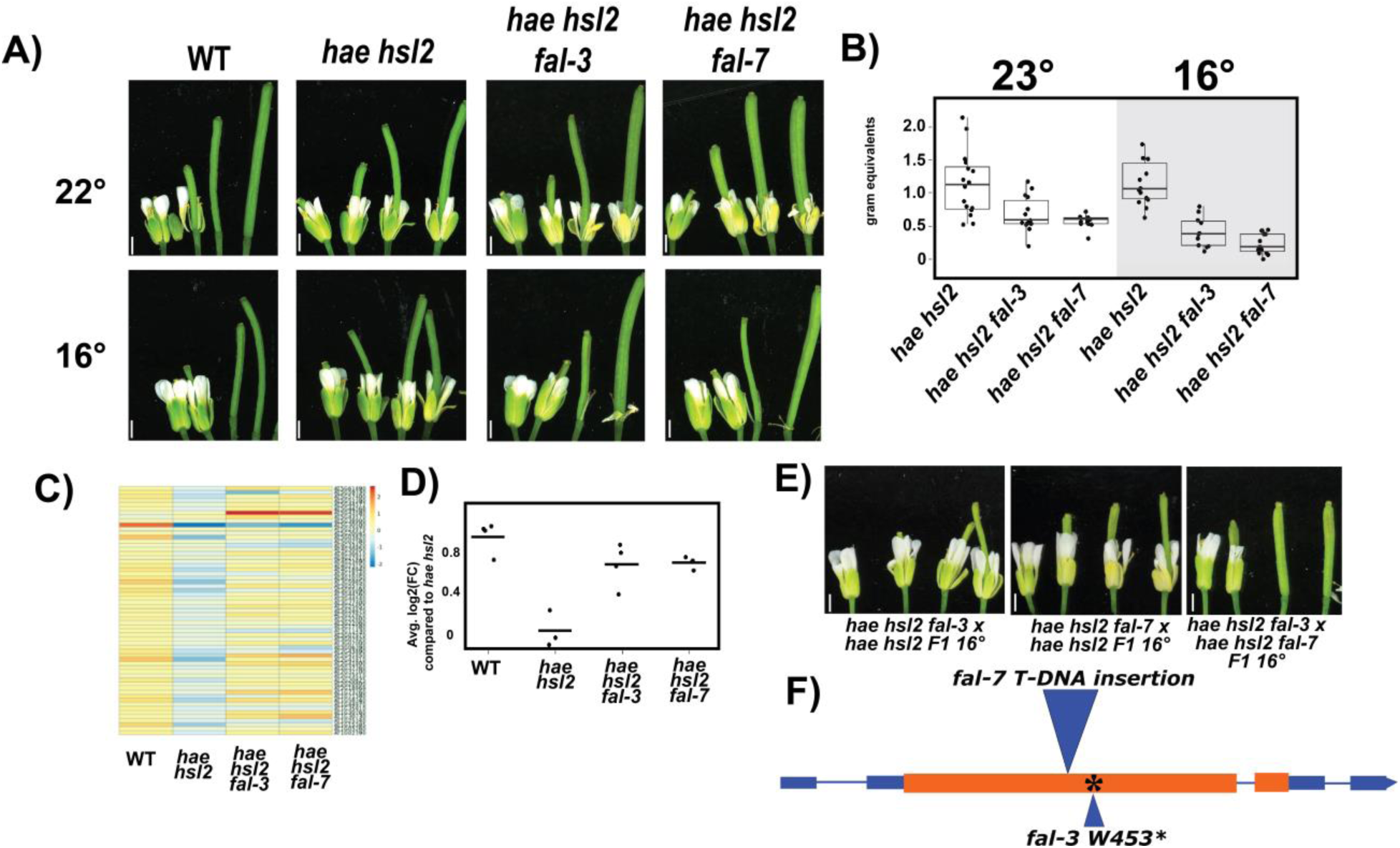
Analysis and identification of *hae hsl2* suppressors. A) Phenotypes of WT, *hae hsl2, hae hsl2 fal-3*, and *hae hsl2 fal-7* at 22° (left) or 16° (right). Siliques were gently tapped to remove remnant floral organs. Scale bar = 1 mm. B) Breakstrength phenotypes of *hae hsl2, hae hsl2 fal-3*, and *hae hsl2 fal-7* at 23° (left) or 16° (right). C) Heatmap of expression values for 67 genes identified as higher in WT in both bulk and single-cell DE analysis comparing WT to *hae hsl2.* Values are log2(FC) compared the overall average expression across all genotypes. D) Average log2(FC) comparing each sample to the average of *hae hsl2* for the genes in part C. E) Complementation crosses of *fal* mutants. Scale bar = 1 mm. F) Gene model of *MKP1* depicting mutations in *fal-3* and *fal-7.* Orange colors represent exons, blue represent UTRs, and thin lines represent introns.

### Mutation of *MAP KINASE PHOSPHATASE-1/MKP1* underlies the *hae hsl2 fal* suppression phenotype

To identify the causative mutations in *fal-3* and *fal-7* we performed complementation crosses, which indicated that *fal-3* and *fal-7* are allelic, recessive mutations [Figure 4E]. To identify the underlying mutations, we performed TAIL-PCR and found *fal-7* harbors a T-DNA insertion in the first exon of *MAP KINASE PHOSPHATASE-1/MKP1,* a gene encoding a phosphatase known to negatively regulate MPK3 and MPK6 during biotic and abiotic stress signaling in a cool-temperature enhanced manner [Figure 4F]^47,48^. Because MPK3/6 are also involved in abscission signaling^18^, and because the effect of *fal-3* and *fal-7* is cold-enhanced, *MKP1* became a very strong candidate gene. In *fal-3*, we identified a SNP causing a premature stop codon in the first exon of *MKP1* leading to truncation of nearly half the protein [Figure 4F]. In addition, segregation analysis of a back-cross population of *hae hsl2 x hae hsl2 fal-7* indicated the *hae hsl2* suppression effect is linked to the insertion in *MKP1* [Supplemental Figure 6] confirming that the phenotype of *fal-3* and *fal-7* is due to mutation of *MKP1*.

These results together indicate MKP1 is a negative regulator of abscission that buffers low levels of basal signaling which could otherwise induce abscission in the absence of HAE/HSL2 activation. This molecular thresholding mechanism is likely critical to prevent errant signal amplification of background noise and to ensure coherent activation of downstream signaling across the AZ. By analogy, this resembles the safety mechanism of grounding in electrical systems where the neutral and ground wires in the main electrical panel are bonded to set neutral voltage to the reference ground value of zero. This allows unwanted energy a dissipation path, preventing unwanted voltage noise on inactive circuits. In a similar fashion, MKP1 eliminates spurious MAPK phosphorylation that could lead to premature pathway activation prior to abscission signal induction.

### Single-cell RNA-Seq identifies spatial domains of the abscission zone

While the above analysis relied on pseudo-bulk measurements of gene expression across the entire AZ, we were next interested to explore whether we could potentially obtain more fine-grained information on the function of distinct cell populations within the AZ. An important report from Lee et al^15^ provided a detailed examination of the spatial organization of the AZ, identifying a differentiated group of cells on the distal side at the base of the abscising organ (termed *secession cells*) and a distinct group on the proximal side of the abscising organ (termed *residuum cells*) [Figure 5A]. Functionally, secession cells form a lignin “brace,” which is thought to focus the enzymatic activity of secreted hydrolases to the middle lamella of the fracture plane at the site of abscission, while also creating a rigid frame which detaches once sufficient weakening of the fracture plane has occurred.

**Figure 5:**
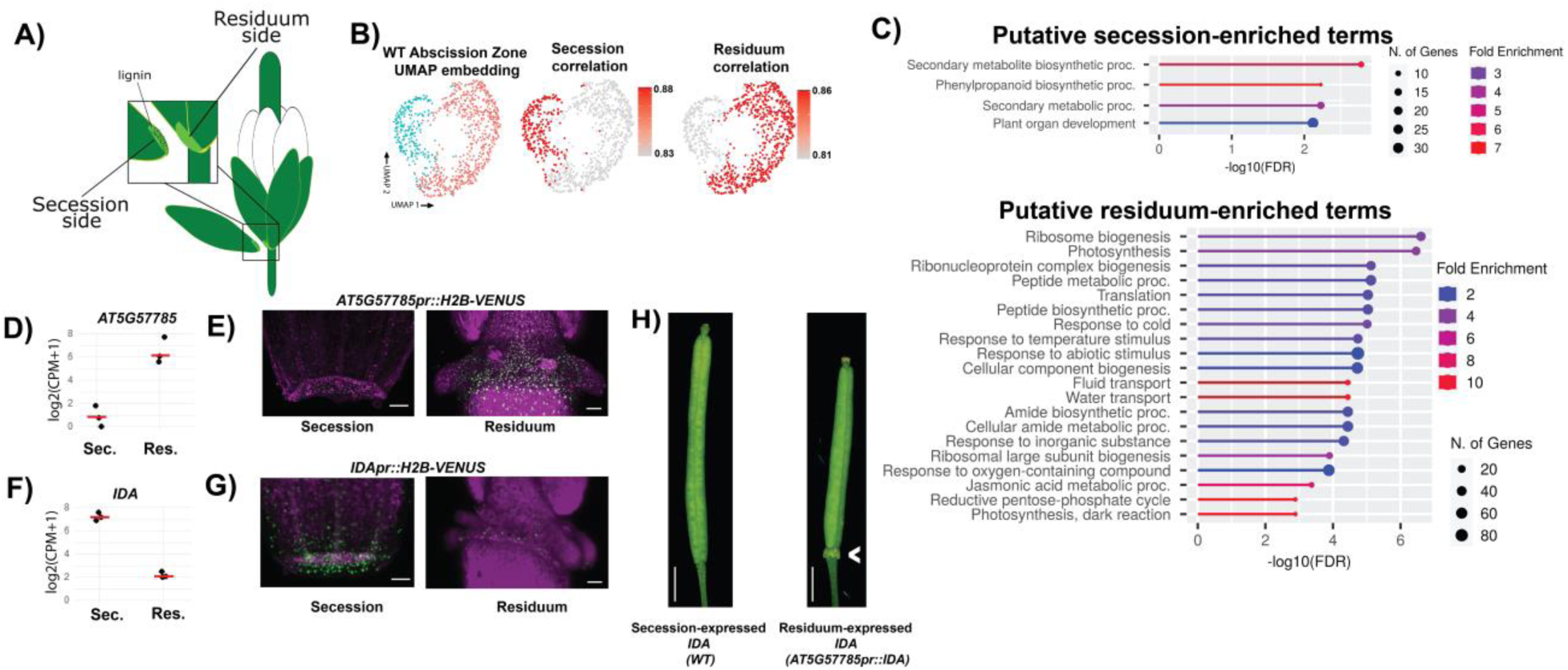
Spatial analysis of the AZ. A) Schematic view of the spatial organization of the AZ. B) Low resolution Louvain clustering and UMAP embedding of WT AZ cells (left) and Spearman correlation with sorted bulk secession data (middle) or residuum data (right). C) GO Term enrichment of genes differentially enriched in the putative secession cells (top panel) or putative residuum cells (bottom panel). D) Pseudobulk log2(CPM + 1) of *AT5G57785* in putative secession (“sec.”) and residuum (“res.”) cells. E) *AT5G57785pr::H2B-VENUS* expression in secession and residuum cells. F) Pseudobulk log2(CPM + 1) of *IDA* in putative secession and residuum cells. G) *IDApr::H2B-VENUS* expression in secession and residuum cells H) *IDA* expression phenotype in WT (left) and residuum specific *AT5G57785pr::IDA* misexpression phenotype (right). I) Low resolution Louvain clustering and UMAP embedding of *hae hsl2* AZ cells (left) and Spearman

This process had previously been detailed in stage 16 flowers, just as abscission is occurring^15^. We hypothesized we may be able to detect evidence of the differentiation process at stage 15 from our data. Interestingly, when performing low resolution Louvain clustering of WT AZ cells and embedding in UMAP space, two groups of cells emerge [Figure 5B, left panel]. We tested the hypothesis that these cells represent secession and residuum cells by performing cluster-wise Spearman correlation analysis with previously published bulk RNA-Seq data derived from sorted secession cells and, separately, sorted residuum cells^15^. Consistent with our hypothesis, we observed a strong association of one of the two clusters of cells with secession bulk data [Figure 5B, middle panel] and strong association of the second group of cells with residuum bulk data [Figure 5B, right panel].

To validate the hypothesis that these cell clusters represent distinct sides of the AZ, we performed a pseudo-bulk differential enrichment analysis to identify genes enriched in the putative secession and residuum cells. We identified 306 and 540 genes displaying evidence of enrichment in these cell types, respectively [Supplemental Table 10]. GO term enrichment analysis of the secession cell-associated genes in revealed modest enrichment of pathways related to phenylpropanoid biosynthesis [Figure 5C, top panel]. Since one of the main functions of the phenylpropanoid pathway is lignin formation, this is consistent with the idea that putative secession cells are in the early stages of forming a lignin brace. In contrast, the genes associated with putative residuum cells are enriched in terms related to cellular activity such as protein production [Figure 5C, lower panel]. This is consistent with classical observations that AZ cells are sites of high levels of protein synthesis^49^. Interestingly, we performed a similar analysis of putative secession and residuum cells in mutant samples, and we found evidence of significantly fewer genes enriched in the mutant cell types, suggesting *HAE/HSL2* may drive cellular differentiation of these two cell types within the AZ [Supplemental Figure 7 and Supplemental Table 11].

To provide final validation of the identification of AZ sub-types, we identified one gene, *AT5G57785*, encoding a predicted mitochondrial protein of unknown function, as a gene highly enriched in the putative residuum side of the AZ [Fig 5D]. Promoter::reporter analysis confirmed this expression pattern, showing the gene is relatively low-expressed in secession cells, but highly expressed across the residuum [Figure 5E, Supplemental Table 10]. In contrast, and intriguingly, we found one of the most highly enriched genes in the putative secession cells is *IDA*, encoding the activating ligand of HAE/HSL2 [Figure 5F, Supplemental Table 10]. We confirmed this pattern by generating promoter::reporter lines, demonstrating strong expression in secession cells and only patchy expression in residuum cells [Figure 5G]. These results confirm the ability of single-cell RNA-Seq to discriminate AZ sub-types.

### Localization of *IDA* expression to abscising organs terminates *HAE/HSL2* signaling after organ separation

Finally, we noted that it is interesting that while *IDA* is highly enriched on the secession side of the AZ, other signaling components such as *HAE, HSL2,* and *MKP1* are detected at similar levels in both residuum and secession cells [Supplemental Figure 8]. This specific enrichment of *IDA* in secession cells suggested to us a potential elegant regulatory control mechanism. In this model, the soluble IDA peptide is secreted by secession cells to activate signaling locally, at the site of organ attachment, while diffusing across the secession-residuum boundary to activate signaling in residuum cells in trans. However, once cell separation occurs, the source of the activating IDA ligand (i.e., the base of the abscising organ) is physically detached from the plant, leading to signal attenuation.

One prediction of this model is that if *IDA* were mis-expressed in the residuum cells we might observe some evidence of HAE pathway over-activation. To test this prediction, we expressed the *IDA* gene in residuum cells by fusing it to the residuum specific *AT5G57785* promoter fragment characterized above. Indeed, we observed plants displaying highly enlarged, disorganized AZs [Figure 5G]. This phenotype has previously been observed in cases where the *IDA-HAE/HSL2* pathway is over-activated, for instance through general over-expression of *IDA* or mis-expression of the activating TF *WRKY57* ^18,19,30^. This result indicates that localizing *IDA* on the secession side of the AZ ensures the clean and abrupt cessation of signaling that occurs in WT after abscission has occurred. Importantly, we have previously shown over-expression of *HAE* does not lead to such a pathway over-activation phenotype, instead yielding phenotypically WT plants^50^. Thus, these results together demonstrate a novel mechanism by which the physical removal of the source of IDA is likely the main signal attenuation mechanism to deactivate signaling once abscising organs are shed. A diagram of the model of abscission signal control is shown in Figure 6.

**Figure 6:**
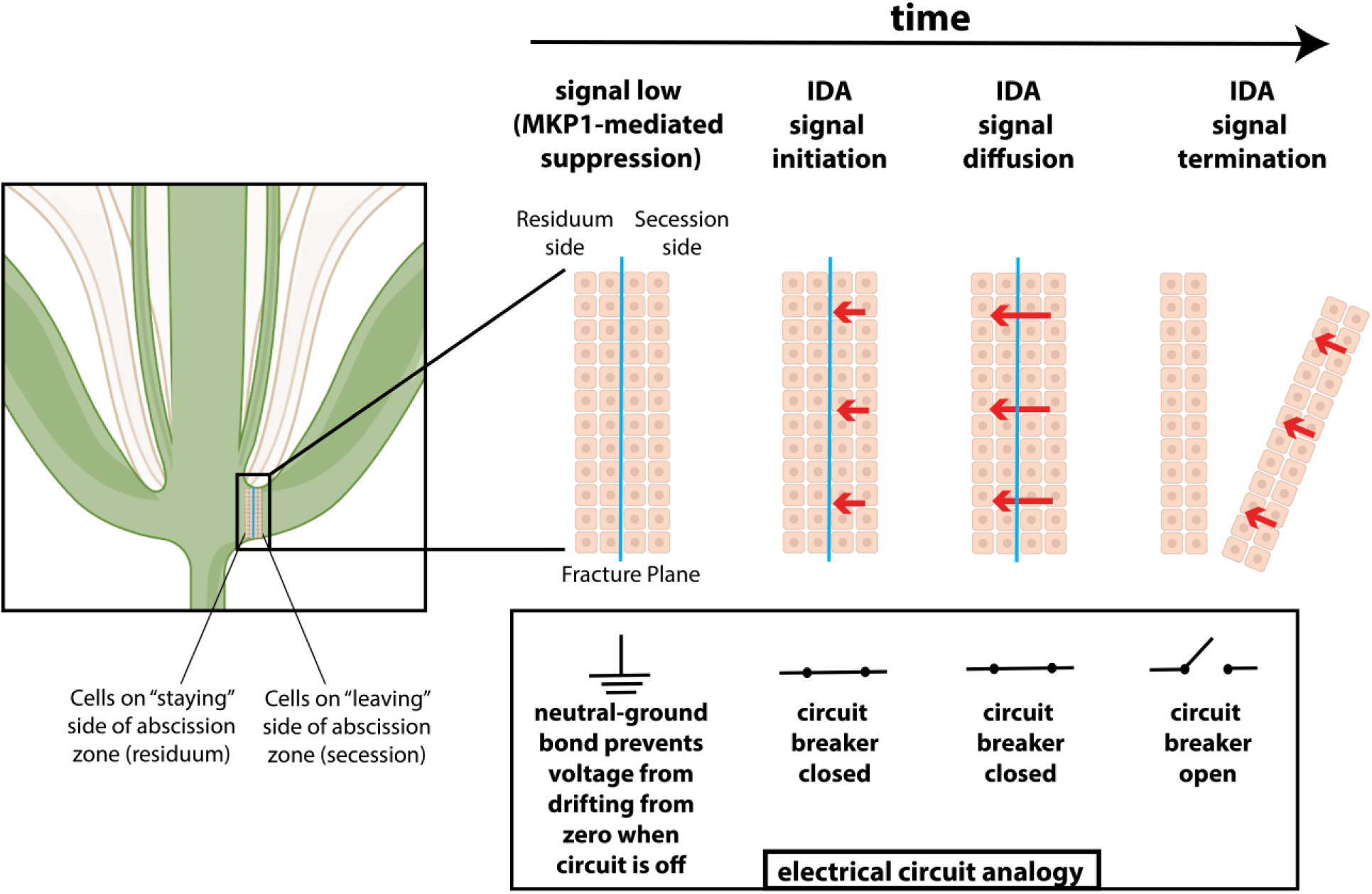
Diagram depicting the molecular signaling circuit controlling abscission signal initiation and termination.

## Discussion

This work highlights two important emergent properties of abscission signaling. First, the pathway must be tightly regulated to prevent signaling absent inducing stimulus which might lead to premature abscission (exemplified here by MKP1 maintaining basal signaling below a threshold level prior to pathway activation). This could be necessary since the abscission pathway is forcefully activated by positive feedback, placing the pathway under the control of a molecular hair trigger^20^. Second, activation of the pathway should be restricted to the site of organ attachment and be turned off after abscission has occurred. This avoids widespread and prolonged signaling leading to wounding. Our work shows that during Arabidopsis floral abscission, this is accomplished by localizing the source of the inducing signal IDA to the abscising organ. The ingeniously designed abscission switch constitutes a failsafe biophysical circuit breaker that terminates a highly amplified abscission activation signal, along with the potentially deleterious cellular processes that result from it. Diffusion of IDA peptide from source secession cells establishes a gradient across both sides of the abscission zone that does not appear necessary for abscission to take place. However, mis-expression of *IDA* in residuum cells, creating an altered IDA gradient, results in malformed residuum abscission scars, demonstrating the regulatory logic of restricting *IDA* expression to the AZ cells on the abscising organ. We propose both highlighted mechanisms are likely general features of the regulation of abscission across species and organ types.

It is interesting to note that Thomas Edison invented the electrical circuit breaker in 1879 to prevent fires arising from electrical short-circuits and overloading^51^. Nature appears to have solved a similar problem over the course of millions of years of evolution of abscission in plants. Equipping abscission signaling circuitry with both a physical circuit breaker to terminate the process and a noise reduction mechanism, resembling electrical grounding, to keep the circuitry completely off when not needed, illustrates how simple, yet ingenious and effective, plant signaling system design can be.

## Methods

### Plant growth

The *hae-3 hsl2-3* mutant was used in all experiments^52^. Wild type, *hae-3 hsl2-3, hae-3 hsl2-3 HAEpr::HAE-YFP, hae-3 hsl2-3 HAEpr::HAE-YFP K711E, hae-3 hsl2-3 fal-3,* and *hae-3 hsl2-3 fal-7* are all in the Columbia background. The *fal* suppressor mutants also contain *erecta* and *glabrous* mutations in order to phenotypically differentiate contaminating wild-type seeds from mutants in the *hae hsl2* suppressor screen^26^. Plants were grown in a 16-hour light cycle at either 22° or 16°, except for the breakstrength experiments which were conducted at 23° and 16°. Plants were grown in peat moss/vermiculite potting soil and fertilized every 3 weeks with ½ strength Miracle-Gro (Scotts Miracle-Gro Company).

### Protoplasting and FACS

For each protoplast sample, we isolated approximately 20 1 mm sections of stage 15 floral receptacles of plants approximately 2 weeks post-bolting, in a similar manner as our previous work^26,53^. For protoplasting, we employed the isolation method of Evrard et al, developed for rice roots, with minor modifications^54^. In brief, we prepared digest solution as follows: per 30 mls we added 400 mM mannitol (2.2 g), 20 mM MES hydrate (117 mg), 20 mM KCl (600 μL of 1 M KCl), after which the pH was adjusted to 5.7 with KOH. We would typically make 500 mls of this solution, filter sterilize, and store at 4° for several months. On the day of protoplasting, per 30 mls, we added 1.25% Cellulase R10 (375 mg), 1.25% Cellulase RS (375 mg), 0.3% Macerozyme R10 (90 mg), 0.12% Pectolyase-Y23 (36 mg) and heated 10 min in 60° water bath before cooling to room temperature (all digest enzymes purchased from Duchefa Biochemie except Pectolyase-Y23, which was purchased from MP Biomedicals). Finally we added 10 mM CaCl2 (300 μL of 1 M CaCl2), 0.1% BSA (30 mg), and 5.38 μL of β-mercaptoethanol. We sliced the receptacles into quarters under a dissecting scope on 4% agarose plates with a fine micro-scalpel #715 (Feather), used forceps to transfer into 5 mls of digest solution in small petri dishes, and vacuum infiltrated for 7 minutes at -25 inHg in a dome desiccator. We digested the tissue for 2.25 hours at 80 RPM on a rotary platform shaker at 25°, using fine forceps and micro-scalpel to additionally slice the softened tissue at around 1.5 hours. We next filtered the cells twice through 40 um filters, spun for 5 minutes at 500 x g in a swinging bucket rotor in 5 ml sorting tubes, and rinsed twice with wash buffer (digest buffer with no enzymes). Between washes we spun cells for 3 minutes at 500 x g. Finally, we resuspended cells in wash buffer. It should be noted that it is now recognized calcium in the resuspension buffer reduces the efficiency of reverse transcription^55,56^, so it is recommended that for follow-up studies, at least in the final resuspension step, calcium be omitted^56^. For receptacle single-cell samples, cells were counted on a hemacytometer C-chip (SKC, Inc.) and the concentration adjusted to 1000 cells/μl (total yield ranged from approximately 50,000-100,000 cells). For sorted samples, cells were prepared the same manner and sorted for YFP+ on a BD Diva cell-sorter into either excess wash buffer (for single-cell samples) or directly into RNA-Later for bulk samples. Single-cell sorted samples were then spun down and resuspended in 15 μl wash buffer. We took 2.5 μl of cells for dilution and counting on a hemacytometer C-chip. Final yield for sorted samples was between 400-3000 cells, all of which were run on a Chromium chip.

### scRNA-Seq library generation

We used the 10x Genomics v3 3’ Single-cell RNA-Seq kit for all samples. For the receptacle WT and *hae hsl2* samples we followed the manufacturer’s protocol, loading 16,000 cells per sample for a target capture of 10,000 single cells. For sorted samples, we divided a single reaction into four 25 μl reactions and ran on four lanes of a single chip. For sorted sample library preparation, we reduced the volume of all reagents only by ½, which was enabled by excess reagents accumulated due to prior emulsion failures (although we expect miniaturization to ¼-scale library preparations would be feasible). Raw reads have been deposited at the Sequence Read Archive under BioProject PRJNA857332.

### scRNA-Seq library sequencing and preprocessing

Unless otherwise noted, all analyses were performed in R 3.6.3 and are included, along with the output of sessionInfo(), as Jupyter Notebooks in the Supplemental Code. We used our previously published scKB procedure to align and produce count matrices for downstream analysis^57^. This pipeline uses kallisto, bustools, busparse, and BSgenome^58–61^ to align and quantify counts to the Arabidopsis TAIR10 genome. The following analyses are recorded in Notebook 1: For samples WT and *hae hsl2* receptacle samples #1 we had performed species-mixed experiments containing rice and Arabidopsis cells, so we aligned to a concatenated rice-Arabidopsis MSU7/TAIR10 genome using a combined gff file and retained only reads mapping to Arabidopsis^50,52^. We pooled reads mapping to spliced and unspliced transcripts in order to make a single matrix of gene expression values. We next ran EmptyDrops^63^ in order to identify putative empty droplets containing no cells with “ignore” parameter = 500 and “lower” parameter = 300. We then constructed Seurat objects with the expression matrices. We finally used doubletFinder^64^ to identify putative doublets using the approximate doublet rate employed by 10X as .004/500 * # loaded cells. The resulting Seurat objects were used for downstream analysis. Our estimated number of recovered cells after all these steps are: WT receptacle #1: 7509 cells, WT receptacle #2: 7571 cells, WT sorted #1: 878 cells, WT sorted #2: 211 cells, *hae hsl2* receptacle #1: 14492 cells, *hae hsl2* receptacle #2: 7578 cells, *hae hsl2* sorted #1: 1302 cells, *hae hsl2* sorted #2: 1125 cells.

### Exploratory analysis of scRNA-Seq data and identification of the AZ

Sample integration (Notebook 2) was performed by running SCTransform on each sample before integrating, excluding mitochondrial genes, plastid genes, and genes altered by protoplasting (defined as those with absolute value of log2(FC) > 1) [Supplemental Table 12]. We performed PCA, constructed a shared nearest neighbor graph, and identified clusters using the SLM algorithm. Visualization was performed by UMAP embedding.

Identification of the AZ (Notebook 3) was performed by first plotting *HAE*, *QRT2*, and *PGAZAT* in WT cells of the UMAP embedding with the “min” parameter set to .5. We next calculated the pseudobulk expression profile for all clusters identified at resolution .75 by summing all counts for each gene. We next calculated Spearman correlation with bulk sorted *HAE+* and *QRT2+* RNA-Seq data. The *HAE* data (Supplemental Table 2) was Lexogen Quant-seq 3’ and consequently is directly convertible to Transcripts per Million (TPM) making it comparable to 3’ RNA-Seq data generated by Chromium 10X. Because the *QRT2* data (Supplemental Table 3) include separate secession and residuum derived cells, we first created an approximate composite AZ transcriptome by summing the counts per million for both secession and residuum datasets with equal weighting. This dataset was full-transcript RNA-Seq, so we estimated TPM using the formula below (Supplemental Table 4):

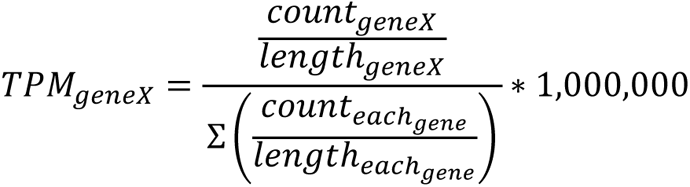

We then plotted the correlation value for each cluster on the previously generated UMAP embedding.

### Promoter cloning and reporter imaging

Fragments ranging in size from 1 to 2.5 kilobases upstream of *MYB62*, *PECTIN LYASE-LIKE*, and *GDSL-TYPE ESTERASE* were PCR amplified with PfuUltraII polymerase and cloned into pENTR (ThermoFisher), then recombined into pMDC111^65^ to create promoter::GFP fusions (primers are listed in Supplemental Table 13). The *FAR5, IDA,* and *AT5G57785* upstream regions were PCR amplified by KAPA Hifi polymerase (Roche) and cloned into an *Oryza sativa* H2B-VENUS fusion construct created by gene synthesis (Twist Bioscience). The *H2B-VENUS* construct was first cloned into pENTR and recombined into pGWB501^66^. This construct was created to include two AarI sites for Golden Gate cloning of promoter fragments in one step (sequence in sup). The *AT5G57785pr::IDA* misexpression construct was generated by cloning a synthetic *IDA* gene containing an upstream AarI cassette into pGWB501 [Twist Biosciences]. This allowed us to directory clone the *AT5G57785* promoter fragment above into this vector (sequence in sup). Primers are included in Supplemental Table 13. Siliques for the initial AZ identification experiment were imaged on a Zeiss Axiozoom stereomicroscope with UV illumination. *IDA* and *AT5G57785* reporter lines were images using a Zeiss LSM880 microscope with an EC Plan-Neofluar 10x/0.30 M27 objective. The lasers were 561nm and 488nm, with filters of 508-561nm and 570-670nm. Beam splitter information is as follows: MBS: MBS 488/561, MBS_InVis:Plate, DBS1: Mirror. Each set of images (residuum and secession AZs) were imaged from the same flower using the same settings.

### Non-Abscission zone Cell Annotation

Clusters were defined by twice running the Seurat FindClusters function with both a low modularity parameter (res = 2) and a high modularity parameter (res = 200), which results in clusters containing hundreds/thousands of cells (broadly-resolved) and those having only tens of cells each (finely-resolved), respectively. We then calculated expression z-scores of known marker genes [Supplemental Table 14] in each cluster (both broadly and finely-resolved). These clusters were then annotated by comparing the average marker gene z-scores. Cells that were annotated with the same cell identity in broadly-resolved and finely-resolved clusters were considered confidently annotated. While those that were not were labeled as “Unknown.” This annotation approach combining results of complementary modularity resolution was particularly useful for annotating rare cell-types while maintaining low noise levels (Notebook 4).

### AZ pseudo-bulk DEG analysis comparing WT and *hae hsl2*

We performed pseudo-bulking for the AZ (Seurat cluster 11) for each WT and *hae hsl2* sample, yielding four samples for each genotype (Notebook 5). We then used edgeR to normalize and calculate CPM and LCPM matrices. We modeled whether the cells were sorted as a nuisance factor (to account for variation due to FACS), and otherwise only included genotype as a factor (WT or mutant). We then constructed a contrast testing the hypothesis that the genotype factor for the difference in expression between WT and *hae hsl2* for each gene was equal to 0. The output of edgeR has been included as Supplemental Table 7. A file with our edgeR functions has been included in Supplemental Code. This code requires an Arabidopsis annotation file derived from TAIR10 included as Supplemental Table 15.

### GO term analysis

For GO analysis we used shinyGO v 0.76^67^. For all analyses we displayed Biological Process terms at FDR < .01. For GO analysis of WT AZs we used the output of the Seurat FindAllMarkers function to select genes log2(Fold Enrichment) > 1 with FDR < .05 in the AZ. We used all genes from the FindAllMarkers analysis as a universe (i.e., genes expressed in at least one cluster). For the WT-mutant DE GO analysis, we took genes defined as DE log2(FC) > 1 with FDR < .05 with all genes expressed in at least 3 samples as universe. GO analysis of the bulk DE/single-cell DE intersection was performed with a universe as the genes defined to be expressed in the WT/*hae hsl2* pseudo-bulk DEG analysis.

### AZ subclustering

AZ cells (those from cluster 11) were sub-divided into those of WT and *hae hsl2* origin, and each dataset was reintegrated using a similar process as above (Notebook 6). We then performed coarse clustering with resolution = .1 in Seurat which identified two transcriptionally distinct groups for both genotypes. We performed pseudo-bulk Spearman correlation analysis comparing expression of each of the two clusters in each genotype to previously published FACS secession and residuum datasets^15^. For enrichment analysis we took the pseudo-bulked putative residuum cells and secession cells and performed edgeR analysis as above, except we pooled the cells derived from the two FACS samples for both mutant and WT due to a particularly low number of cells in WT sorted sample #1. The secession and residuum associated genes are those with FDR < .05 and log2(fold enrichment) > 1 in the respective cell types. We will note because the cells were clustered first before running the edgeR analysis, the resulting gene lists were not unbiased estimates of differential expression. However, because we performed identical procedures for WT and mutant, our Fisher’s exact test is valid because it is testing the hypothesis that the proportion of genes in this “enriched” set is different between the two genotypes.

### Bulk RNA-Seq library generation and sequencing

For the sorted *HAE-YFP* bulk samples, we sorted into 20 μl of RNA-Later then used the magnetic bead based Direct-zol-96 MagBead RNA-Isolation kit (Zymo) to isolate RNA. For the protoplast test, we cut three receptacles per replicate and digested as above. We harvested the tissue by spinning at 500 x g and removing supernatant, leaving digested cells and undigested tissue in place, before freezing in liquid N2. We simultaneously collected three receptacles per replicate in our unprotoplasted control where tissue was placed in 10 μl of RNA-Later in the cap of a 1.7 ml microcentrifuge tube, tapped to the bottom of the tube, immediately frozen in liquid N2, homogenized with a blue pestle, and performed RNA isolation as above. We performed three replicates in each condition.

For the *fal-3/fal-7* bulk RNA-Seq experiment, we isolated receptacles from three stage 15 flowers for each replicate of *er gl* (WT grandparent), *er gl hae-3 hsl2-3* (mutant parent), and *er gl hae-3 hsl2-3 fal-3/fal-7* (suppressors). Tissue was placed into the cap of a 1.7 ml tube containing 10 μl RNA-Later. After tapping the tissue to the bottom of the tube, we froze in liquid nitrogen. We then ground the tissue in liquid N2 using blue pestles, and used the Zymo RNA-isolation kit as before. For all samples, RNA integrity was checked with Bioanalyzer RNA Nano kit and quantified by Qubit.

For library generation, we used Lexogen Quantseq RNA-Seq using the manufacturer’s protocol, with instructions for “Low input” for the sorted samples due to input of only 5-10 ng total RNA input per sample. We used the Unique Molecular Identifier (UMI) PCR add-on kit (Lexogen). Libraries were indexed and sequenced on an Illumina NextSeq, High Output setting. Reads were aligned to the TAIR10 genome using the STAR aligner, deduplicated using UMI-Tools, and counted with HTSeq-Count. Counts were analyzed with edgeR. We defined “expressed genes” to be those with observed reads in three or more libraries. Raw reads have been deposited at the Sequence Read Archive under BioProject PRJNA857332.

### PAGE analysis of abscission-associated gene expression in *fal* mutants

We constructed an expression matrix for the 67 bulk/single-cell intersection gene set for WT, *hae hsl2, hae hsl2 fal-3,* and *hae hsl2 fal-3* using the log2(CPM) values generated by edgeR (Notebooks 7 and 8). We then summed the log2(CPM) values of each gene for each sample and normalized to the average summed values of the *hae hsl2* samples before dividing by the number of genes in the analysis. The resulting quantity represents the average log2(FC) for each sample compared to the *hae hsl2* average. Last, we performed pairwise T-tests with Bonferroni Correction assuming equal variance.

### Breakstrength measurements

The breakstrength of petals of stage 16 flowers (i.e., silique 1-2 mm above petals) in *er gl hae-3 hsl2-3* and *er gl hae-3 hsl2-3 fal-3/7* were measured using our previously described petal break strength meter and analysis script^45^. In brief, the petals were clamped to the meter and the flower pulled down with forceps until the petal detached. The maximum voltage was extracted from the output file of the meter. This voltage reading was converted to an equivalent force after calculation of a standard curve based on voltage readings of the meter attached to a varying number of objects of known weight (i.e., paper clips). For measurements taken at each temperature, we performed pairwise T-tests with Bonferroni Correction assuming equal variance.

### *fal* mutant identification

The *hae-3 hsl2-3* suppressor screen was performed as previously described^26^. For identification, we used TAIL-PCR^68^ to amplify a PCR fragment in *fal-7* which was analyzed by Sanger sequencing. For *fal-3,* we used TAIL-PCR to identify an insertion in an exon of AT2G07690, which is a member of the Minichromosome Maintenance gene family involved in initiation of DNA replication. However, given the similar phenotype of *fal-7* we hypothesized there may exist an additional mutation in *MKP1*. We designed Sanger sequencing primers and tiled the coding sequence, detecting a G-> A SNP at bp 1357 in the *MKP1* coding sequence using primers listed in Supplemental Table 13. We performed linkage analysis in the *hae hsl2 x hae hsl2 fal-7* F2 population using genotyping primers listed in Supplemental Table 13.

## Conflict of Interest

The authors declare they have no conflict of interest.

## Funding

This work was supported by NSF MCB0743955 to JCW, NIH 1R35GM131725 to PNB, who is also supported by the Howard Hughes Medical Institute as an investigator, USDA 2021-67034-35139 to IWT, and Deutsche Forschungsgemeinschaft (DFG) 2403 to UO and CWH.

## Acknowledgements

Figure 5I was created with BioRender.

---

**Supplemental Figure 1:**
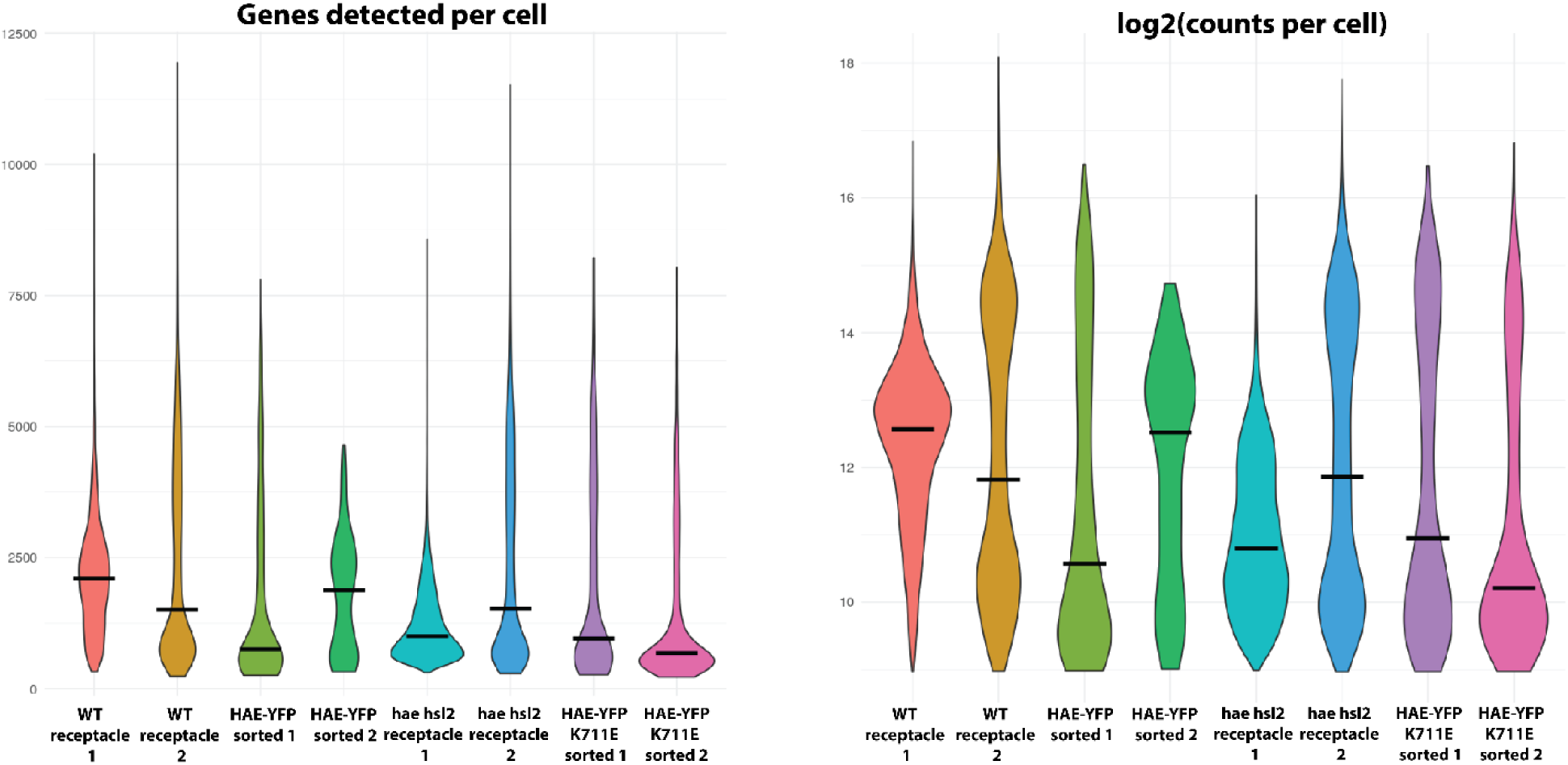
Plots of the number of genes detected per cell and log2(counts per cell)

**Supplemental Figure 2:**
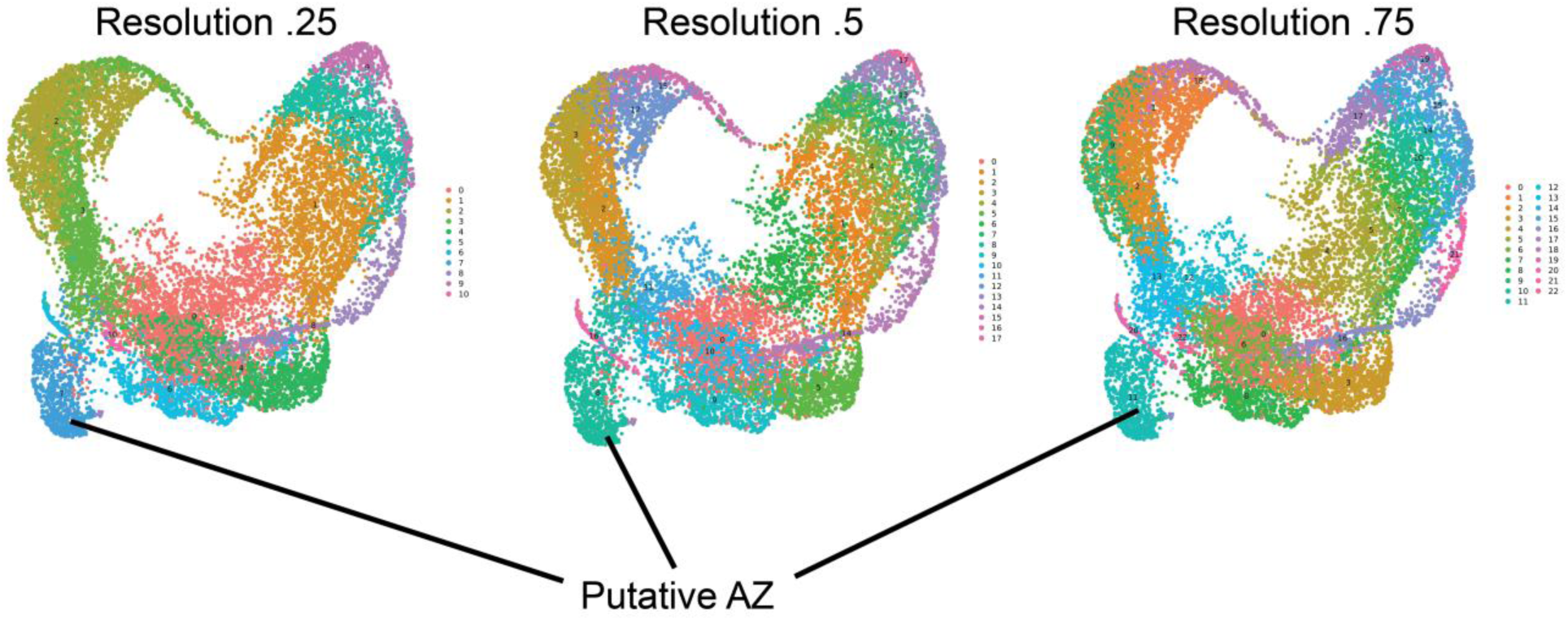
The putative AZ cluster is similar across a range of clustering resolutions.

**Supplemental Figure 3:**
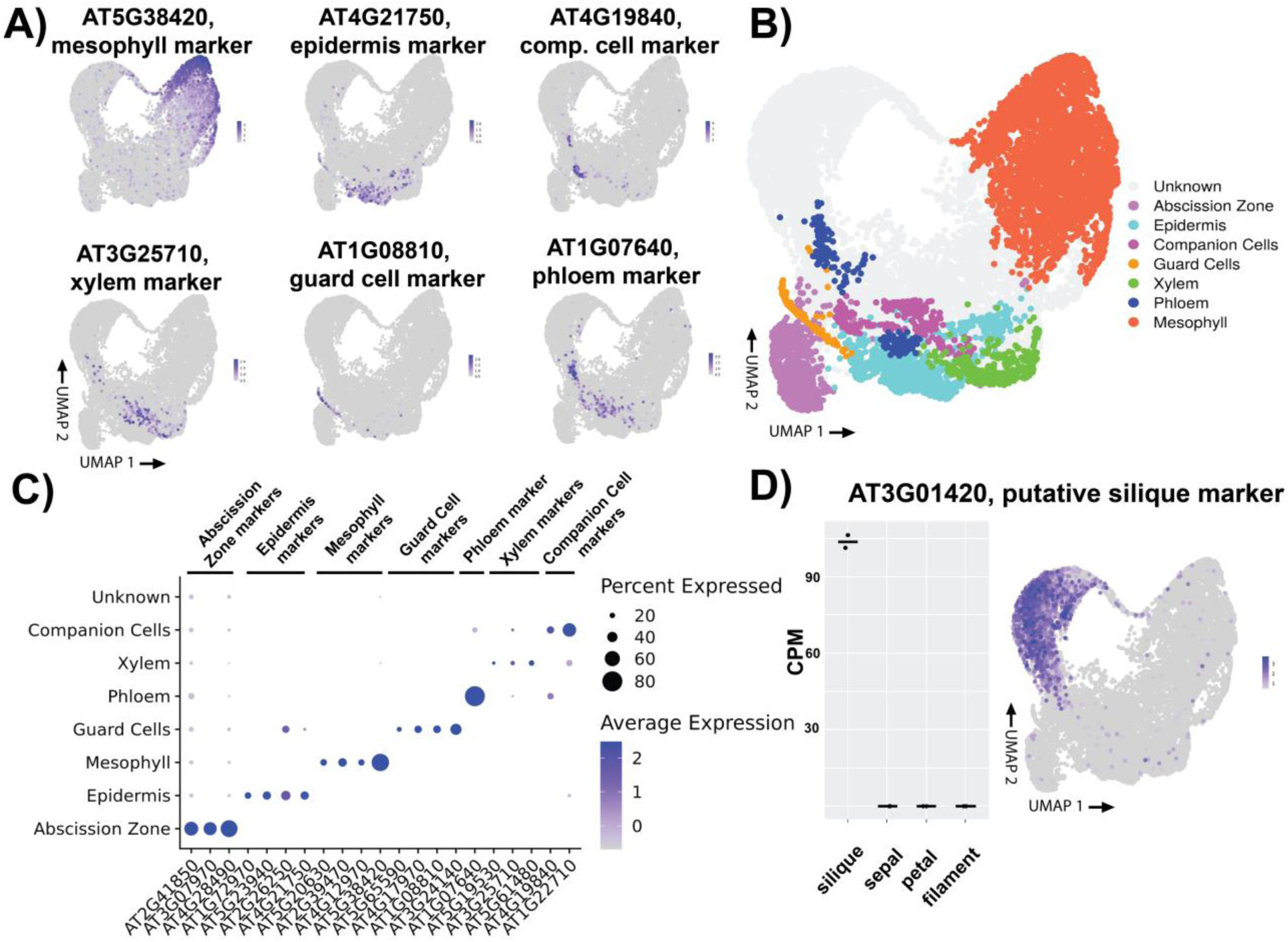
Identification of additional cell types. A) Plotting cell type markers identified from a prior single-cell study of Arabidopsis leaves. B) Tentative cell-type identification based on expression of known marker cell-type marker genes. C) Distribution of marker gene expression across putative cell types. D) Expression of putative silique marker gene in previously published bulk data (left panel), and expression of the same putative silique marker gene from our single-cell data (right panel).

**Supplemental Figure 4:**
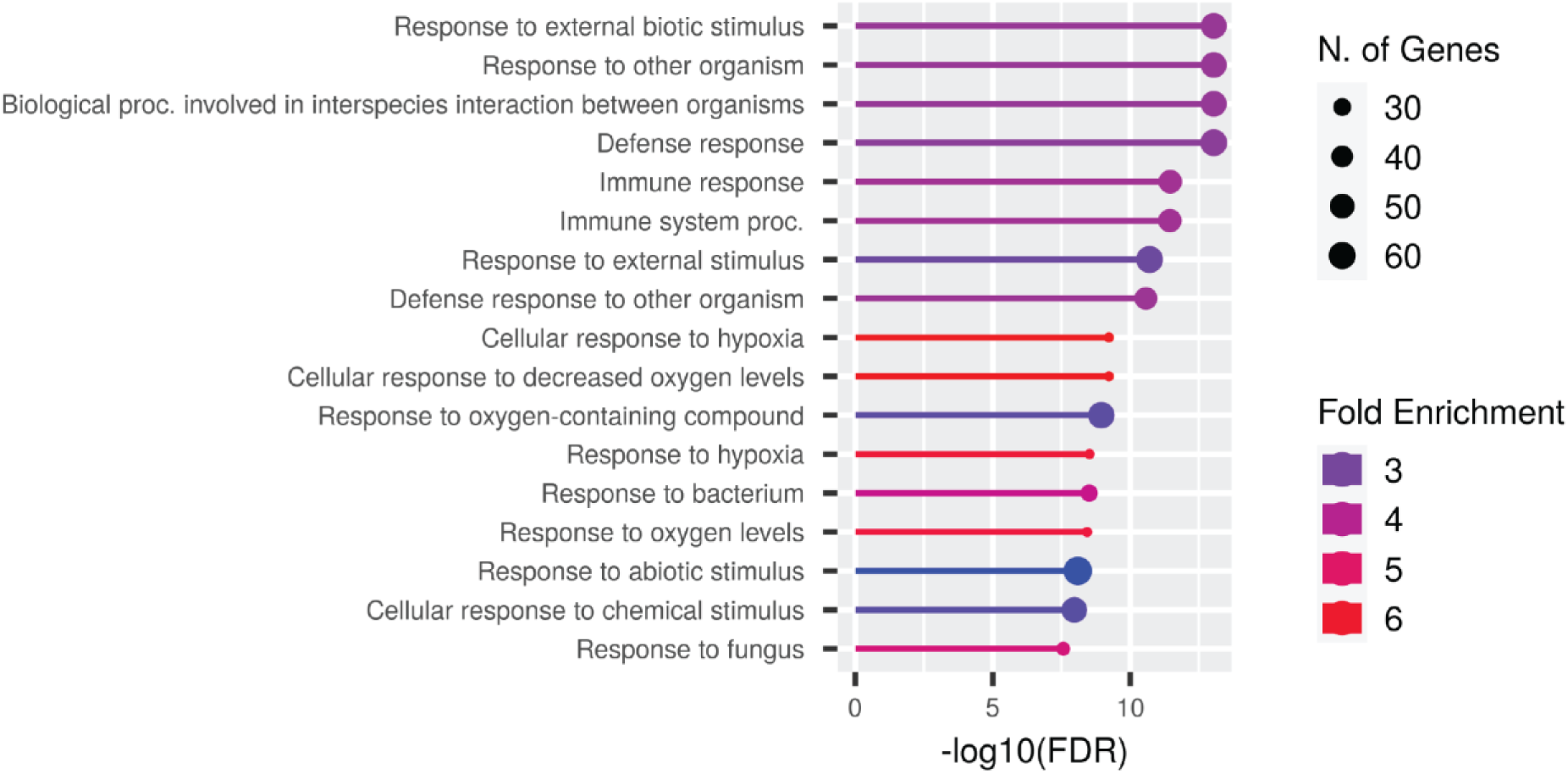
GO term enrichment analysis of genes higher in *hae hsl2* compared to WT AZs.

**Supplemental Figure 5:**
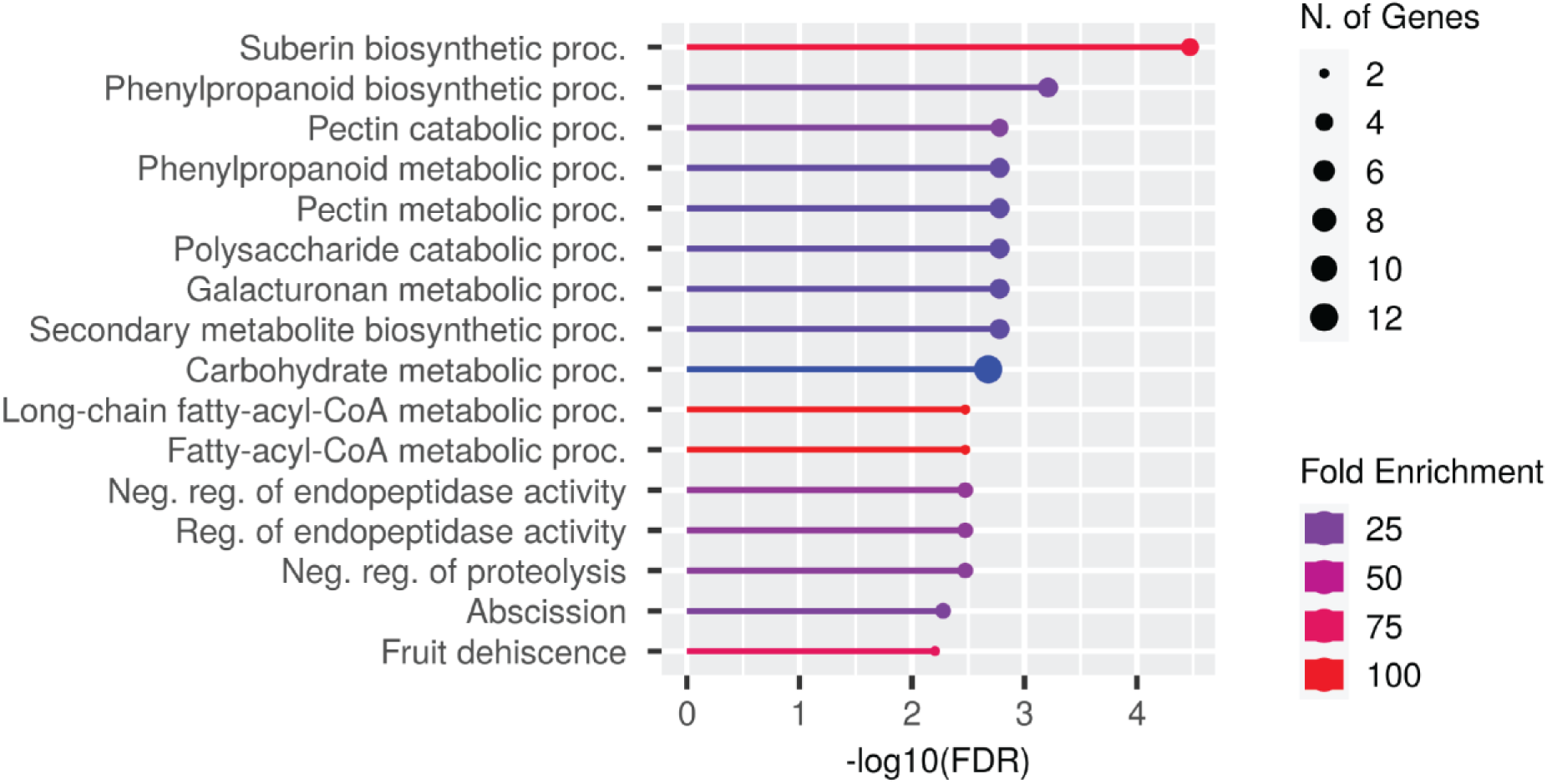
GO term enrichment analysis of genes lower in *hae hsl2* compared to WT from both bulk and single-cell RNA-Seq.

**Supplemental Figure 6:**
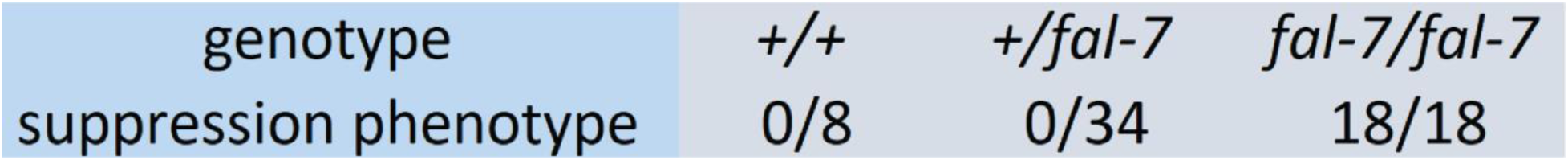
Association of phenotype and genotype in an F2 *hae hsl2 x hae hsl2 fal-7* backcross population.

**Supplemental Figure 7:**
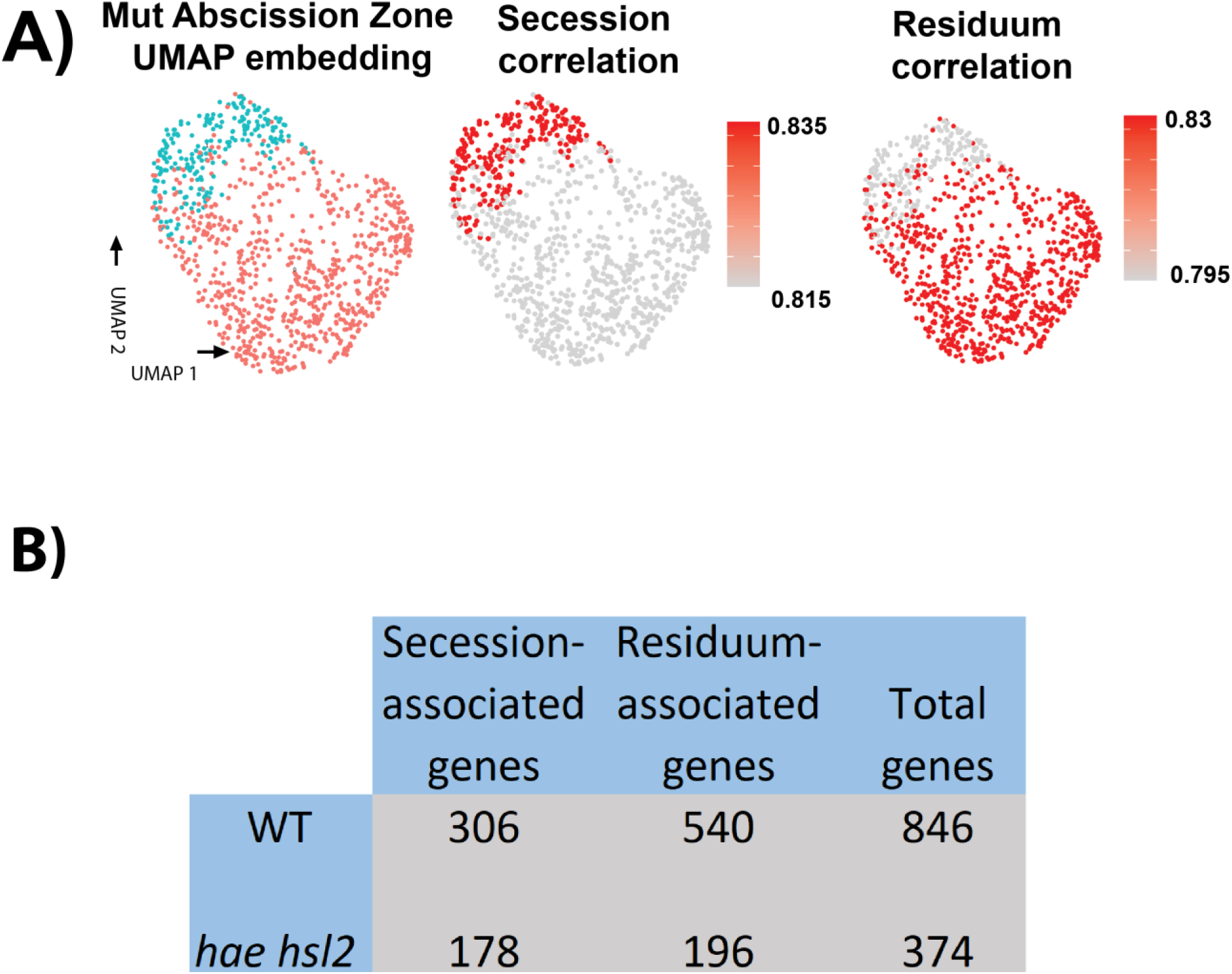
Sub-clustering and differential enrichment analysis of mutant AZ cells. A) UMAP and coloration of low-resolution mutant AZ cell clusters (left), along with correlation to bulk secession and residuum RNA-Seq data (right) B) Table of differentially enriched genes in secession and residuum cells across both WT and mutant samples

**Supplemental Figure 8:**
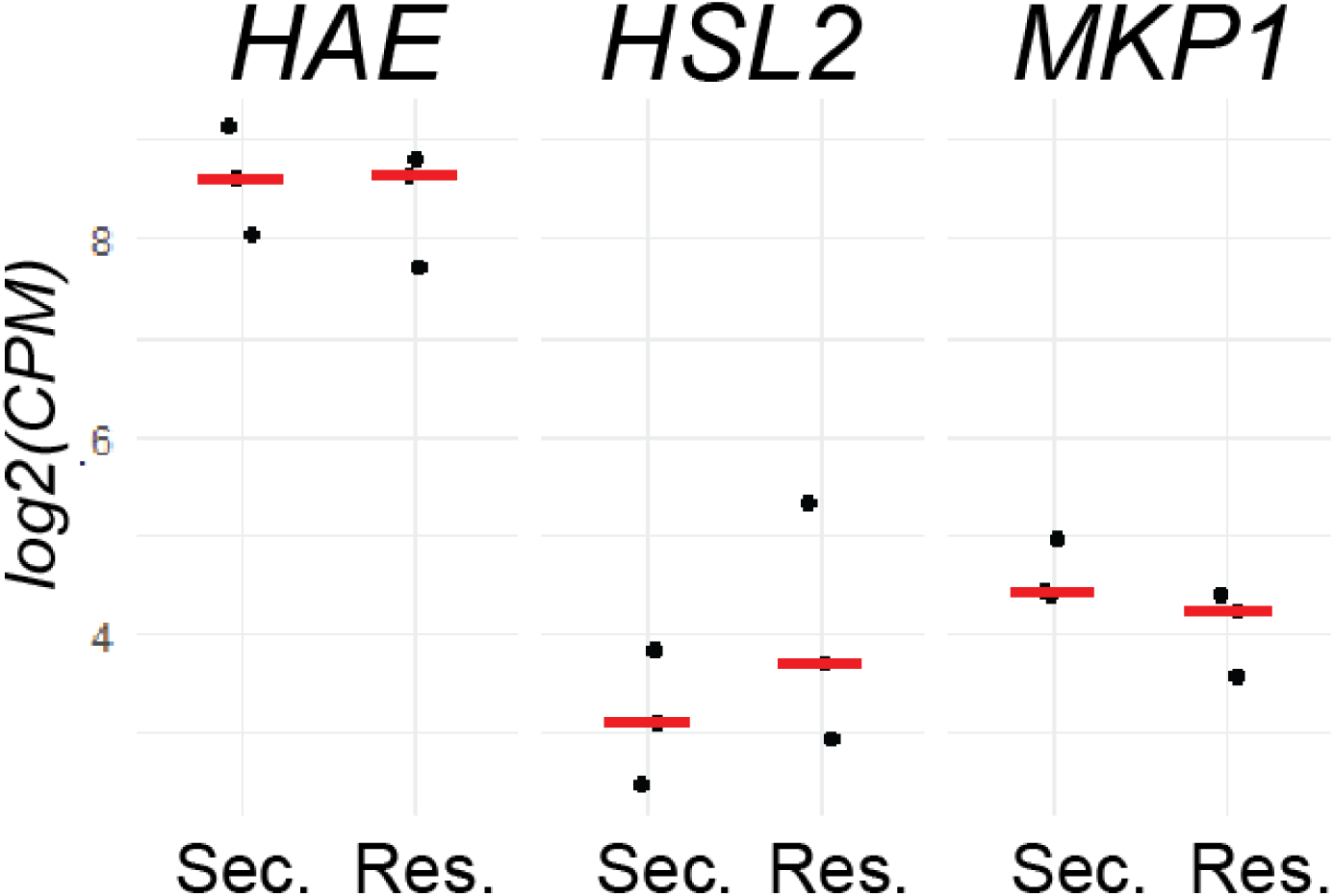
Pseudobulk log2(CPM) of *HAE, HSL2*, and *MKP1* in putative secession and residuum cells.

## Notes

https://www.ncbi.nlm.nih.gov/sra

### Competing Interest Statement

The authors have declared no competing interest.

### Summary of Updates

We have performed additional experimentation to define the expression pattern of the IDA gene and to validate the functional relevance of this expression pattern.

