## Supplemental Code for "Arabidopsis uses a molecular grounding mechanism and a biophysical circuit breaker to limit floral abscission signaling": edgeR_function.R

```
counts_to_reads_df <- function(path_to_counts) {

    #get all files in bulk directory and load them in
    file_list <- list.files(path=path_to_counts, recursive = T)
    reads <- list()
    targets_rows = data.frame()

    for (i in 1:length(file_list)){
      reads[[i]]=read.table(paste(path_to_counts, file_list[i], sep = "/"), sep="\t")
      rownames(reads[[i]]) =reads[[i]]$V1
      if (i == 1) {
        target_rows=reads[[i]][,1]
      }
      target_rows = intersect(target_rows, reads[[i]][,1])
      print(file_list[[i]])
    }

    #make df
    reads_df <- data.frame(matrix(ncol=0, nrow=length(target_rows)))

    #loop over all read objects and put in column of df
    for (i in 1:length(reads)){
      reads_df[,i]=reads[[i]][target_rows,2]
    }

    rownames(reads_df) = target_rows
    return(reads_df)
}

edgeR_2_sample <- function(reads_df, s1_label, s2_label, s1_indices, s2_indices, annotations, design, my.contrasts) {
  
  
  #make DGElist
  x3 <- DGEList(counts = reads_df, genes = rownames(reads_df))
  
  
  #reads per library uniquely mapped to a gene:
  
  #make cpm and lcpm
  cpm <- cpm(x3)
  lcpm <- cpm(x3, log=TRUE)
  
  #keep only genes that are expressed. "Expressed" here means counts observed in at least 3 samples
  dim(x3)
  keep.exprs <- rowSums(cpm>0)>=3
  x3 <- x3[keep.exprs,, keep.lib.sizes=FALSE]
  dim(x3) #compare to dim(x) above
  
  #normalize data after removing low expressed genes
  x3 <- calcNormFactors(x3)
  
  #cpm, lcpm of normalized values
  cpm <- cpm(x3)
  lcpm <- cpm(x3, log=TRUE)
  
   #fit normalized dge list to model matrix
  x3 <- estimateDisp(x3, design)
  fit <- glmQLFit(x3, design)

  #tr <- glmTreat(fit, coef=2, lfc=1)
  #topTags(tr)
  
  #QLF tests 
  s1_v_s2 <- glmQLFTest(fit, contrast=my.contrasts[,"s1_v_s2"]) 
 
   
  #extract results. toptags adds FDR column.
  sec2 <- data.frame(topTags(s1_v_s2, n = "Inf")$table)
  #sec2=sec2[c(1,7,2,3,4,5,6)]
  sec2[[paste(s1_label,"av", sep="_")]] <- apply(cpm[sec2[["genes"]],s1_indices], 1,mean)
  sec2[[paste(s2_label,"av", sep="_")]] <- apply(cpm[sec2[["genes"]],s2_indices], 1,mean)
  
  #return(sec2)
  sec2 = cbind(sec2, cpm[sec2$genes,c(s1_indices, s2_indices)])
  #return(sec2)
  sec2=merge(sec2, annotations, by.x=c("genes"), by.y=c("gene_id"), all.x = T, all.y = F)
  sec2= sec2[!duplicated(sec2$genes), ]
  sample_names = paste(c(rep(s1_label, length(s1_indices)), rep(s2_label, length(s2_indices))), c(1:length(s1_indices), 1:length(s2_indices)), sep = "_")
  colnames(sec2)[c(9:(8+length(s1_indices) + length(s2_indices)))] <- sample_names
  #rownames(sec2)=sec2$genes
  return(sec2)

}
```
