## Supplemental Code for "Arabidopsis uses a molecular grounding mechanism and a biophysical circuit breaker to limit floral abscission signaling": notebook_01_sc_Ko_to_Seurat_object_3_8_21.pdf

### WB01_sc_KB_to_Seurat_object_3_8_21

June 30, 2022

[1]:
```
#THIS SCRIPT WILL PERFORM EMPTYDROPS ANALYSIS, SEURAT OBJECT CREATION, AND␣
↪MULTIPLET IDENTIFICATION

suppressMessages(library(BUSpaRse))
suppressMessages(library(Matrix))
suppressMessages(library(tidyverse))
suppressMessages(library(Seurat))
suppressMessages(library(DropletUtils))
suppressMessages(library(DoubletFinder))

proto_genes=read.csv("../data/bulk_data/protoplasting.csv")
proto_list=as.character(proto_genes[abs(proto_genes$logFC) > 2,]$genes)

# Slightly modified from BUSpaRse, just to avoid installing a few dependencies␣
↪not used here
read_count_output <- function(dir, name) {
  dir <- normalizePath(dir, mustWork = TRUE)
  m <- readMM(paste0(dir, "/", name, ".mtx"))
  m <- Matrix::t(m)
  m <- as(m, "dgCMatrix")
  # The matrix read has cells in rows
  ge <- ".genes.txt"
  genes <- readLines(file(paste0(dir, "/", name, ge)))
  barcodes <- readLines(file(paste0(dir, "/", name, ".barcodes.txt")))
  colnames(m) <- barcodes
  rownames(m) <- genes
  return(m)
}
```

[2]:
```
sessionInfo()
```

```
R version 3.6.3 (2020-02-29)
Platform: x86_64-conda_cos6-linux-gnu (64-bit)
Running under: Ubuntu 20.04.2 LTS

Matrix products: default
BLAS/LAPACK: /home/robotmessenger810/anaconda3/envs/r_3/lib/libopenblasp-r0.3.9.so
```

```
locale:
 [1] LC_CTYPE=en_US.UTF-8       LC_NUMERIC=C
 [3] LC_TIME=en_US.UTF-8        LC_COLLATE=en_US.UTF-8
 [5] LC_MONETARY=en_US.UTF-8    LC_MESSAGES=en_US.UTF-8
 [7] LC_PAPER=en_US.UTF-8       LC_NAME=C
 [9] LC_ADDRESS=C               LC_TELEPHONE=C
[11] LC_MEASUREMENT=en_US.UTF-8 LC_IDENTIFICATION=C

attached base packages:
[1] parallel  stats4    stats     graphics  grDevices utils     datasets
[8] methods   base

other attached packages:
 [1] DoubletFinder_2.0.3         DropletUtils_1.6.1
 [3] SingleCellExperiment_1.8.0  SummarizedExperiment_1.16.1
 [5] DelayedArray_0.12.3         BiocParallel_1.20.1
 [7] matrixStats_0.61.0          Biobase_2.46.0
 [9] GenomicRanges_1.38.0        GenomeInfoDb_1.22.1
[11] IRanges_2.20.2              S4Vectors_0.24.4
[13] BiocGenerics_0.32.0         Seurat_3.1.5
[15] forcats_0.5.0               stringr_1.4.0
[17] dplyr_1.0.7                 purrr_0.3.4
[19] readr_1.4.0                 tidyr_1.1.3
[21] tibble_3.1.6                ggplot2_3.3.5
[23] tidyverse_1.3.0             Matrix_1.2-18
[25] BUSpaRse_1.0.0

loaded via a namespace (and not attached):
  [1] readxl_1.3.1            uuid_0.1-4              backports_1.4.1
  [4] BiocFileCache_1.10.2    plyr_1.8.6              igraph_1.2.5
  [7] repr_1.1.0              lazyeval_0.2.2          splines_3.6.3
 [10] listenv_0.8.0           digest_0.6.29           ensembldb_2.10.2
 [13] htmltools_0.5.0         fansi_0.5.0             magrittr_2.0.1
 [16] memoise_1.1.0           BSgenome_1.54.0         cluster_2.1.2
 [19] ROCR_1.0-11             limma_3.42.2            globals_0.12.5
 [22] Biostrings_2.54.0       modelr_0.1.6            RcppParallel_5.1.5
 [25] R.utils_2.9.2           askpass_1.1             prettyunits_1.1.1
 [28] colorspace_2.0-2        blob_1.2.1              rvest_0.3.5
 [31] rappdirs_0.3.3          ggrepel_0.9.1           haven_2.3.1
 [34] crayon_1.4.2            RCurl_1.98-1.2          jsonlite_1.7.2
 [37] zeallot_0.1.0           survival_3.1-12         zoo_1.8-8
 [40] ape_5.3                 glue_1.6.0              gtable_0.3.0
 [43] zlibbioc_1.32.0         XVector_0.26.0          leiden_0.3.9
 [46] plyranges_1.6.10        Rhdf5lib_1.8.0          future.apply_1.6.0
 [49] HDF5Array_1.14.4        scales_1.1.1            edgeR_3.28.1
 [52] DBI_1.1.2               Rcpp_1.0.7              viridisLite_0.4.0
 [55] progress_1.2.2          dqrng_0.3.0             reticulate_1.16
 [58] rsvd_1.0.3              bit_4.0.4               tsne_0.1-3
```

```
 [61] htmlwidgets_1.5.1       httr_1.4.2              RColorBrewer_1.1-2
 [64] ellipsis_0.3.2          ica_1.0-2               R.methodsS3_1.8.1
 [67] pkgconfig_2.0.3         XML_3.99-0.3            uwot_0.1.8
 [70] dbplyr_1.4.2            locfit_1.5-9.4          utf8_1.2.2
 [73] reshape2_1.4.4          tidyselect_1.1.1        rlang_0.4.12
 [76] AnnotationDbi_1.48.0    munsell_0.5.0           cellranger_1.1.0
 [79] tools_3.6.3             cli_3.1.0               generics_0.1.1
 [82] RSQLite_2.2.0           broom_0.5.5             ggridges_0.5.2
 [85] evaluate_0.14           bit64_0.9-7             fs_1.5.2
 [88] fitdistrplus_1.1-1      RANN_2.6.1              AnnotationFilter_1.10.0
 [91] pbapply_1.5-0           future_1.18.0           nlme_3.1-147
 [94] R.oo_1.23.0             xml2_1.3.1              biomaRt_2.42.1
 [97] compiler_3.6.3          rstudioapi_0.13         png_0.1-7
[100] plotly_4.9.2.1          curl_4.3                reprex_0.3.0
[103] stringi_1.7.6           GenomicFeatures_1.38.2  lattice_0.20-45
[106] IRdisplay_0.7.0         ProtGenerics_1.18.0     vctrs_0.3.8
[109] pillar_1.6.4            lifecycle_1.0.1         lmtest_0.9-38
[112] RcppAnnoy_0.0.19        data.table_1.14.2       cowplot_1.1.1
[115] bitops_1.0-7            irlba_2.3.5             patchwork_1.1.1
[118] rtracklayer_1.46.0      R6_2.5.1                gridExtra_2.3
[121] KernSmooth_2.23-20      codetools_0.2-18        MASS_7.3-54
[124] assertthat_0.2.1        rhdf5_2.30.1            openssl_1.4.6
[127] withr_2.4.3             sctransform_0.2.1       GenomicAlignments_1.22.1
[130] Rsamtools_2.2.3         GenomeInfoDbData_1.2.2  hms_1.1.0
[133] grid_3.6.3              IRkernel_1.1            Rtsne_0.15
[136] pbdZMQ_0.3-6            lubridate_1.7.8         base64enc_0.1-3
```

[2]:
```
files <- list.files(path="../data/scKB_outs/", full.names=TRUE, recursive=FALSE)
files_base = list.files(path="../data/scKB_outs/", full.names=FALSE,␣
↪recursive=FALSE)
```

[ ]:
```
#prevent warnings from printing
defaultW <- getOption("warn")
options(warn = -1)

seu_list = list()

#loop through all files and perform empty drops quantification and make seurat␣
↪objects
for (i in 1:length(files)){
    sample = files[i]

    #read in spliced matrix and retrain only Arabidopsis gene counts (important␣
↪for species mixing experiments)
    spliced = read_count_output(sample, "spliced")
```

```r
  spliced = spliced[grepl("AT",unlist(spliced@Dimnames[1]), fixed=TRUE),,␣
↪drop=FALSE]

  #read in unspliced matrix and retrain only Arabidopsis gene counts
  unspliced = read_count_output(sample, "unspliced")
  unspliced = unspliced[grepl("AT",unlist(unspliced@Dimnames[1]),␣
↪fixed=TRUE),, drop=FALSE]

  #Find barcodes identified in both spliced and unspliced count matrices
  shared = intersect(colnames(spliced), colnames(unspliced))

  #filter to barcodes present in both spliced and unspliced matrices
  spliced = spliced[,shared]
  unspliced = unspliced[,shared]

  combined = spliced + unspliced

  #run empty drops on combined count matrix
  empty_drops = emptyDrops(combined[grepl(pattern = "AT[1-5]",␣
↪unlist(spliced@Dimnames[1])),, drop=FALSE], ignore = 500, lower = 300)

  #We take all cells from spliced and unspliced that were called by␣
↪emptydrops, and then we sum them into combined matrix
  shared = intersect(intersect(colnames(spliced), colnames(unspliced)),␣
↪rownames(empty_drops[!is.na(empty_drops$FDR) & empty_drops$FDR < .001,]))

  spliced = spliced[,shared]
  unspliced = unspliced[,shared]

  combined = spliced + unspliced

  #create seurat object
  seu_obj <- CreateSeuratObject(combined, min.cells = 3)

  #add spliced and unspliced as assays
  spliced_assay <- CreateAssayObject(counts = spliced)
  unspliced_assay <- CreateAssayObject(counts = unspliced)

  seu_obj[["spliced"]] = spliced_assay
  seu_obj[["unspliced"]] = unspliced_assay

  #mito and plastid read percent
  seu_obj = PercentageFeatureSet(seu_obj, pattern = "ATM", col.name =␣
↪"percent.mito", assay = "RNA")
  seu_obj = PercentageFeatureSet(seu_obj, pattern = "ATC", col.name =␣
↪"percent.cp", assay = "RNA")
```

```
  #doublet finder
  df_seu <- NormalizeData(seu_obj)
  df_seu <- FindVariableFeatures(df_seu, selection.method = "vst", nfeatures␣
↪= 2000)
  df_seu <- ScaleData(df_seu)
  df_seu <- RunPCA(df_seu)
  df_seu <- RunTSNE(df_seu, dims = 1:15)

  #simple approximate expected doublet rate based on equation from 10x data:␣
↪doublet_percent = .004/500 * #_cells
  nExp_poi <- round((0.004 /500*(nrow(df_seu@meta.data)) * nrow(df_seu@meta.
↪data))


  sweep.sweep.df_seu <- paramSweep_v3(df_seu, PCs = 1:15, sct = FALSE)
  sweep.stats_df_seu <- summarizeSweep(sweep.sweep.df_seu, GT = FALSE)
  bcmvn_sweep.df_seu <- find.pK(sweep.stats_df_seu)

  pK = double(bcmvn_sweep.df_seu[max(bcmvn_sweep.
↪df_seu$BCmetric)==bcmvn_sweep.df_seu$BCmetric,2])
  df_seu <- doubletFinder_v3(df_seu, PCs = 1:15, pN = 0.25, pK = pK, nExp =␣
↪nExp_poi, reuse.pANN = FALSE, sct = FALSE)

  seu_obj <- subset(seu_obj, subset = (percent.mito < 10) & df_seu@meta.
↪data[,dim(df_seu@meta.data)[2]] =="Singlet")

  #set original experiment
  seu_obj@meta.data$orig.ident = files_base[i]

  #set genotype
  if (files_base[i] %in% c("sc_101", "sc_103", "sc_26_combined", "sc_67",␣
↪"sc_69")) {
      seu_obj@meta.data$geno = "WT"
  }
  else {
      seu_obj@meta.data$geno = "mutant"
  }

  #set experiment
  if (files_base[i] %in% c("sc_101", "sc_102", "sc_103", "sc_104", "sc_69",␣
↪"sc_70")) {
      seu_obj@meta.data$experiment = "sorted"
  }
```

```
    else {
        seu_obj@meta.data$experiment = "nonsorted"
    }


    print(sample)
    print("original # cells: ")
    print(nrow(df_seu@meta.data))
    print("singlet # cells: ")
    print(nrow(seu_obj@meta.data))

    saveRDS(seu_obj, file = paste("../data/seurat_objects/seurat_raw_1_4_22/",␣
 ↪files_base[i], ".rds", sep=""))
}
```
