## Supplemental Code for "Arabidopsis uses a molecular grounding mechanism and a biophysical circuit breaker to limit floral abscission signaling": notebook_02_nT_mut_integration.pdf

### WB02_WT_mut_integration

June 30, 2022

[1]:
```
#THIS SCRIPT PERFORMS SCTRANSFORM AND THEN INTEGRATES THE WT AND MUTANT OBJECTS␣
↪INTO A SINGLE SEURAT OBJECT

suppressMessages(library(Seurat))
library(ggplot2)

proto_genes=read.csv("../data/bulk_data/protoplasting.csv")
proto_list=as.character(proto_genes[abs(proto_genes$logFC) > 1,]$genes)

#FUNCTION USED IN THIS SCRIPT
#takes a list of Seurat objects with SCTransform already run
seu_integrate <- function(..., filename, nfeatures){
    seu.list <- list(...) # THIS WILL BE A LIST STORING EVERYTHING:

    ref.genes = rownames(seu.list[[1]]@assays$RNA)
    assay_list <- rep("SCT", length(seu.list))


    # integration
    rc.features <- SelectIntegrationFeatures(object.list = seu.list, nfeatures␣
↪= nfeatures)

    #remove genes influenced by protoplasting to a high degree as well as␣
↪plastid/mitochodrial genes
    rc.features <- rc.features[(!c(grepl("ATMG",rc.features) | grepl("ATCG",rc.
↪features) | rc.features%in%proto_list))]

    seu.list <- PrepSCTIntegration(object.list = seu.list, anchor.features = rc.
↪features, verbose = TRUE, assay = assay_list)
    seu.list <- lapply(X = seu.list, FUN = RunPCA, verbose = FALSE, features =␣
↪rc.features)
    rc.anchors <- FindIntegrationAnchors(object.list = seu.list, normalization.
↪method = "SCT", anchor.features = rc.features, verbose = TRUE, reference=1,␣
↪reduction = "rpca")
    to_integrate <- Reduce(intersect, lapply(rc.anchors@object.list, rownames))

    #integrate data and keep full geneset
```

```
    rc.integrated <- IntegrateData(anchorset = rc.anchors,  features.to.
↪integrate = to_integrate, normalization.method = "SCT", verbose = TRUE)
    rc.integrated <- RunPCA(rc.integrated, npcs = 50, verbose = FALSE, approx =␣
↪FALSE)

    #save object
    saveRDS(rc.integrated, file = paste("../data/intd_seu_objects/",filename,".
↪rds", sep = ""))
    return(rc.integrated)
#    }
}
```

[2]:
```
sessionInfo()
```

```
R version 3.6.3 (2020-02-29)
Platform: x86_64-conda_cos6-linux-gnu (64-bit)
Running under: Ubuntu 20.04.2 LTS

Matrix products: default
BLAS/LAPACK: /home/robotmessenger810/anaconda3/envs/r_3/lib/libopenblasp-r0.3.9.so

locale:
 [1] LC_CTYPE=en_US.UTF-8       LC_NUMERIC=C
 [3] LC_TIME=en_US.UTF-8        LC_COLLATE=en_US.UTF-8
 [5] LC_MONETARY=en_US.UTF-8    LC_MESSAGES=en_US.UTF-8
 [7] LC_PAPER=en_US.UTF-8       LC_NAME=C
 [9] LC_ADDRESS=C               LC_TELEPHONE=C
[11] LC_MEASUREMENT=en_US.UTF-8 LC_IDENTIFICATION=C

attached base packages:
[1] stats     graphics  grDevices utils     datasets  methods   base

other attached packages:
[1] ggplot2_3.3.5 Seurat_3.1.5

loaded via a namespace (and not attached):
 [1] httr_1.4.2         tidyr_1.1.3        jsonlite_1.7.2     viridisLite_0.4.0
 [5] splines_3.6.3      leiden_0.3.9       ggrepel_0.9.1      globals_0.12.5
 [9] pillar_1.6.4       lattice_0.20-45    glue_1.6.0         reticulate_1.16
[13] uuid_0.1-4         digest_0.6.29      RColorBrewer_1.1-2 colorspace_2.0-2
[17] cowplot_1.1.1      htmltools_0.5.0    Matrix_1.2-18      plyr_1.8.6
[21] pkgconfig_2.0.3    tsne_0.1-3         listenv_0.8.0      purrr_0.3.4
[25] patchwork_1.1.1    scales_1.1.1       RANN_2.6.1         Rtsne_0.15
[29] tibble_3.1.6       generics_0.1.1     ellipsis_0.3.2     withr_2.4.3
[33] repr_1.1.0         ROCR_1.0-11        pbapply_1.5-0      lazyeval_0.2.2
[37] survival_3.1-12    magrittr_2.0.1     crayon_1.4.2       evaluate_0.14
[41] future_1.18.0      fansi_0.5.0        nlme_3.1-147       MASS_7.3-54
```

```
[45] ica_1.0-2          tools_3.6.3        fitdistrplus_1.1-1 data.table_1.14.2
[49] lifecycle_1.0.1    stringr_1.4.0      plotly_4.9.2.1     munsell_0.5.0
[53] cluster_2.1.2      irlba_2.3.5        compiler_3.6.3     rsvd_1.0.3
[57] rlang_0.4.12       grid_3.6.3         ggridges_0.5.2     pbdZMQ_0.3-6
[61] IRkernel_1.1       RcppAnnoy_0.0.19   htmlwidgets_1.5.1  igraph_1.2.5
[65] base64enc_0.1-3    gtable_0.3.0       codetools_0.2-18   DBI_1.1.2
[69] reshape2_1.4.4     R6_2.5.1           gridExtra_2.3      zoo_1.8-8
[73] dplyr_1.0.7        uwot_0.1.8         future.apply_1.6.0 utf8_1.2.2
[77] KernSmooth_2.23-20 ape_5.3            stringi_1.7.6      parallel_3.6.3
[81] IRdisplay_0.7.0    Rcpp_1.0.7         sctransform_0.2.1  vctrs_0.3.8
[85] png_0.1-7          tidyselect_1.1.1   lmtest_0.9-38
```

[2]:

```
# THIS IS THE PREPROCESSING TO GET TO THE INTEGRATED SEURAT OBJECT.
#WT
wt_1_seu = readRDS(file = "../data/seurat_objects/seurat_raw_3_11_21/
 ↪sc_26_combined.rds")
wt_2_seu = readRDS(file = "../data/seurat_objects/seurat_raw_3_11_21/sc_67.rds")
YFP_1_seu = readRDS(file = "../data/seurat_objects/seurat_raw_3_11_21/sc_101.
 ↪rds")
YFP_2_seu = readRDS(file = "../data/seurat_objects/seurat_raw_3_11_21/sc_103.
 ↪rds")

#set experimental condition
wt_1_seu@meta.data$condition = "wt_unsorted"
wt_2_seu@meta.data$condition = "wt_unsorted"
YFP_1_seu@meta.data$condition = "wt_sorted"
YFP_2_seu@meta.data$condition = "wt_sorted"

#set batch
wt_1_seu@meta.data$batch = "1"
wt_2_seu@meta.data$batch = "2"
YFP_1_seu@meta.data$batch  = "3"
YFP_2_seu@meta.data$batch  = "3"

# THIS IS THE PREPROCESSING TO GET TO THE INTEGRATED SEURAT OBJECT.
#MUTANT
mut_1_seu = readRDS(file = "../data/seurat_objects/seurat_raw_3_11_21/
 ↪sc_27_combined.rds")
mut_2_seu = readRDS(file = "../data/seurat_objects/seurat_raw_3_11_21/sc_68.
 ↪rds")
KE_1_seu = readRDS(file = "../data/seurat_objects/seurat_raw_3_11_21/sc_102.
 ↪rds")
KE_2_seu = readRDS(file = "../data/seurat_objects/seurat_raw_3_11_21/sc_104.
 ↪rds")

#set experimental condition
```

```
mut_1_seu@meta.data$condition = "mut_unsorted"
mut_2_seu@meta.data$condition = "mut_unsorted"
KE_1_seu@meta.data$condition = "mut_sorted"
KE_2_seu@meta.data$condition = "mut_sorted"

#set batch
mut_1_seu@meta.data$batch  = "1"
mut_2_seu@meta.data$batch  = "2"
KE_1_seu@meta.data$batch = "3"
KE_2_seu@meta.data$batch = "3"
```

[ ]:

```
#SCtransform
#WT
wt_1_seu = SCTransform(wt_1_seu)
wt_2_seu = SCTransform(wt_2_seu)
YFP_1_seu = SCTransform(YFP_1_seu)
YFP_2_seu = SCTransform(YFP_2_seu)

#mutant
mut_1_seu = SCTransform(mut_1_seu)
mut_2_seu = SCTransform(mut_2_seu)
KE_1_seu = SCTransform(KE_1_seu)
KE_2_seu = SCTransform(KE_2_seu)
```

[ ]:

```
#Integrate
seu_intd_wt_mut = seu_integrate(wt_1_seu, wt_2_seu, YFP_1_seu, YFP_2_seu,␣
↪mut_2_seu, mut_1_seu, KE_1_seu, KE_2_seu, filename = "4_12_22_WT_mut",␣
↪nfeatures = 3000)
```

[2]:

```
#Load object
seu_intd_wt_mut = readRDS(file = "../data/intd_seu_objects/4_12_22_WT_mut.rds")
```

[ ]:

```
#clustera and UMAP embed
resolution = .75
set.seed(42)
DefaultAssay(seu_intd_wt_mut) <- "integrated"
options(repr.plot.width=12, repr.plot.height=12)
# Run the standard workflow for visualization and clustering
seu_intd_wt_mut <- RunPCA(seu_intd_wt_mut, npcs = 100, verbose = FALSE, approx␣
↪= FALSE)
seu_intd_wt_mut <- FindNeighbors(seu_intd_wt_mut, dims = 1:20, verbose = FALSE)
seu_intd_wt_mut <- FindClusters(seu_intd_wt_mut, resolution = resolution,␣
↪algorithm = 3, verbose = FALSE)
seu_intd_wt_mut <- RunUMAP(seu_intd_wt_mut, reduction = "pca", dims = 1:20,␣
↪verbose = FALSE)
```

[ ]:

```
#plot
options(repr.plot.width= 30, repr.plot.height=18)
DimPlot(seu_intd_wt_mut, reduction = "umap", label = FALSE, pt.size = 2, split.
↪by = "geno")#, cols = c("0" = "red"))
```

[ ]:

```
#Sweep acrss a few clustering resoluutions
res = c(seq(.25, .75, .25))
res

options(repr.plot.width= 30, repr.plot.height=18)

for (r in res) {
    resolution = r
    set.seed(42)
    DefaultAssay(seu_intd_wt_mut) <- "integrated"
    # Run the standard workflow for visualization and clustering
    seu_intd_wt_mut <- RunPCA(seu_intd_wt_mut, npcs = 100, verbose = FALSE,␣
↪approx = FALSE)
    seu_intd_wt_mut <- FindNeighbors(seu_intd_wt_mut, dims = 1:20, verbose =␣
↪FALSE)
    seu_intd_wt_mut <- FindClusters(seu_intd_wt_mut, resolution = resolution,␣
↪algorithm = 3, verbose = FALSE)
    seu_intd_wt_mut <- RunUMAP(seu_intd_wt_mut, reduction = "pca", dims = 1:20,␣
↪verbose = FALSE)
    plot = DimPlot(seu_intd_wt_mut, reduction = "umap", label = TRUE, pt.size =␣
↪2, split.by = "geno")#, cols = c("0" = "red"))
    print(plot)
    ggsave(file=paste0("../data/for_figures/UMAPs/","res_sweep",as.
↪character(r),".png"), plot=plot, width=20, height=10)

}
```
