## Supplemental Code for "Arabidopsis uses a molecular grounding mechanism and a biophysical circuit breaker to limit floral abscission signaling": notebook_03_nT_AZ_annotation.pdf

### WB03_WT_AZ_annotation

June 30, 2022

[1]:
```
#THIS SCRIPT PERFORMS AZ CLUSTER IDENTIFICATION

suppressMessages(library(Seurat))
library(ggplot2)

bulk_data = read.csv("../../data/buckets/single_cell_bucket_3_4_21/IWT_RNA_seq/
 ↪scRNA_flowers/outputs/bulk_edger_10_16_20.csv")

annotations = read.csv("R_functions/gene_descriptions.csv", header = F)
colnames(annotations) = c("gene_id", "description")
annotations$gene_id = substr(annotations$gene_id, 1, 9)

bp = read.csv("../data/shiny_go_analysis/figure_3/bp.csv")
cc = read.csv("../data/shiny_go_analysis/figure_3/cc.csv")
mf = read.csv("../data/shiny_go_analysis/figure_3/mf.csv")
```

[ ]:
```
options(repr.plot.width= 20, repr.plot.height=10)
DimPlot(seu_intd_wt_mut, reduction = "umap", label = TRUE, pt.size = 2, split.
↪by = "geno")#, cols = c("0" = "red"))
```

[ ]:
```
seu_intd_wt = subset(seu_intd_wt_mut, subset = geno == "WT")
```

[ ]:
```
#findmarkers
cluster_AZ_all = FindAllMarkers(seu_intd_wt,  logfc.threshold = 0, max.cells.
↪per.ident = 1000)
```

[ ]:
```
write.csv(cluster_AZ_all, file = paste("../data/for_figures/",␣
↪"AZ_markers_WT_ALL_res_75_April_25_22", ".csv", sep=""))
```

[ ]:
```
cluster_AZ_all = readRDS(paste0("../data/markers/",␣
↪"AZ_markers_WT_ALL_res_75_April_19_22", ".rds"))
```

[ ]:
```
head(cluster_AZ_all[cluster_AZ_all$cluster == 11,])
```

[ ]:
```
#FOR GO ANALYSIS
#write AZ specific genes as well as all genes with high enough expression to be␣
↪included in the analysis (ie the universe of genes for gene set testing)
write.csv(cluster_AZ_all[cluster_AZ_all$cluster == 11,], file = paste("../data/
↪for_figures/", "AZ_spec_genes_universe_WT_res_75_4_25_22", ".csv", sep=""),␣
↪row.names = FALSE)
write.csv(unique(cluster_AZ_all$gene), file = paste("../data/for_figures/",␣
↪"WT_universe_spec_genes_WT_res_75_4_25_22", ".csv", sep=""), row.names =␣
↪FALSE)
```

[1]:
```
#QRT2 data
kwak_ptpms=read.csv("../data/counts/kwak_ptpms.csv")
rownames(kwak_ptpms) = kwak_ptpms$X
kwak_ptpms[,c(1,2,3)] =NULL
colnames(kwak_ptpms) = "counts"

#HAE_YFP sorted
YFP_KE = read.csv("../data/counts/HAE_sorted.csv")
YFP_av = data.frame(YFP_KE[,2])
rownames(YFP_av) = YFP_KE[,1]
```

[ ]:
```
DefaultAssay(seu_intd_wt) = "RNA"
```

[ ]:
```
#get pseudobulk for each cluster to compare with kwak data
pbs = list()
count = 1
for (l in levels(seu_intd_wt@meta.data$seurat_clusters)) {
    pbs[[count]] = rowSums(as.matrix(GetAssayData(seu_intd_wt, slot =␣
↪"counts")[, WhichCells(seu_intd_wt, ident = l)]))
    count = count + 1
}

saveRDS(pbs, "../data/counts/cluster_pbs_4_13_22")
```

[ ]:
```
pbs = readRDS("../data/counts/cluster_pbs_4_13_22")
```

[ ]:
```
#convert pseudobulk to TPM
count = 1
for (c in pbs) {
    pbs[[count]] = data.frame(pbs[[count]])/sum(data.
 ↪frame(pbs[[count]]))*1000000
    rns = rownames(pbs[[count]])
    pbs[[count]] = pbs[[count]][order(rns),, drop = FALSE]
    count = count + 1
}
```

[ ]:
```
#QRT2
#set dataset
dataset = kwak_ptpms
cors_spearman = vector()
count = 1

seu_intd_wt@meta.data$kwak_cor = NULL

for (cluster in c(1:length(levels(seu_intd_wt@meta.data$seurat_clusters)))){
    test = cbind(pbs[[cluster]][intersect(rownames(pbs[[cluster]]),␣
 ↪rownames(dataset)),],dataset[intersect(rownames(pbs[[cluster]]),␣
 ↪rownames(dataset)),])
    cors_spearman[count] = cor(log(test[,1]+.1), log(test[,2]+.1), method =␣
 ↪"spearman")
    count = count + 1
}

for (i in c(1:length(levels(seu_intd_wt@meta.data$seurat_clusters)))){
    seu_intd_wt@meta.data$kwak_cor[seu_intd_wt@meta.data$seurat_clusters ==␣
 ↪toString(i-1)] = cors_spearman[i]
}

plot = FeaturePlot(seu_intd_wt,  features = "kwak_cor", pt.size = 1.5, cols =␣
 ↪c("gray", "red"))
print(plot)
ggsave(file=paste0("../data/for_figures/UMAPs/kwak_cor_wt_2_1_22.png"),␣
 ↪plot=plot, width=10, height=10)
```

[ ]:
```
#HAE
#set dataset
dataset = YFP_av
cors_spearman = vector()
count = 1

seu_intd_wt@meta.data$HAE_YFP = NULL

for (cluster in c(1:length(levels(seu_intd_wt@meta.data$seurat_clusters)))){
```

```
    test = cbind(pbs[[cluster]][intersect(rownames(pbs[[cluster]]),␣
↪rownames(dataset)),],dataset[intersect(rownames(pbs[[cluster]]),␣
↪rownames(dataset)),])
    cors_spearman[count] = cor(log(test[,1]+.1), log(test[,2]+.1), method =␣
↪"spearman")
    count = count + 1
}

for (i in c(1:length(levels(seu_intd_wt@meta.data$seurat_clusters)))){
    seu_intd_wt@meta.data$HAE_YFP[seu_intd_wt@meta.data$seurat_clusters ==␣
↪toString(i-1)] = cors_spearman[i]
}

plot = FeaturePlot(seu_intd_wt,  features = "HAE_YFP", pt.size = 1.5, cols =␣
↪c("white", "red"))
print(plot)
ggsave(file=paste0("../data/for_figures/UMAPs/HAE_YFP_cor_wt_2_1_22.png"),␣
↪plot=plot, width=10, height=10)
```
