## Supplemental Code for "Arabidopsis uses a molecular grounding mechanism and a biophysical circuit breaker to limit floral abscission signaling": notebook_04_cell_annotation.pdf

### WB04_cell_annotation

June 30, 2022

[1]:
```
#THIS SCRIPT ATTEMPTS TO ANNOTATE OTHER CLUSTERS IN THE INTEGRATED OBJECT

suppressMessages(library(Seurat))
suppressMessages(library(tidyverse))
suppressMessages(library(dplyr))
suppressMessages(library(cowplot))
suppressMessages(library(data.table))
```

[2]:
```
sessionInfo()
```

```
R version 3.6.3 (2020-02-29)
Platform: x86_64-conda_cos6-linux-gnu (64-bit)
Running under: Ubuntu 20.04.2 LTS

Matrix products: default
BLAS/LAPACK: /home/robotmessenger810/anaconda3/envs/r_3/lib/libopenblasp-r0.3.9.so

locale:
 [1] LC_CTYPE=en_US.UTF-8       LC_NUMERIC=C
 [3] LC_TIME=en_US.UTF-8        LC_COLLATE=en_US.UTF-8
 [5] LC_MONETARY=en_US.UTF-8    LC_MESSAGES=en_US.UTF-8
 [7] LC_PAPER=en_US.UTF-8       LC_NAME=C
 [9] LC_ADDRESS=C               LC_TELEPHONE=C
[11] LC_MEASUREMENT=en_US.UTF-8 LC_IDENTIFICATION=C

attached base packages:
[1] stats     graphics  grDevices utils     datasets  methods   base

other attached packages:
 [1] data.table_1.14.2 cowplot_1.1.1     forcats_0.5.0     stringr_1.4.0
 [5] dplyr_1.0.7       purrr_0.3.4       readr_1.4.0       tidyr_1.1.3
 [9] tibble_3.1.6      ggplot2_3.3.5     tidyverse_1.3.0   Seurat_3.1.5

loaded via a namespace (and not attached):
 [1] nlme_3.1-147       tsne_0.1-3         fs_1.5.2           lubridate_1.7.8
 [5] RcppAnnoy_0.0.19   RColorBrewer_1.1-2 httr_1.4.2         repr_1.1.0
 [9] sctransform_0.2.1  tools_3.6.3        backports_1.4.1    utf8_1.2.2
[13] R6_2.5.1           irlba_2.3.5        KernSmooth_2.23-20 uwot_0.1.8
```

```
[17] DBI_1.1.2          lazyeval_0.2.2     colorspace_2.0-2   withr_2.4.3
[21] tidyselect_1.1.1   gridExtra_2.3      compiler_3.6.3     cli_3.1.0
[25] rvest_0.3.5        xml2_1.3.1         plotly_4.9.2.1     scales_1.1.1
[29] lmtest_0.9-38      ggridges_0.5.2     pbapply_1.5-0      pbdZMQ_0.3-6
[33] digest_0.6.29      base64enc_0.1-3    pkgconfig_2.0.3    htmltools_0.5.0
[37] dbplyr_1.4.2       readxl_1.3.1       htmlwidgets_1.5.1  rlang_0.4.12
[41] rstudioapi_0.13    generics_0.1.1     zoo_1.8-8          jsonlite_1.7.2
[45] ica_1.0-2          magrittr_2.0.1     patchwork_1.1.1    Matrix_1.2-18
[49] Rcpp_1.0.7         IRkernel_1.1       munsell_0.5.0      fansi_0.5.0
[53] ape_5.3            reticulate_1.16    lifecycle_1.0.1    stringi_1.7.6
[57] MASS_7.3-54        Rtsne_0.15         plyr_1.8.6         grid_3.6.3
[61] parallel_3.6.3     listenv_0.8.0      ggrepel_0.9.1      crayon_1.4.2
[65] lattice_0.20-45    haven_2.3.1        IRdisplay_0.7.0    splines_3.6.3
[69] hms_1.1.0          pillar_1.6.4       igraph_1.2.5       uuid_0.1-4
[73] future.apply_1.6.0 reshape2_1.4.4     codetools_0.2-18   leiden_0.3.9
[77] reprex_0.3.0       glue_1.6.0         evaluate_0.14      modelr_0.1.6
[81] png_0.1-7          vctrs_0.3.8        cellranger_1.1.0   gtable_0.3.0
[85] RANN_2.6.1         future_1.18.0      assertthat_0.2.1   rsvd_1.0.3
[89] broom_0.5.5        survival_3.1-12    viridisLite_0.4.0  cluster_2.1.2
[93] globals_0.12.5     fitdistrplus_1.1-1 ellipsis_0.3.2     ROCR_1.0-11
```

[ ]:
```
seu_intd_wt_mut = readRDS(file = "../data/intd_seu_objects/4_12_22_WT_mut.rds")
```

[ ]:
```
resolution = .75
set.seed(42)
DefaultAssay(seu_intd_wt_mut) <- "integrated"
options(repr.plot.width=12, repr.plot.height=12)
# Run the standard workflow for visualization and clustering
#all_intd_sct <- ScaleData(all_intd_sct, verbose = FALSE)
seu_intd_wt_mut <- RunPCA(seu_intd_wt_mut, npcs = 100, verbose = FALSE, approx␣
↪= FALSE)
#From RunPCA doc: Features to compute PCA on. If features=NULL, PCA will be run␣
↪using the variable features for the Assay.
#Note that the features must be present in the scaled data. Any requested␣
↪features that are not scaled or have 0 variance
#will be dropped, and the PCA will be run using the remaining features.

#previously run 20 PCs as of 2/14/22
seu_intd_wt_mut <- FindNeighbors(seu_intd_wt_mut, dims = 1:20, verbose = FALSE)
seu_intd_wt_mut <- FindClusters(seu_intd_wt_mut, resolution = resolution,␣
↪algorithm = 3, verbose = FALSE)
seu_intd_wt_mut <- RunUMAP(seu_intd_wt_mut, reduction = "pca", dims = 1:20,␣
↪verbose = FALSE)
```

[ ]:
```
options(repr.plot.width= 20, repr.plot.height=10)
```

```
DimPlot(seu_intd_wt_mut, reduction = "umap", label = TRUE, pt.size = 2, split.
 ↪by = "geno")
```

[ ]:
```
#seu_intd_wt = subset(seu_intd_wt_mut, subset = geno == "WT")
seu_intd_wt = seu_intd_wt_mut
```

[ ]:
```
known.good.markers <- read.csv("../data/cell_type_markers/markers.csv", header␣
 ↪= F)
colnames(known.good.markers) = c("Name", "Locus", "Celltype")
known.good.markers <- known.good.markers[known.good.markers$Locus %in%␣
 ↪rownames(seu_intd_wt@assays$RNA),]
known.good.markers$Celltype <- gsub("abscission_zone", "Abscission Zone", known.
 ↪good.markers$Celltype, ignore.case = FALSE, perl = FALSE,fixed = T, useBytes␣
 ↪= FALSE);
known.good.markers$Celltype <- gsub("companion_cells", "Companion Cells", known.
 ↪good.markers$Celltype, ignore.case = FALSE, perl = FALSE,fixed = T, useBytes␣
 ↪= FALSE);
known.good.markers$Celltype <- gsub("xylem", "Xylem", known.good.
 ↪markers$Celltype, ignore.case = FALSE, perl = FALSE,fixed = T, useBytes =␣
 ↪FALSE);
known.good.markers$Celltype <- gsub("phloem", "Phloem", known.good.
 ↪markers$Celltype, ignore.case = FALSE, perl = FALSE,fixed = T, useBytes =␣
 ↪FALSE);
known.good.markers$Celltype <- gsub("vascular_subtype_1", "Vascular Subtype",␣
 ↪known.good.markers$Celltype, ignore.case = FALSE, perl = FALSE,fixed = T,␣
 ↪useBytes = FALSE);
known.good.markers$Celltype <- gsub("epidermis", "Epidermis", known.good.
 ↪markers$Celltype, ignore.case = FALSE, perl = FALSE,fixed = T, useBytes =␣
 ↪FALSE);
known.good.markers$Celltype <- gsub("sieve_element", "Sieve Element", known.
 ↪good.markers$Celltype, ignore.case = FALSE, perl = FALSE,fixed = T, useBytes␣
 ↪= FALSE);
known.good.markers$Celltype <- gsub("tracheary_element", "Tracheary Element",␣
 ↪known.good.markers$Celltype, ignore.case = FALSE, perl = FALSE,fixed = T,␣
 ↪useBytes = FALSE);
known.good.markers$Celltype <- gsub("mesophyll", "Mesophyll", known.good.
 ↪markers$Celltype, ignore.case = FALSE, perl = FALSE,fixed = T, useBytes =␣
 ↪FALSE);
known.good.markers$Celltype <- gsub("guard_cells", "Guard Cells", known.good.
 ↪markers$Celltype, ignore.case = FALSE, perl = FALSE,fixed = T, useBytes =␣
 ↪FALSE);
```

[ ]:
```
known.good.markers = known.good.markers[!known.good.markers$Name %in% c("IDA",␣
 ↪"PGAZAT", "HSL2", "HAESA"),]
known.good.markers
```

[ ]:
```
AZ_new = c("AT3G44550", "AT3G59850", "AT5G03820", "AT1G68320")
AZ_new_names = c("FAR5", "PLL", "GDSL","MYB62")
AZ_new_df = data.frame(matrix(ncol = 3, nrow = 4))
colnames(AZ_new_df) = c("Name", "Locus", "Celltype")
AZ_new_df$Locus = AZ_new
AZ_new_df$Celltype= "Abscission Zone"
AZ_new_df$Name = AZ_new_names
known.good.markers = rbind(known.good.markers, AZ_new_df)
```

[ ]:
```
known.good.markers
```

[ ]:
```
#MAY NOT NEED THIS CELL

DefaultAssay(seu_intd_wt) = "SCT"
options(repr.plot.width=8, repr.plot.height=8)

for (g in as.character(known.good.markers$Locus)) {
    plot = (FeaturePlot(seu_intd_wt, feature = g, pt.size = 4, order = TRUE,␣
 ↪min =.50))
    ggsave(file=paste0("../data/for_figures/gene_plots/figure_2_pngs/
 ↪celltype_plots/", g, "_", known.good.markers[known.good.markers$Locus ==␣
 ↪g,]$Celltype  ,".png"), plot=plot, width=10, height=10)

}
```

[ ]:
```
#MAY NOT NEED THIS CELL

DefaultAssay(seu_intd_wt) = "SCT"
g = "AT3G01420"
plot = (FeaturePlot(seu_intd_wt, feature = g, pt.size = 4, order = TRUE, min =.
 ↪50))
options(repr.plot.width=8, repr.plot.height=8)
ggsave(file=paste0("../data/for_figures/gene_plots/figure_2_pngs/celltype_plots/
 ↪", g, "_silique"  ,".png"), plot=plot, width=10, height=10)
print(plot)
```

[ ]:
```
resolution = 2
set.seed(42)
DefaultAssay(seu_intd_wt) <- "integrated"
options(repr.plot.width=12, repr.plot.height=12)
# Run the standard workflow for visualization and clustering
#all_intd_sct <- ScaleData(all_intd_sct, verbose = FALSE)
seu_intd_wt <- RunPCA(seu_intd_wt, npcs = 100, verbose = FALSE, approx = FALSE)
#From RunPCA doc: Features to compute PCA on. If features=NULL, PCA will be run␣
 ↪using the variable features for the Assay.
#Note that the features must be present in the scaled data. Any requested␣
 ↪features that are not scaled or have 0 variance
```

```
#will be dropped, and the PCA will be run using the remaining features.

#previously run 20 PCs as of 2/14/22
seu_intd_wt <- FindNeighbors(seu_intd_wt, dims = 1:20, verbose = FALSE)
seu_intd_wt <- FindClusters(seu_intd_wt, resolution = resolution, algorithm =␣
↪3, verbose = FALSE)
seu_intd_wt <- RunUMAP(seu_intd_wt, reduction = "pca", dims = 1:20, verbose =␣
↪FALSE)
```

[ ]:
```
zscore <- function(x){(x-mean(x))/sd(x)}

msc <- c()
for (i in as.character(unique(known.good.markers$Celltype))){
    if (length(known.good.markers[which(known.good.markers$Celltype==␣
↪i),]$Locus)>1){
    msc <- cbind(msc, as.numeric(apply(apply(seu_intd_wt@assays$SCT@data[known.
↪good.markers[which(known.good.markers$Celltype== i),]$Locus,], 1, zscore),␣
↪1, mean)))
    } else {
    msc <- cbind(msc, as.numeric(zscore(seu_intd_wt@assays$SCT@data[known.good.
↪markers[which(known.good.markers$Celltype== i),]$Locus,])))
    }

}
colnames(msc) <- as.character(unique(known.good.markers$Celltype))
rownames(msc) <- colnames(seu_intd_wt)
```

[ ]:
```
DefaultAssay(seu_intd_wt) <- "integrated"
suppressMessages(suppressWarnings(
  seu_intd_wt <- FindClusters(seu_intd_wt, resolution = 2, algorithm = 3)
))
```

[ ]:
```
anno <- seu_intd_wt$seurat_clusters
for (i in unique(seu_intd_wt$seurat_clusters)){
    if (max(apply(msc[which(seu_intd_wt$seurat_clusters==i),],2,mean))>0){
        ct <- names(which.
↪max(apply(msc[which(seu_intd_wt$seurat_clusters==i),],2,mean)))
    } else {
        ct <- "NA"
    }
        anno <- gsub(paste0("^",i,"$"), ct, anno, ignore.case = FALSE, perl =␣
↪FALSE,fixed = FALSE, useBytes = FALSE)
}
```

[ ]:
```
seu_intd_wt$score.crude.anno <- anno
```

[ ]:
```
# Plot marker annotation
order <- c("Abscission Zone", "Base of Sepals/Petals","Columella", "Lateral␣
↪Root Cap", "Atrichoblast", "Epidermis", "Mesophyll", "Guard Cells",␣
↪"Phloem","Sieve Element", "Xylem", "Vascular Subtype","Companion␣
↪Cells","Phloem Pole Pericycle", "Protoxylem", "Tracheary Element", "Unknown")
palette <- c("#9400d3", "#DCD0FF","#5ab953", "#bfef45", "#008080", "#21B6A8",␣
↪"#82b6ff", "#0000FF","#e6194b", "#dd77ec", "#9a6324", "#ffe119", "#ff9900",␣
↪"#ffd4e3", "#9a6324", "#ddaa6f", "#EEEEEE")
seu_intd_wt$score.crude.anno <- factor(seu_intd_wt$score.crude.anno , levels =␣
↪order[sort(match(unique(seu_intd_wt$score.crude.anno),order))])
color <- palette[sort(match(unique(seu_intd_wt$score.crude.anno),order))]
options(repr.plot.width=20, repr.plot.height=10)
DimPlot(seu_intd_wt, reduction = "umap", group.by = "score.crude.anno", split.
↪by = "geno", cols = color)+ggtitle("Z-Score Annotation Crude")
```

[ ]:
```
# Find clusters, here we choose Leiden clustering algorithm with resolution 0.5.
↪ Parameter "algorithm": 1 = original Louvain algorithm; 2 = Louvain␣
↪algorithm with multilevel refinement; 3 = SLM algorithm; 4 = Leiden algorithm
DefaultAssay(seu_intd_wt) <- "integrated"
suppressMessages(suppressWarnings(
  seu_intd_wt <- FindClusters(seu_intd_wt, resolution = 200, algorithm = 3)
))
```

[ ]:
```
anno <- seu_intd_wt$seurat_clusters
for (i in unique(seu_intd_wt$seurat_clusters)){
    if (max(apply(msc[which(seu_intd_wt$seurat_clusters==i),],2,mean))>0){
        ct <- names(which.
↪max(apply(msc[which(seu_intd_wt$seurat_clusters==i),],2,mean)))
    } else {
        ct <- "NA"
    }
        anno <- gsub(paste0("^",i,"$"), ct, anno, ignore.case = FALSE, perl =␣
↪FALSE,fixed = FALSE, useBytes = FALSE)
}

seu_intd_wt$score.anno <- anno
# Plot marker annotation
order <- c("Abscission Zone", "Base of Sepals/Petals","Columella", "Lateral␣
↪Root Cap", "Atrichoblast", "Epidermis", "Mesophyll", "Guard Cells",␣
↪"Phloem","Sieve Element", "Xylem", "Vascular Subtype","Companion␣
↪Cells","Phloem Pole Pericycle", "Protoxylem", "Tracheary Element", "Unknown")
palette <- c("#9400d3", "#DCD0FF","#5ab953", "#bfef45", "#008080", "#21B6A8",␣
↪"#82b6ff", "#0000FF","#e6194b", "#dd77ec", "#9a6324", "#ffe119", "#ff9900",␣
↪"#ffd4e3", "#9a6324", "#ddaa6f", "#EEEEEE")
```

```
seu_intd_wt$score.anno <- factor(seu_intd_wt$score.anno , levels =␣
↪order[sort(match(unique(seu_intd_wt$score.anno),order))])
#color <- palette[sort(match(unique(seu_intd_wt$score.anno),order))]
#options(repr.plot.width=12, repr.plot.height=10)
#DimPlot(seu_intd_wt, reduction = "umap", group.by = "score.anno", cols =␣
↪color)+ggtitle("Z-Score Annotation")
```

[ ]:
```
#Consensus Annotation
dat <- data.frame(seu_intd_wt$score.anno, seu_intd_wt$score.crude.anno)
seu_intd_wt$consensus.anno <- apply(dat,1,function(x){if (is.
↪na(x[1])){"Unknown"} else if (is.na(x[2])){"Unknown"} else if␣
↪(x[1]==x[2]){x[1]} else {"Unknown"}})
#seu_intd_wt$consensus.anno <- apply(dat,1,function(x){if(x[1]==x[2]){x[1]}else␣
↪if(x[1]=="Trichoblast" & x[2]=="Atrichoblast"){"Trichoblast"}
#    else if(x[1]=="Late Metaxylem" & x[2]=="Phloem"){"Late Metaxylem"}else␣
↪if(x[1]=="Cortex" & x[2]=="Sclerenchyma"){"Cortex"}
#        else if(x[1]=="Exodermis" & x[2]=="Sclerenchyma"){"Exodermis"}else␣
↪if(x[1]=="Exodermis" & x[2]=="Endodermis"){"Exodermis"}
#            else if(x[1]=="Cortex" & x[2]=="Endodermis"){"Cortex"}else␣
↪if(x[1]=="Pericycle" & x[2]=="Endodermis"){"Pericycle"}else␣
↪if(x[1]=="Phloem" & x[2]=="Endodermis"){"Phloem"}
#                    else if(x[1]=="Late Metaxylem" & x[2]=="Endodermis"){"Late␣
↪Metaxylem"}else if(x[3]=="Maturation1"|x[3]=="Maturation2"){x[2]}else␣
↪{"Unknown"}})
order <- c("Abscission Zone", "Base of Sepals/Petals","Columella", "Lateral␣
↪Root Cap", "Atrichoblast", "Epidermis", "Mesophyll", "Guard Cells",␣
↪"Phloem","Sieve Element", "Xylem", "Vascular Subtype","Companion␣
↪Cells","Phloem Pole Pericycle", "Protoxylem", "Tracheary Element", "Unknown")
palette <- c("#9400d3", "#DCD0FF","#5ab953", "#bfef45", "#008080", "#21B6A8",␣
↪"#82b6ff", "#0000FF","#e6194b", "#dd77ec", "#9a6324", "#ffe119", "#ff9900",␣
↪"#ffd4e3", "#9a6324", "#ddaa6f", "#EEEEEE")
seu_intd_wt$consensus.anno <- factor(seu_intd_wt$consensus.anno , levels =␣
↪order[sort(match(unique(seu_intd_wt$consensus.anno),order))])
color <- palette[sort(match(unique(seu_intd_wt$consensus.anno),order))]
options(repr.plot.width=10, repr.plot.height=10)
DimPlot(seu_intd_wt, reduction = "umap", group.by = "consensus.anno", pt.size =␣
↪3, cols = color)+ggtitle("consensus.anno")

seu_intd_wt$celltype.consensus.anno <- seu_intd_wt$consensus.anno
```

[ ]:
```
options(repr.plot.width=20, repr.plot.height=10)
DimPlot(seu_intd_wt, reduction = "umap", group.by = "consensus.anno", pt.size =␣
↪3, split.by = "geno",cols = color)+ggtitle("consensus.anno")

seu_intd_wt$celltype.consensus.anno <- seu_intd_wt$consensus.anno
```

[ ]:
```
color <- palette[sort(match( c("Mesophyll", "Phloem", "Xylem", "Guard Cells",␣
↪"Companion Cells", "Epidermis", "Abscission Zone", "Unknown"), order))]
seu_intd_wt@meta.data[seu_intd_wt@meta.data$consensus.anno == "Sieve␣
↪Element",]$consensus.anno = "Unknown"
seu_intd_wt@meta.data[seu_intd_wt@meta.data$consensus.anno == "Tracheary␣
↪Element",]$consensus.anno = "Unknown"
seu_intd_wt@meta.data[seu_intd_wt@meta.data$consensus.anno == "Vascular␣
↪Subtype",]$consensus.anno = "Unknown"
```

[ ]:
```
#c("#9400d3", ,"#5ab953", , , "#21B6A8", "#0000FF",, "#dd77ec", "#9a6324",␣
↪"#ffe119", "#ff9900", "#ffd4e3", "#9a6324", "#ddaa6f", "#EEEEEE")
options(repr.plot.width= 20, repr.plot.height=10)
palette <- c("#EEEEEE", "#dd77ec", "#42f5ef", "#f542ef", "#ff9900", "#42f548",␣
↪"#0000FF", "#f56642")
plot = DimPlot(seu_intd_wt, reduction = "umap", group.by = "consensus.anno", ␣
↪split.by = "geno", order = c("Mesophyll", "Phloem", "Xylem", "Guard Cells",␣
↪"Companion Cells", "Epidermis", "Abscission Zone", "Unknown"), pt.size = 4,␣
↪cols =palette)+ggtitle("consensus.anno")
ggsave(file=paste0("../data/for_figures/gene_plots/figure_2_pngs/celltype_plots/
↪celltype_UMAP.png"), plot=plot, width=20, height=10)
print(plot)
```
