## Supplemental Code for "Arabidopsis uses a molecular grounding mechanism and a biophysical circuit breaker to limit floral abscission signaling": notebook_05_pseudo_bulk.pdf

### WB05_pseudo_bulk

June 30, 2022

[1]:
```
#THIS SCRIPT PERFORMS PSEUDOBULK ANALYSIS ON THE AZ IN WT AND MUTANT

suppressMessages(library(Seurat))
library(here)
source(here("R_functions","edgeR_function.R"))

annotations = read.csv("R_functions/gene_descriptions.csv", header = F)
colnames(annotations) = c("gene_id", "description")
annotations$gene_id = substr(annotations$gene_id, 1, 9)

proto_genes=read.csv("../data/bulk_data/protoplasting.csv")
proto_list=as.character(proto_genes[abs(proto_genes$logFC) > 1,]$genes)
```

```
here() starts at /home/robotmessenger810/sc_analysis/code
```

[2]:
```
sessionInfo()
```

```
R version 3.6.3 (2020-02-29)
Platform: x86_64-conda_cos6-linux-gnu (64-bit)
Running under: Ubuntu 20.04.2 LTS

Matrix products: default
BLAS/LAPACK: /home/robotmessenger810/anaconda3/envs/r_3/lib/libopenblasp-r0.3.9.so

locale:
 [1] LC_CTYPE=en_US.UTF-8       LC_NUMERIC=C
 [3] LC_TIME=en_US.UTF-8        LC_COLLATE=en_US.UTF-8
 [5] LC_MONETARY=en_US.UTF-8    LC_MESSAGES=en_US.UTF-8
 [7] LC_PAPER=en_US.UTF-8       LC_NAME=C
 [9] LC_ADDRESS=C               LC_TELEPHONE=C
[11] LC_MEASUREMENT=en_US.UTF-8 LC_IDENTIFICATION=C

attached base packages:
[1] stats     graphics  grDevices utils     datasets  methods   base

other attached packages:
[1] here_0.1     Seurat_3.1.5
```

```
loaded via a namespace (and not attached):
 [1] httr_1.4.2          tidyr_1.1.3         jsonlite_1.7.2      viridisLite_0.4.0
 [5] splines_3.6.3       leiden_0.3.9        ggrepel_0.9.1       globals_0.12.5
 [9] pillar_1.6.4        lattice_0.20-45     glue_1.6.0          reticulate_1.16
[13] uuid_0.1-4          digest_0.6.29       RColorBrewer_1.1-2 colorspace_2.0-2
[17] cowplot_1.1.1       htmltools_0.5.0     Matrix_1.2-18       plyr_1.8.6
[21] pkgconfig_2.0.3     tsne_0.1-3          listenv_0.8.0       purrr_0.3.4
[25] patchwork_1.1.1     scales_1.1.1        RANN_2.6.1          Rtsne_0.15
[29] tibble_3.1.6        generics_0.1.1      ggplot2_3.3.5       ellipsis_0.3.2
[33] repr_1.1.0          ROCR_1.0-11         pbapply_1.5-0       lazyeval_0.2.2
[37] survival_3.1-12     magrittr_2.0.1      crayon_1.4.2        evaluate_0.14
[41] future_1.18.0       fansi_0.5.0         nlme_3.1-147        MASS_7.3-54
[45] ica_1.0-2           tools_3.6.3         fitdistrplus_1.1-1 data.table_1.14.2
[49] lifecycle_1.0.1     stringr_1.4.0       plotly_4.9.2.1      munsell_0.5.0
[53] cluster_2.1.2       irlba_2.3.5         compiler_3.6.3      rsvd_1.0.3
[57] rlang_0.4.12        grid_3.6.3          ggridges_0.5.2      pbdZMQ_0.3-6
[61] IRkernel_1.1        RcppAnnoy_0.0.19    htmlwidgets_1.5.1   igraph_1.2.5
[65] base64enc_0.1-3     gtable_0.3.0        codetools_0.2-18    DBI_1.1.2
[69] reshape2_1.4.4      R6_2.5.1            gridExtra_2.3       zoo_1.8-8
[73] dplyr_1.0.7         uwot_0.1.8          future.apply_1.6.0 utf8_1.2.2
[77] rprojroot_2.0.2     KernSmooth_2.23-20 ape_5.3             stringi_1.7.6
[81] parallel_3.6.3      IRdisplay_0.7.0     Rcpp_1.0.7          sctransform_0.2.1
[85] vctrs_0.3.8         png_0.1-7           tidyselect_1.1.1    lmtest_0.9-38
```

[ ]:

```
seu_intd_wt_mut = readRDS(file = "../data/intd_seu_objects/4_12_22_WT_mut.rds")

resolution = .75
set.seed(42)
DefaultAssay(seu_intd_wt_mut) <- "integrated"
options(repr.plot.width=12, repr.plot.height=12)
seu_intd_wt_mut <- RunPCA(seu_intd_wt_mut, npcs = 100, verbose = FALSE, approx␣
↪= FALSE)
seu_intd_wt_mut <- FindNeighbors(seu_intd_wt_mut, dims = 1:20, verbose = FALSE)
seu_intd_wt_mut <- FindClusters(seu_intd_wt_mut, resolution = resolution,␣
↪algorithm = 3, verbose = FALSE)
seu_intd_wt_mut <- RunUMAP(seu_intd_wt_mut, reduction = "pca", dims = 1:20,␣
↪verbose = FALSE)
```

[ ]:

```
cluster = "11" #AZ cluster
seu_intd_wt_mut@active.assay = "RNA"

wt_1_AZ <- rowSums(as.matrix(GetAssayData(subset(seu_intd_wt_mut, subset = orig.
↪ident == "sc_26_combined"), slot = "counts")[,␣
↪WhichCells(subset(seu_intd_wt_mut, subset = orig.ident == "sc_26_combined"),␣
↪ident = cluster)]))
```

```
wt_2_AZ <- rowSums(as.matrix(GetAssayData(subset(seu_intd_wt_mut, subset = orig.
↪ident == "sc_67"), slot = "counts")[, WhichCells(subset(seu_intd_wt_mut,␣
↪subset = orig.ident == "sc_67"), ident = cluster)]))
YFP_1_AZ <- rowSums(as.matrix(GetAssayData(subset(seu_intd_wt_mut, subset =␣
↪orig.ident == "sc_101"), slot = "counts")[,␣
↪WhichCells(subset(seu_intd_wt_mut, subset = orig.ident == "sc_101"), ident =␣
↪cluster)]))
YFP_2_AZ <- rowSums(as.matrix(GetAssayData(subset(seu_intd_wt_mut, subset =␣
↪orig.ident == "sc_103"), slot = "counts")[,␣
↪WhichCells(subset(seu_intd_wt_mut, subset = orig.ident == "sc_103"), ident =␣
↪cluster)]))
```

[ ]:

```
cluster = "11"
seu_intd_wt_mut@active.assay = "RNA"

mut_1_AZ <- rowSums(as.matrix(GetAssayData(subset(seu_intd_wt_mut, subset =␣
↪orig.ident == "sc_27_combined"), slot = "counts")[,␣
↪WhichCells(subset(seu_intd_wt_mut, subset = orig.ident == "sc_27_combined"),␣
↪ident = cluster)]))
mut_2_AZ <- rowSums(as.matrix(GetAssayData(subset(seu_intd_wt_mut, subset =␣
↪orig.ident == "sc_68"), slot = "counts")[,␣
↪WhichCells(subset(seu_intd_wt_mut, subset = orig.ident == "sc_68"), ident =␣
↪cluster)]))
KE_1_AZ <- rowSums(as.matrix(GetAssayData(subset(seu_intd_wt_mut, subset = orig.
↪ident == "sc_102"), slot = "counts")[, WhichCells(subset(seu_intd_wt_mut,␣
↪subset = orig.ident == "sc_102"), ident = cluster)]))
KE_2_AZ <- rowSums(as.matrix(GetAssayData(subset(seu_intd_wt_mut, subset = orig.
↪ident == "sc_104"), slot = "counts")[, WhichCells(subset(seu_intd_wt_mut,␣
↪subset = orig.ident == "sc_104"), ident = cluster)]))
```

[ ]:

```
gene_intersection = intersect(names(wt_1_AZ), names(mut_1_AZ))

wt_1_AZ = wt_1_AZ[gene_intersection]
wt_2_AZ = wt_2_AZ[gene_intersection]
YFP_1_AZ = YFP_1_AZ[gene_intersection]
YFP_2_AZ = YFP_2_AZ[gene_intersection]
mut_1_AZ = mut_1_AZ[gene_intersection]
mut_2_AZ = mut_2_AZ[gene_intersection]
KE_1_AZ = KE_1_AZ[gene_intersection]
KE_2_AZ = KE_2_AZ [gene_intersection]
```

[ ]:

```
pb_df = data.frame(cbind(wt_1_AZ , wt_2_AZ, YFP_1_AZ, YFP_2_AZ, mut_1_AZ ,␣
↪mut_2_AZ, KE_1_AZ, KE_2_AZ))
colnames(pb_df) = c("WT1", "WT2", "YFP1", "YFP2", "mut1", "mut2", "KE1", "KE2")
rownames(pb_df) = gene_intersection
```

[ ]:
```
write.csv(pb_df, "../data/pseudo_bulk_data/AZ_pbs_4_19_22.csv")
```

[ ]:
```
pb_df = read.csv("../data/pseudo_bulk_data/AZ_pbs_4_19_22.csv")
rownames(pb_df) = pb_df[,1]
pb_df[,1] <- NULL
```

[ ]:
```
#account for factors in experiment
phenotype=as.factor(c("wt", "wt", "wt", "wt", "mut", "mut", "mut", "mut"))
batch=as.factor(c(0,0,1,1,0,0,1,1))
design <- model.matrix(~phenotype+batch)

#double check design matrix isn't singular
print(paste("determinant of XT*X of design matrix is: ",␣
↪det(t(design)%*%(design))))

#making contrast matrix for tests of interest
my.contrasts <- makeContrasts(s1_v_s2=phenotypewt, levels=design)
```

[ ]:
```
my.contrasts
design
```

[ ]:
```
#put experimental covariates in
bulk_edger_1 =  edgeR_2_sample(pb_df, "WT", "mut", c(1,2,3,4), c(5,6,7,8),␣
↪annotations, design, my.contrasts)
```

[ ]:
```
WT_higher_1 = bulk_edger_1[bulk_edger_1$FDR < .2 & bulk_edger_1$logFC > 1,]
WT_lower_1 = bulk_edger_1[bulk_edger_1$FDR < .2 & bulk_edger_1$logFC < -1,]
```

[ ]:
```
dim(WT_higher_1)
```

[ ]:
```
write.csv(bulk_edger_1, "../data/pseudo_bulk_data/AZ_edger_4_19_22_factors.csv")
```

[ ]:
```
bulk_edger_1 = read.csv("../data/pseudo_bulk_data/AZ_edger_4_19_22_factors.csv")
write.csv(bulk_edger_1, "../data/for_figures/AZ_edger_4_19_22_factors.csv")
```

[ ]:
```
dim(bulk_edger_1)
```
