## Supplemental Code for "Arabidopsis uses a molecular grounding mechanism and a biophysical circuit breaker to limit floral abscission signaling": notebook_06_subclustering.pdf

### WB06_subclustering

June 30, 2022

[1]:
```
#THIS SCRIPT PERFORMS SUBCLUSTERING OF THE WT AND MUTANT AZ CELLS, IDENTIFIES␣
↪PUTATIVE SECESSION AND RESIDUUM CELLS, AND PERFORMS GENE ENRICHMENT ANALYSIS.

library(here)
library(Matrix)
library(tidyverse)
library(Seurat)
library(edgeR)
library(limma)
source(here("R_functions","edgeR_function.R"))

annotations = read.csv("R_functions/gene_descriptions.csv", header = F)
colnames(annotations) = c("gene_id", "description")
annotations$gene_id = substr(annotations$gene_id, 1, 9)

proto_genes=read.csv("../data/bulk_data/protoplasting.csv")
proto_list=as.character(proto_genes[abs(proto_genes$logFC) > 1,]$genes)
bulk_data = read.csv("/home/robotmessenger810/data/buckets/
↪single_cell_bucket_3_4_21/IWT_RNA_seq/scRNA_flowers/outputs/
↪bulk_edger_10_16_20.csv")
```

[2]:
```
sessionInfo()
```

```
R version 3.6.3 (2020-02-29)
Platform: x86_64-conda_cos6-linux-gnu (64-bit)
Running under: Ubuntu 20.04.2 LTS

Matrix products: default
BLAS/LAPACK: /home/robotmessenger810/anaconda3/envs/r_3/lib/libopenblasp-r0.3.9.so

locale:
 [1] LC_CTYPE=en_US.UTF-8       LC_NUMERIC=C
 [3] LC_TIME=en_US.UTF-8        LC_COLLATE=en_US.UTF-8
 [5] LC_MONETARY=en_US.UTF-8    LC_MESSAGES=en_US.UTF-8
 [7] LC_PAPER=en_US.UTF-8       LC_NAME=C
 [9] LC_ADDRESS=C               LC_TELEPHONE=C
[11] LC_MEASUREMENT=en_US.UTF-8 LC_IDENTIFICATION=C
```

```
attached base packages:
[1] stats     graphics  grDevices utils     datasets  methods   base

other attached packages:
 [1] edgeR_3.28.1    limma_3.42.2    Seurat_3.1.5    forcats_0.5.0
 [5] stringr_1.4.0   dplyr_1.0.7     purrr_0.3.4     readr_1.4.0
 [9] tidyr_1.1.3     tibble_3.1.6    ggplot2_3.3.5   tidyverse_1.3.0
[13] Matrix_1.2-18   here_0.1

loaded via a namespace (and not attached):
 [1] Rtsne_0.15          colorspace_2.0-2   ellipsis_0.3.2     ggridges_0.5.2
 [5] rprojroot_2.0.2     IRdisplay_0.7.0    base64enc_0.1-3    fs_1.5.2
 [9] rstudioapi_0.13     leiden_0.3.9       listenv_0.8.0      ggrepel_0.9.1
[13] fansi_0.5.0         lubridate_1.7.8    xml2_1.3.1         codetools_0.2-18
[17] splines_3.6.3       IRkernel_1.1       jsonlite_1.7.2     broom_0.5.5
[21] ica_1.0-2           cluster_2.1.2      dbplyr_1.4.2       png_0.1-7
[25] uwot_0.1.8          sctransform_0.2.1  compiler_3.6.3     httr_1.4.2
[29] backports_1.4.1     assertthat_0.2.1   lazyeval_0.2.2     cli_3.1.0
[33] htmltools_0.5.0     tools_3.6.3        rsvd_1.0.3         igraph_1.2.5
[37] gtable_0.3.0        glue_1.6.0         RANN_2.6.1         reshape2_1.4.4
[41] Rcpp_1.0.7          cellranger_1.1.0   vctrs_0.3.8        ape_5.3
[45] nlme_3.1-147        lmtest_0.9-38      globals_0.12.5     rvest_0.3.5
[49] lifecycle_1.0.1     irlba_2.3.5        future_1.18.0      MASS_7.3-54
[53] zoo_1.8-8           scales_1.1.1       hms_1.1.0          parallel_3.6.3
[57] RColorBrewer_1.1-2 reticulate_1.16    pbapply_1.5-0      gridExtra_2.3
[61] stringi_1.7.6       repr_1.1.0         rlang_0.4.12       pkgconfig_2.0.3
[65] evaluate_0.14       lattice_0.20-45    ROCR_1.0-11        patchwork_1.1.1
[69] htmlwidgets_1.5.1   cowplot_1.1.1      tidyselect_1.1.1   RcppAnnoy_0.0.19
[73] plyr_1.8.6          magrittr_2.0.1     R6_2.5.1           generics_0.1.1
[77] pbdZMQ_0.3-6        DBI_1.1.2          pillar_1.6.4       haven_2.3.1
[81] withr_2.4.3         fitdistrplus_1.1-1 survival_3.1-12    future.apply_1.6.0
[85] tsne_0.1-3          modelr_0.1.6       crayon_1.4.2       uuid_0.1-4
[89] KernSmooth_2.23-20 utf8_1.2.2         plotly_4.9.2.1     locfit_1.5-9.4
[93] grid_3.6.3          readxl_1.3.1       data.table_1.14.2  reprex_0.3.0
[97] digest_0.6.29       munsell_0.5.0      viridisLite_0.4.0
```

[3]:
```
seu_intd_wt_mut = readRDS(file = "../data/intd_seu_objects/4_12_22_WT_mut.rds")
```

[ ]:
```
resolution = .75
set.seed(42)
DefaultAssay(seu_intd_wt_mut_mut) <- "integrated"
options(repr.plot.width=12, repr.plot.height=12)
# Run the standard workflow for visualization and clustering
seu_intd_wt_mut_mut <- RunPCA(seu_intd_wt_mut_mut, npcs = 100, verbose = FALSE,␣
↪approx = FALSE)
```

```
seu_intd_wt_mut_mut <- FindNeighbors(seu_intd_wt_mut_mut, dims = 1:20, verbose␣
↪= FALSE)
seu_intd_wt_mut_mut <- FindClusters(seu_intd_wt_mut_mut, resolution =␣
↪resolution, algorithm = 3, verbose = FALSE)
seu_intd_wt_mut_mut <- RunUMAP(seu_intd_wt_mut_mut, reduction = "pca", dims = 1:
↪20, verbose = FALSE)
```

[ ]:
```
cluster = "11"

wt_1_AZ <- subset(seu_intd_wt_mut, subset = orig.ident == "sc_26_combined")[,␣
↪WhichCells(subset(seu_intd_wt_mut, subset = orig.ident == "sc_26_combined"),␣
↪ident = cluster)]
wt_2_AZ <- subset(seu_intd_wt_mut, subset = orig.ident == "sc_67")[,␣
↪WhichCells(subset(seu_intd_wt_mut, subset = orig.ident == "sc_67"), ident =␣
↪cluster)]
YFP_1_AZ <- subset(seu_intd_wt_mut, subset = orig.ident == "sc_101")[,␣
↪WhichCells(subset(seu_intd_wt_mut, subset = orig.ident == "sc_101"), ident =␣
↪cluster)]
YFP_2_AZ <- subset(seu_intd_wt_mut, subset = orig.ident == "sc_103")[,␣
↪WhichCells(subset(seu_intd_wt_mut, subset = orig.ident == "sc_103"), ident =␣
↪cluster)]
```

[ ]:
```
wt_1_seu = SCTransform(wt_1_AZ)
wt_2_seu = SCTransform(wt_2_AZ)
YFP_1_seu = SCTransform(YFP_1_AZ)
YFP_2_seu = SCTransform(YFP_2_AZ)
```

[ ]:
```
seu_intd_wt_AZ = seu_integrate(wt_1_seu, wt_2_seu, YFP_1_seu, YFP_2_seu,␣
↪filename = "AZ_only_WT_4_19_22", nfeatures = 3000)
```

[2]:
```
seu_intd_wt_AZ = readRDS("../data/intd_seu_objects/AZ_only_WT_4_19_22.rds")
```

[ ]:
```
resolution = .1
set.seed(42)
DefaultAssay(seu_intd_wt_AZ) <- "integrated"
options(repr.plot.width=12, repr.plot.height=12)
# Run the standard workflow for visualization and clustering
seu_intd_wt_AZ <- ScaleData(seu_intd_wt_AZ, verbose = FALSE)
seu_intd_wt_AZ <- RunPCA(seu_intd_wt_AZ, npcs = 100, verbose = FALSE, approx =␣
↪FALSE)
seu_intd_wt_AZ <- FindNeighbors(seu_intd_wt_AZ, dims = 1:20, verbose = FALSE)
seu_intd_wt_AZ <- FindClusters(seu_intd_wt_AZ, resolution = resolution,␣
↪algorithm = 3, verbose = FALSE)
seu_intd_wt_AZ <- RunUMAP(seu_intd_wt_AZ, reduction = "pca", dims = 1:20,␣
↪verbose = FALSE)
```

[23]:
```
options(repr.plot.width=8, repr.plot.height=8)
DefaultAssay(seu_intd_wt_AZ) = "SCT"
plot = FeaturePlot(seu_intd_wt_AZ, "AT4G28490", min = .5,  pt.size = 6, order =␣
↪T)
ggsave(file="../data/for_figures/UMAPs/AZ_WT_HAE_featureplot.png", plot=plot,␣
↪width=8, height=8)
plot = FeaturePlot(seu_intd_wt_AZ, "AT1G68765",  min = .5,  pt.size = 6, order␣
↪= T)
ggsave(file="../data/for_figures/UMAPs/AZ_WT_IDA_featureplot.png", plot=plot,␣
↪width=8, height=8)
```

[ ]:
```
options(repr.plot.width= 10, repr.plot.height=10)
plot = DimPlot(seu_intd_wt_AZ, reduction = "umap", label = TRUE, pt.size = 4)
print(plot)
ggsave(file="../data/for_figures/UMAPs/AZ_WT_UMAP.png", plot=plot, width=10,␣
↪height=10)
```

[ ]:
```
DefaultAssay(seu_intd_wt_AZ) <- "RNA"

#get pseudobulk for each cluster to compare with kwak data
pbs = list()
count = 1
for (l in levels(seu_intd_wt_AZ@meta.data$seurat_clusters)) {
    pbs[[count]] = rowSums(as.matrix(GetAssayData(seu_intd_wt_AZ, slot =␣
↪"counts")[,WhichCells(seu_intd_wt_AZ, ident = l)]))
    count = count + 1
}

#saveRDS(pbs, "../data/counts/AZ_wt_cluster_pbs_3_1_22")
```

[ ]:
```
#convert pseudobulk to TPM
count = 1
for (c in pbs) {
    pbs[[count]] = data.frame(pbs[[count]])/sum(data.
↪frame(pbs[[count]]))*1000000
    rns = rownames(pbs[[count]])
    pbs[[count]] = pbs[[count]][order(rns),, drop = FALSE]
    count = count + 1
}
```

[ ]:
```
#secession
kwak_ptpms_raw=read.csv("../data/counts/kwak_ptpms.csv")
rownames(kwak_ptpms_raw) = kwak_ptpms_raw$X
kwak_ptpms = kwak_ptpms_raw
kwak_ptpms[,c(1,3,4)] =NULL

#secession
```

```
#set dataset
dataset = kwak_ptpms

cors_spearman = vector()
count = 1

seu_intd_wt_AZ@meta.data$kwak_cor = NULL

for (cluster in c(1:length(levels(seu_intd_wt_AZ@meta.data$seurat_clusters)))){
    test = cbind(pbs[[cluster]][intersect(rownames(pbs[[cluster]]),␣
↪rownames(dataset)),],dataset[intersect(rownames(pbs[[cluster]]),␣
↪rownames(dataset)),])
    cors_spearman[count] = cor(log(test[,1]+.1), log(test[,2]+.1), method =␣
↪"spearman")
    count = count + 1
}

for (i in c(1:length(levels(seu_intd_wt_AZ@meta.data$seurat_clusters)))){
    seu_intd_wt_AZ@meta.data$kwak_cor[seu_intd_wt_AZ@meta.data$seurat_clusters␣
↪== toString(i-1)] = cors_spearman[i]
}

plot = FeaturePlot(seu_intd_wt_AZ,  features = "kwak_cor", pt.size = 4, cols =␣
↪c("light gray", "red"))
print(plot)
ggsave(file="../data/for_figures/UMAPs/AZ_WT_sec_UMAP.png", plot=plot,␣
↪width=10, height=10)
```

```
[ ]: #residuum
kwak_ptpms_raw=read.csv("../data/counts/kwak_ptpms.csv")
rownames(kwak_ptpms_raw) = kwak_ptpms_raw$X
kwak_ptpms = kwak_ptpms_raw
kwak_ptpms[,c(1,2,4)] =NULL

#residuum
#set dataset
dataset = kwak_ptpms

cors_spearman = vector()
count = 1

seu_intd_wt_AZ@meta.data$kwak_cor = NULL

for (cluster in c(1:length(levels(seu_intd_wt_AZ@meta.data$seurat_clusters)))){
    test = cbind(pbs[[cluster]][intersect(rownames(pbs[[cluster]]),␣
↪rownames(dataset)),],dataset[intersect(rownames(pbs[[cluster]]),␣
↪rownames(dataset)),])
```

```
    cors_spearman[count] = cor(log(test[,1]+.1), log(test[,2]+.1), method =␣
↪"spearman")
    count = count + 1
}

for (i in c(1:length(levels(seu_intd_wt_AZ@meta.data$seurat_clusters)))){
    seu_intd_wt_AZ@meta.data$kwak_cor[seu_intd_wt_AZ@meta.data$seurat_clusters␣
↪== toString(i-1)] = cors_spearman[i]
}

plot = FeaturePlot(seu_intd_wt_AZ,  features = "kwak_cor", pt.size = 4, cols =␣
↪c("light gray", "red"))
print(plot)
ggsave(file="../data/for_figures/UMAPs/AZ_WT_res_UMAP.png", plot=plot,␣
↪width=10, height=10)
```

[ ]:
```
DefaultAssay(seu_intd_wt_AZ) <- "RNA"
wt_sec_v_rec = data.frame(matrix(ncol = 8, nrow␣
↪=dim(seu_intd_wt_AZ@assays$RNA)[1]))
wt_sec_v_rec_red = data.frame(matrix(ncol = 6, nrow␣
↪=dim(seu_intd_wt_AZ@assays$RNA)[1]))

res1_1 = rowSums(as.matrix(GetAssayData(subset(seu_intd_wt_AZ, subset = orig.
↪ident == "sc_26_combined"), slot = "counts")[,␣
↪WhichCells(subset(seu_intd_wt_AZ, subset = orig.ident == "sc_26_combined"),␣
↪ident = "0")]))
res2_1 = rowSums(as.matrix(GetAssayData(subset(seu_intd_wt_AZ, subset = orig.
↪ident == "sc_67"), slot = "counts")[, WhichCells(subset(seu_intd_wt_AZ,␣
↪subset = orig.ident == "sc_67"), ident = "0")]))
res3_1 = rowSums(as.matrix(GetAssayData(subset(seu_intd_wt_AZ, subset = orig.
↪ident == "sc_101"), slot = "counts")[, WhichCells(subset(seu_intd_wt_AZ,␣
↪subset = orig.ident == "sc_101"), ident = "0")]))
res4_1 = rowSums(as.matrix(GetAssayData(subset(seu_intd_wt_AZ, subset = orig.
↪ident == "sc_103"), slot = "counts")[, WhichCells(subset(seu_intd_wt_AZ,␣
↪subset = orig.ident == "sc_103"), ident = "0")]))

sec1_1 = rowSums(as.matrix(GetAssayData(subset(seu_intd_wt_AZ, subset = orig.
↪ident == "sc_26_combined"), slot = "counts")[,␣
↪WhichCells(subset(seu_intd_wt_AZ, subset = orig.ident == "sc_26_combined"),␣
↪ident = "1")]))
sec2_1 = rowSums(as.matrix(GetAssayData(subset(seu_intd_wt_AZ, subset = orig.
↪ident == "sc_67"), slot = "counts")[, WhichCells(subset(seu_intd_wt_AZ,␣
↪subset = orig.ident == "sc_67"), ident = "1")]))
sec3_1 = rowSums(as.matrix(GetAssayData(subset(seu_intd_wt_AZ, subset = orig.
↪ident == "sc_101"), slot = "counts")[, WhichCells(subset(seu_intd_wt_AZ,␣
↪subset = orig.ident == "sc_101"), ident = "1")]))
```

```
sec4_1 = rowSums(as.matrix(GetAssayData(subset(seu_intd_wt_AZ, subset = orig.
↪ident == "sc_103"), slot = "counts")[, WhichCells(subset(seu_intd_wt_AZ,␣
↪subset = orig.ident == "sc_103"), ident = "1")]))

wt_sec_v_rec[,1:8] = c(res1_1, res2_1, res3_1, res4_1, sec1_1, sec2_1, sec3_1,␣
↪sec4_1)
wt_sec_v_rec_red[,1:6] = c(res1_1, res2_1, res3_1 + res4_1, sec1_1, sec2_1,␣
↪sec3_1 + sec4_1)
```

[ ]:
```
colnames(wt_sec_v_rec) = c(rep("res",4), rep("sec",4))
colnames(wt_sec_v_rec_red) = c(rep("res",3), rep("sec",3))
rownames(wt_sec_v_rec) = names(res1_1)
rownames(wt_sec_v_rec_red) = names(res1_1)
```

[ ]:
```
zone=as.factor(c(rep("res",4), rep("sec",4)))
design <- model.matrix(~zone)#+insertion)

#check design matrix isn't singular
print(paste("determinant of XT*X of design matrix is: ",␣
↪det(t(design)%*%(design))))

#making contrast matrix for tests of interest
my.contrasts <- makeContrasts(s1_v_s2=zonesec, levels=design)
wt_zone_edger_1 =  edgeR_2_sample(wt_sec_v_rec, "res", "sec", c(1,2,3,4),␣
↪c(5,6,7,8), annotations, design, my.contrasts)
```

[ ]:
```
zone=as.factor(c(rep("res",3), rep("sec",3)))
sort=as.factor(c("u","u","s","u","u","s"))
design <- model.matrix(~zone + sort)#+insertion)

#check design matrix isn't singular
print(paste("determinant of XT*X of design matrix is: ",␣
↪det(t(design)%*%(design))))

#making contrast matrix for tests of interest
my.contrasts <- makeContrasts(s1_v_s2=zonesec, levels=design)
wt_zone_edger_red =  edgeR_2_sample(wt_sec_v_rec_red, "res", "sec", c(1,2,3),␣
↪c(4,5,6), annotations, design, my.contrasts)
```

[ ]:
```
head(wt_zone_edger_red[wt_zone_edger_red$FDR < .05,],20)
```

[ ]:
```
write.csv(wt_zone_edger_1, "../data/for_figures/wt_zone_edger_4_21_22.csv")
write.csv(wt_zone_edger_red, "../data/for_figures/wt_zone_edger_red_4_21_22.
↪csv")
```

[ ]:
```
wt_zone_edger_1[wt_zone_edger_1$genes=="AT1G01610",]
```

[ ]:
```
cluster = "11"

mut_1_AZ <- subset(seu_intd_wt_mut, subset = orig.ident == "sc_27_combined")[,␣
 ↪WhichCells(subset(seu_intd_wt_mut, subset = orig.ident == "sc_27_combined"),␣
 ↪ident = cluster)]
mut_2_AZ <- subset(seu_intd_wt_mut, subset = orig.ident == "sc_68")[,␣
 ↪WhichCells(subset(seu_intd_wt_mut, subset = orig.ident == "sc_68"), ident =␣
 ↪cluster)]
KE_1_AZ <- subset(seu_intd_wt_mut, subset = orig.ident == "sc_102")[,␣
 ↪WhichCells(subset(seu_intd_wt_mut, subset = orig.ident == "sc_102"), ident =␣
 ↪cluster)]
KE_2_AZ <- subset(seu_intd_wt_mut, subset = orig.ident == "sc_104")[,␣
 ↪WhichCells(subset(seu_intd_wt_mut, subset = orig.ident == "sc_104"), ident =␣
 ↪cluster)]
```

[ ]:
```
mut_1_seu = SCTransform(mut_1_AZ)
mut_2_seu = SCTransform(mut_2_AZ)
KE_1_seu = SCTransform(KE_1_AZ)
KE_2_seu = SCTransform(KE_2_AZ)
```

[ ]:
```
seu_intd_mut_AZ = seu_integrate(mut_1_seu, mut_2_seu, KE_1_seu, KE_2_seu,␣
 ↪filename = "AZ_only_mut_3_1_22", nfeatures = 3000)
```

[ ]:
```
resolution = .1
set.seed(42)
DefaultAssay(seu_intd_mut_AZ) <- "integrated"
options(repr.plot.width=12, repr.plot.height=12)
# Run the standard workflow for visualization and clustering
seu_intd_mut_AZ <- ScaleData(seu_intd_mut_AZ, verbose = FALSE)
seu_intd_mut_AZ <- RunPCA(seu_intd_mut_AZ, npcs = 100, verbose = FALSE, approx␣
 ↪= FALSE)
seu_intd_mut_AZ <- FindNeighbors(seu_intd_mut_AZ, dims = 1:20, verbose = FALSE)
seu_intd_mut_AZ<- FindClusters(seu_intd_mut_AZ, resolution = resolution,␣
 ↪algorithm = 3, verbose = FALSE)
seu_intd_mut_AZ <- RunUMAP(seu_intd_mut_AZ, reduction = "pca", dims = 1:20,␣
 ↪verbose = FALSE)
```

[ ]:
```
options(repr.plot.width= 10, repr.plot.height=10)
plot = DimPlot(seu_intd_mut_AZ, reduction = "umap", label = TRUE, pt.size = 4)
print(plot)
ggsave(file="../data/for_figures/UMAPs/AZ_mut_UMAP.png", plot=plot, width=10,␣
 ↪height=10)
```

[ ]:
```
DefaultAssay(seu_intd_mut_AZ) <- "RNA"

#get pseudobulk for each cluster to compare with kwak data
```

```
pbs_mut = list()
count = 1
for (l in levels(seu_intd_mut_AZ@meta.data$seurat_clusters)) {
    pbs_mut[[count]] = rowSums(as.matrix(GetAssayData(seu_intd_mut_AZ, slot =␣
↪"counts")[,WhichCells(seu_intd_mut_AZ, ident = l)]))
    count = count + 1
}
```

[ ]:

```
#convert pseudobulk to TPM
count = 1
for (c in pbs_mut) {
    pbs_mut[[count]] = data.frame(pbs_mut[[count]])/sum(data.
↪frame(pbs_mut[[count]]))*1000000
    rns = rownames(pbs_mut[[count]])
    pbs_mut[[count]] = pbs_mut[[count]][order(rns),, drop = FALSE]
    count = count + 1
}
```

[ ]:

```
#secession
kwak_ptpms_raw=read.csv("../data/counts/kwak_ptpms.csv")
rownames(kwak_ptpms_raw) = kwak_ptpms_raw$X
kwak_ptpms = kwak_ptpms_raw
kwak_ptpms[,c(1,3,4)] =NULL

#secession
#set dataset
dataset = kwak_ptpms

cors_spearman = vector()
count = 1

seu_intd_mut_AZ@meta.data$kwak_cor = NULL

for (cluster in c(1:length(levels(seu_intd_mut_AZ@meta.data$seurat_clusters)))){
    test = cbind(pbs_mut[[cluster]][intersect(rownames(pbs_mut[[cluster]]),␣
↪rownames(dataset)),],dataset[intersect(rownames(pbs_mut[[cluster]]),␣
↪rownames(dataset)),])
    cors_spearman[count] = cor(log(test[,1]+.1), log(test[,2]+.1), method =␣
↪"spearman")
    count = count + 1
}

for (i in c(1:length(levels(seu_intd_mut_AZ@meta.data$seurat_clusters)))){
    seu_intd_mut_AZ@meta.data$kwak_cor[seu_intd_mut_AZ@meta.
↪data$seurat_clusters == toString(i-1)] = cors_spearman[i]
}
```

```
plot = FeaturePlot(seu_intd_mut_AZ,  features = "kwak_cor", pt.size = 4, cols =␣
 ↪c("light gray", "red"))
print(plot)
ggsave(file="../data/for_figures/UMAPs/AZ_mut_sec_UMAP.png", plot=plot,␣
 ↪width=10, height=10)
```

[ ]:
```
#residuum
kwak_ptpms_raw=read.csv("../data/counts/kwak_ptpms.csv")
rownames(kwak_ptpms_raw) = kwak_ptpms_raw$X
kwak_ptpms = kwak_ptpms_raw
kwak_ptpms[,c(1,2,4)] =NULL

#residuum
#set dataset
dataset = kwak_ptpms

cors_spearman = vector()
count = 1

seu_intd_mut_AZ@meta.data$kwak_cor = NULL

for (cluster in c(1:length(levels(seu_intd_mut_AZ@meta.data$seurat_clusters)))){
    test = cbind(pbs_mut[[cluster]][intersect(rownames(pbs_mut[[cluster]]),␣
 ↪rownames(dataset)),],dataset[intersect(rownames(pbs_mut[[cluster]]),␣
 ↪rownames(dataset)),])
    cors_spearman[count] = cor(log(test[,1]+.1), log(test[,2]+.1), method =␣
 ↪"spearman")
    count = count + 1
}

for (i in c(1:length(levels(seu_intd_mut_AZ@meta.data$seurat_clusters)))){
    seu_intd_mut_AZ@meta.data$kwak_cor[seu_intd_mut_AZ@meta.
 ↪data$seurat_clusters == toString(i-1)] = cors_spearman[i]
}

plot = FeaturePlot(seu_intd_mut_AZ,  features = "kwak_cor", pt.size = 4, cols =␣
 ↪c("light gray", "red"))
print(plot)
ggsave(file="../data/for_figures/UMAPs/AZ_mut_res_UMAP.png", plot=plot,␣
 ↪width=10, height=10)
```

[ ]:
```
DefaultAssay(seu_intd_mut_AZ) <- "RNA"
mut_sec_v_rec = data.frame(matrix(ncol = 8, nrow␣
 ↪=dim(seu_intd_mut_AZ@assays$RNA)[1]))
```

```
mut_sec_v_rec_red = data.frame(matrix(ncol = 6, nrow␣
↪=dim(seu_intd_mut_AZ@assays$RNA)[1]))


res1_1 = rowSums(as.matrix(GetAssayData(subset(seu_intd_mut_AZ, subset = orig.
↪ident == "sc_27_combined"), slot = "counts")[,␣
↪WhichCells(subset(seu_intd_mut_AZ, subset = orig.ident == "sc_27_combined"),␣
↪ident = "0")]))
res2_1 = rowSums(as.matrix(GetAssayData(subset(seu_intd_mut_AZ, subset = orig.
↪ident == "sc_68"), slot = "counts")[, WhichCells(subset(seu_intd_mut_AZ,␣
↪subset = orig.ident == "sc_68"), ident = "0")]))
res3_1 = rowSums(as.matrix(GetAssayData(subset(seu_intd_mut_AZ, subset = orig.
↪ident == "sc_102"), slot = "counts")[, WhichCells(subset(seu_intd_mut_AZ,␣
↪subset = orig.ident == "sc_102"), ident = "0")]))
res4_1 = rowSums(as.matrix(GetAssayData(subset(seu_intd_mut_AZ, subset = orig.
↪ident == "sc_104"), slot = "counts")[, WhichCells(subset(seu_intd_mut_AZ,␣
↪subset = orig.ident == "sc_104"), ident = "0")]))

sec1_1 = rowSums(as.matrix(GetAssayData(subset(seu_intd_mut_AZ, subset = orig.
↪ident == "sc_27_combined"), slot = "counts")[,␣
↪WhichCells(subset(seu_intd_mut_AZ, subset = orig.ident == "sc_27_combined"),␣
↪ident = "1")]))
sec2_1 = rowSums(as.matrix(GetAssayData(subset(seu_intd_mut_AZ, subset = orig.
↪ident == "sc_68"), slot = "counts")[, WhichCells(subset(seu_intd_mut_AZ,␣
↪subset = orig.ident == "sc_68"), ident = "1")]))
sec3_1 = rowSums(as.matrix(GetAssayData(subset(seu_intd_mut_AZ, subset = orig.
↪ident == "sc_102"), slot = "counts")[, WhichCells(subset(seu_intd_mut_AZ,␣
↪subset = orig.ident == "sc_102"), ident = "1")]))
sec4_1 = rowSums(as.matrix(GetAssayData(subset(seu_intd_mut_AZ, subset = orig.
↪ident == "sc_104"), slot = "counts")[, WhichCells(subset(seu_intd_mut_AZ,␣
↪subset = orig.ident == "sc_104"), ident = "1")]))


mut_sec_v_rec[,1:8] = c(res1_1, res2_1, res3_1, res4_1, sec1_1, sec2_1, sec3_1,␣
↪sec4_1 )
mut_sec_v_rec_red[,1:6] = c(res1_1, res2_1, res3_1 + res4_1, sec1_1, sec2_1,␣
↪sec3_1 + sec4_1 )
```

[ ]:
```
colnames(mut_sec_v_rec) = c(rep("res",4), rep("sec",4))
colnames(mut_sec_v_rec_red) = c(rep("res",3), rep("sec",3))
rownames(mut_sec_v_rec) = names(res1_1)
rownames(mut_sec_v_rec_red) = names(res1_1)
```

[ ]:
```
zone=as.factor(c(rep("res",4), rep("sec",4)))
design <- model.matrix(~zone)#+insertion)
```

```
#check design matrix isn't singular
print(paste("determinant of XT*X of design matrix is: ",␣
↪det(t(design)%*%(design))))

#making contrast matrix for tests of interest
my.contrasts <- makeContrasts(s1_v_s2=zonesec, levels=design)
mut_zone_edger_1 =  edgeR_2_sample(mut_sec_v_rec, "res", "sec", c(1,2,3,4),␣
↪c(5,6,7,8), annotations, design, my.contrasts)
```

[ ]:
```
#combined sorted samples
zone=as.factor(c(rep("res",3), rep("sec",3)))
sort=as.factor(c("u","u","s","u","u","s"))
design <- model.matrix(~zone + sort)#+insertion)

#check design matrix isn't singular
print(paste("determinant of XT*X of design matrix is: ",␣
↪det(t(design)%*%(design))))

#making contrast matrix for tests of interest
my.contrasts <- makeContrasts(s1_v_s2=zonesec, levels=design)
mut_zone_edger_red =  edgeR_2_sample(mut_sec_v_rec_red, "res", "sec", c(1,2,3),␣
↪c(4,5,6), annotations, design, my.contrasts)
```

[ ]:
```
head(mut_zone_edger_red[mut_zone_edger_red$FDR<.05,])
```

[ ]:
```
write.csv(mut_zone_edger_1, "../data/for_figures/mut_zone_edger_4_21_22.csv")
write.csv(mut_zone_edger_red, "../data/for_figures/mut_zone_edger_red_4_21_22.
↪csv")
```

[ ]:
```
kwak = read.csv("../data/for_figures/KWAK_data.csv")
rownames(kwak) = kwak[,1]
kwak = kwak[,c(5:10)]
colnames(kwak) = c("res","res","res","sec","sec","sec")
kwak = kwak[c(1:33602),]
```

[ ]:
```
zone=as.factor(c(rep("res",3), rep("sec",3)))
design <- model.matrix(~zone)#+insertion)

#check design matrix isn't singular
print(paste("determinant of XT*X of design matrix is: ",␣
↪det(t(design)%*%(design))))

#making contrast matrix for tests of interest
my.contrasts <- makeContrasts(s1_v_s2=zonesec, levels=design)
```

[ ]:
```
kwak_edger_1 =  edgeR_2_sample(kwak, "res", "sec", c(1,2,3), c(4,5,6),␣
↪annotations, design, my.contrasts)
```

[ ]:
```
write.csv(kwak_edger_1, "../data/for_figures/kwak_edger_4_21_22.csv")
```

[ ]:
```
#takes a list of Seurat objects with SCT transform run
seu_integrate <- function(..., filename, nfeatures){
    seu.list <- list(...) # THIS WILL BE A LIST STORING EVERYTHING:

    ref.genes = rownames(seu.list[[1]]@assays$RNA)
    assay_list <- rep("SCT", length(seu.list))


    # integration
    rc.features <- SelectIntegrationFeatures(object.list = seu.list, nfeatures␣
↪= nfeatures)
    rc.features <- rc.features[(!c(grepl("ATMG",rc.features) | grepl("ATCG",rc.
↪features) | rc.features%in%proto_list))]

    seu.list <- PrepSCTIntegration(object.list = seu.list, anchor.features = rc.
↪features, verbose = TRUE, assay = assay_list)
    seu.list <- lapply(X = seu.list, FUN = RunPCA, verbose = FALSE, features =␣
↪rc.features)
    rc.anchors <- FindIntegrationAnchors(object.list = seu.list, normalization.
↪method = "SCT", anchor.features = rc.features, verbose = TRUE, reference=1,␣
↪reduction = "rpca")

    to_integrate <- Reduce(intersect, lapply(rc.anchors@object.list, rownames))
    # integrate data and keep full geneset

    rc.integrated <- IntegrateData(anchorset = rc.anchors,  features.to.
↪integrate = to_integrate, normalization.method = "SCT", verbose = TRUE)
    rc.integrated <- RunPCA(rc.integrated, npcs = 50, verbose = FALSE, approx =␣
↪FALSE)

    #save object
    saveRDS(rc.integrated, file = paste("../data/intd_seu_objects/",filename,".
↪rds", sep = ""))
    return(rc.integrated)
#    }
}
```
