## Supplemental Code for "Arabidopsis uses a molecular grounding mechanism and a biophysical circuit breaker to limit floral abscission signaling": notebook_07_sc_vs_bulk_RNA_seq.pdf

### WB07_sc_vs_bulk_RNA_seq

June 30, 2022

[1]:
```
#THIS SCRIPT CALCULATES THE INTERSECTION OF DE GENES OF PRIOR BULK COMPARISOIN␣
↪OF WT AND MUTANT COMPARED TO SINGLE-CELL ANALYSIS

library(edgeR)
library(here)
source(here("R_functions","edgeR_function.R"))

annotations = read.csv("R_functions/gene_descriptions.csv", header = F)
colnames(annotations) = c("gene_id", "description")
annotations$gene_id = substr(annotations$gene_id, 1, 9)
```

```
Loading required package: limma

here() starts at /home/robotmessenger810/sc_analysis/code
```

[2]:
```
sessionInfo()
```

```
R version 3.6.3 (2020-02-29)
Platform: x86_64-conda_cos6-linux-gnu (64-bit)
Running under: Ubuntu 20.04.2 LTS

Matrix products: default
BLAS/LAPACK: /home/robotmessenger810/anaconda3/envs/r_3/lib/libopenblasp-r0.3.9.so

locale:
 [1] LC_CTYPE=en_US.UTF-8       LC_NUMERIC=C
 [3] LC_TIME=en_US.UTF-8        LC_COLLATE=en_US.UTF-8
 [5] LC_MONETARY=en_US.UTF-8    LC_MESSAGES=en_US.UTF-8
 [7] LC_PAPER=en_US.UTF-8       LC_NAME=C
 [9] LC_ADDRESS=C               LC_TELEPHONE=C
[11] LC_MEASUREMENT=en_US.UTF-8 LC_IDENTIFICATION=C

attached base packages:
[1] stats     graphics  grDevices utils     datasets  methods   base

other attached packages:
[1] here_0.1     edgeR_3.28.1 limma_3.42.2
```

```
loaded via a namespace (and not attached):
 [1] Rcpp_1.0.7      locfit_1.5-9.4  lattice_0.20-45 fansi_0.5.0
 [5] rprojroot_2.0.2 digest_0.6.29   utf8_1.2.2      crayon_1.4.2
 [9] IRdisplay_0.7.0 grid_3.6.3      repr_1.1.0      lifecycle_1.0.1
[13] jsonlite_1.7.2  evaluate_0.14   pillar_1.6.4    rlang_0.4.12
[17] uuid_0.1-4      vctrs_0.3.8     ellipsis_0.3.2  IRkernel_1.1
[21] tools_3.6.3     compiler_3.6.3  base64enc_0.1-3 pbdZMQ_0.3-6
[25] htmltools_0.5.0
```

[ ]:
```
#bulk samples
bulk = counts_to_reads_df("../data/bulk_data/Col_v_h3h3_bulk_counts_first" )

#remove no_feature, ambiguous, etc reads
bulk = bulk[1:(dim(bulk)[1]-5),]

#account for factors in experiment
phenotype=as.factor(c("wt", "wt", "wt", "wt", "wt", "wt", "mut", "mut", "mut",␣
↪"mut", "mut", "mut"))
batch=as.factor(c(0,0,0,1,1,1,0,0,0,1,1,1))
design <- model.matrix(~phenotype+batch)#+insertion)

#double check design matrix isn't singular
print(paste("determinant of XT*X of design matrix is: ",␣
↪det(t(design)%*%(design))))

#making contrast matrix for tests of interest
my.contrasts <- makeContrasts(s1_v_s2=phenotypewt, levels=design)

#run edger
bulk_edger_2 =  edgeR_2_sample(bulk, "WT", "mut", c(1,2,3,4,5,6),␣
↪c(7,8,9,10,11,12), annotations, design, my.contrasts)
```

[ ]:
```
#pseudo bulk samples
pb = read.csv("../data/pseudo_bulk_data/AZ_pbs_4_19_22.csv")
rownames(pb) = pb[,1]
pb[,1] <- NULL

#account for factors in experiment
phenotype=as.factor(c("wt", "wt", "wt", "wt", "mut", "mut", "mut", "mut"))
method=as.factor(c(0,0,1,1,0,0,1,1))
design <- model.matrix(~phenotype+method)#+insertion)

#double check design matrix isn't singular
print(paste("determinant of XT*X of design matrix is: ",␣
↪det(t(design)%*%(design))))
```

```
#making contrast matrix for tests of interest
my.contrasts <- makeContrasts(s1_v_s2=phenotypewt, levels=design)

#run edger
pb_edger_1 =  edgeR_2_sample(pb, "WT", "mut", c(1,2,3,4), c(5,6,7,8),␣
↪annotations, design, my.contrasts)
```

[ ]:
```
sig_intersect_WT_up = intersect(pb_edger_1[pb_edger_1$FDR<.
↪05&pb_edger_1$logFC>1,]$genes, bulk_edger_2[bulk_edger_2$FDR<.
↪05&bulk_edger_2$logFC>1,]$genes)
length(sig_intersect_WT_up)
write.csv(sig_intersect_WT_up, "../data/pseudo_bulk_data/
↪pb_bulk_sig_intersect_WT_up.csv")
```
