## Supplemental Code for "Arabidopsis uses a molecular grounding mechanism and a biophysical circuit breaker to limit floral abscission signaling": notebook_08_fal.ipynb

```
# %% [code]
#THIS SCRIPT PLOTS A HEATMAP OF BULK RNA-SEQ GENE EXPRESSION VALUES FOR WT, MUTANT, AND THE FAL SUPPRESSORS USING THE LIST OF GENES GENERATED FROM SCRIPT 6

library(limma)
library(edgeR)
library(here)
library(ggplot2)
library(gplots)
source(here("R_functions","edgeR_function.R"))
annotations = read.csv("R_functions/gene_descriptions.csv", header = F)
colnames(annotations) = c("gene_id", "description")
annotations$gene_id = substr(annotations$gene_id, 1, 9)

library(pheatmap)

# %% [code]
sessionInfo()

# %% [code]
#bulk_pb_intersection
bulk_pb_intersection = read.csv("../data/pseudo_bulk_data/pb_bulk_sig_intersect_WT_up.csv")

# %% [code]
#bulk samples
bulk = counts_to_reads_df("../data/bulk_data/Col_h3h3_fals")

# %% [code]
colnames(bulk) = c("Col_1","fal7_1","Col_2","fal7_2","Col_3","fal7_3","Col_4","fal7_4","h3h3_1","fal3_1","h3h3_2","fal3_2", "h3h3_3","fal3_3","h3h3_4","fal3_4")

# %% [code]
#remove poor h3h3 sample
bulk2 = bulk[,-13]
head(bulk2)

# %% [code]
#make DGElist
x3 <- DGEList(counts = bulk2, genes = rownames(bulk2))


#reads per library uniquely mapped to a gene:

#make cpm and lcpm
cpm <- cpm(x3)
lcpm <- cpm(x3, log=TRUE)

#keep only genes that are expressed. "Expressed" here means counts observed in at least 3 samples
dim(x3)
keep.exprs <- rowMeans(cpm)>=.5
x3 <- x3[keep.exprs,, keep.lib.sizes=FALSE]
dim(x3) #compare to dim(x) above

#normalize data after removing low expressed genes
x3 <- calcNormFactors(x3)

#cpm, lcpm of normalized values
cpm <- cpm(x3)
lcpm <- cpm(x3, log=TRUE)

de_sub = lcpm[rownames(lcpm) %in% unlist(bulk_pb_intersection$x),]

# %% [code]
head(cpm)
write.csv(cpm, "../data/bulk_data/fal.csv")

# %% [code]

de_sub_av = cbind(rowMeans(de_sub[,c(1,3,5,7)]), rowMeans(de_sub[,c(9,11,14)]), rowMeans(de_sub[,c(2,4,6,8)]), rowMeans(de_sub[,c(10,12,13, 15)]))

# %% [code]
colnames(de_sub_av) = c("wt", "h3h3", "fal7", "fal3")

# %% [code]
de_sub_av = de_sub_av[,c(1,2,4,3)]
fals = de_sub_av

# %% [code]
dim(de_sub)

# %% [code]
#only WT up genes
mat = de_sub_av-rowMeans(de_sub_av[,(c(1,2))])
hmp = heatmap(mat, Rowv=NA, Colv=NA)
mat = mat[rev(hmp$rowInd),]
hmp = heatmap(mat, Rowv=NA, Colv=NA)

# %% [code]
pheatmap(mat,  cluster_rows=FALSE, cluster_cols=FALSE)

# %% [code]
for_plot = data.frame(matrix(ncol = 2, nrow = length(colSums(de_sub))))
colnames(for_plot) = c("genotype", "value")
for_plot$value= colSums(de_sub)
for_plot$genotype = substr(names(colSums(de_sub)),1,4)
for_plot
ggplot(data = for_plot, aes(x=genotype, y = value)) +geom_point()
```
