## Supplemental Code for "Arabidopsis uses a molecular grounding mechanism and a biophysical circuit breaker to limit floral abscission signaling": notebook_08_fal.pdf

### WB08_fal

June 30, 2022

[ ]:
```
#THIS SCRIPT PLOTS A HEATMAP OF BULK RNA-SEQ GENE EXPRESSION VALUES FOR WT,␣
↪MUTANT, AND THE FAL SUPPRESSORS USING THE LIST OF GENES GENERATED FROM␣
↪SCRIPT 6

library(limma)
library(edgeR)
library(here)
library(ggplot2)
library(gplots)
source(here("R_functions","edgeR_function.R"))
annotations = read.csv("R_functions/gene_descriptions.csv", header = F)
colnames(annotations) = c("gene_id", "description")
annotations$gene_id = substr(annotations$gene_id, 1, 9)

library(pheatmap)
```

[2]:
```
sessionInfo()
```

```
R version 3.6.3 (2020-02-29)
Platform: x86_64-conda_cos6-linux-gnu (64-bit)
Running under: Ubuntu 20.04.2 LTS

Matrix products: default
BLAS/LAPACK: /home/robotmessenger810/anaconda3/envs/r_3/lib/libopenblasp-r0.3.9.so

locale:
 [1] LC_CTYPE=en_US.UTF-8       LC_NUMERIC=C
 [3] LC_TIME=en_US.UTF-8        LC_COLLATE=en_US.UTF-8
 [5] LC_MONETARY=en_US.UTF-8    LC_MESSAGES=en_US.UTF-8
 [7] LC_PAPER=en_US.UTF-8       LC_NAME=C
 [9] LC_ADDRESS=C               LC_TELEPHONE=C
[11] LC_MEASUREMENT=en_US.UTF-8 LC_IDENTIFICATION=C

attached base packages:
[1] stats     graphics  grDevices utils     datasets  methods   base

other attached packages:
[1] pheatmap_1.0.12 gplots_3.1.1    ggplot2_3.3.5   here_0.1
```

```
[5] edgeR_3.28.1    limma_3.42.2

loaded via a namespace (and not attached):
 [1] Rcpp_1.0.7         RColorBrewer_1.1-2 pillar_1.6.4       compiler_3.6.3
 [5] bitops_1.0-7       base64enc_0.1-3    tools_3.6.3        digest_0.6.29
 [9] uuid_0.1-4         jsonlite_1.7.2     evaluate_0.14      lifecycle_1.0.1
[13] tibble_3.1.6       gtable_0.3.0       lattice_0.20-45    pkgconfig_2.0.3
[17] rlang_0.4.12       DBI_1.1.2          IRdisplay_0.7.0    IRkernel_1.1
[21] withr_2.4.3        repr_1.1.0         dplyr_1.0.7        caTools_1.18.0
[25] gtools_3.9.2       generics_0.1.1     vctrs_0.3.8        tidyselect_1.1.1
[29] locfit_1.5-9.4     rprojroot_2.0.2    grid_3.6.3         glue_1.6.0
[33] R6_2.5.1           fansi_0.5.0        pbdZMQ_0.3-6       purrr_0.3.4
[37] magrittr_2.0.1     scales_1.1.1       ellipsis_0.3.2     htmltools_0.5.0
[41] colorspace_2.0-2   KernSmooth_2.23-20 utf8_1.2.2         munsell_0.5.0
[45] crayon_1.4.2
```

[ ]:
```
#bulk_pb_intersection
bulk_pb_intersection = read.csv("../data/pseudo_bulk_data/
↪pb_bulk_sig_intersect_WT_up.csv")
```

[ ]:
```
#bulk samples
bulk = counts_to_reads_df("../data/bulk_data/Col_h3h3_fals")
```

[3]:
```
colnames(bulk) =␣
↪c("Col_1","fal7_1","Col_2","fal7_2","Col_3","fal7_3","Col_4","fal7_4","h3h3_1","fal3_1","h3
↪"h3h3_3","fal3_3","h3h3_4","fal3_4")
```

[ ]:
```
#remove poor h3h3 sample
bulk2 = bulk[,-13]
head(bulk2)
```

[ ]:
```
#make DGElist
x3 <- DGEList(counts = bulk2, genes = rownames(bulk2))


#reads per library uniquely mapped to a gene:

#make cpm and lcpm
cpm <- cpm(x3)
lcpm <- cpm(x3, log=TRUE)

#keep only genes that are expressed. "Expressed" here means counts observed in␣
↪at least 3 samples
dim(x3)
keep.exprs <- rowMeans(cpm)>=.5
x3 <- x3[keep.exprs,, keep.lib.sizes=FALSE]
```

```
dim(x3) #compare to dim(x) above

#normalize data after removing low expressed genes
x3 <- calcNormFactors(x3)

#cpm, lcpm of normalized values
cpm <- cpm(x3)
lcpm <- cpm(x3, log=TRUE)

de_sub = lcpm[rownames(lcpm) %in% unlist(bulk_pb_intersection$x),]
```

[ ]:
```
head(cpm)
write.csv(cpm, "../data/bulk_data/fal.csv")
```

[ ]:
```
de_sub_av = cbind(rowMeans(de_sub[,c(1,3,5,7)]), rowMeans(de_sub[,c(9,11,14)]),␣
↪rowMeans(de_sub[,c(2,4,6,8)]), rowMeans(de_sub[,c(10,12,13, 15)]))
```

[ ]:
```
colnames(de_sub_av) = c("wt", "h3h3", "fal7", "fal3")
```

[ ]:
```
de_sub_av = de_sub_av[,c(1,2,4,3)]
fals = de_sub_av
```

[ ]:
```
dim(de_sub)
```

[ ]:
```
#only WT up genes
mat = de_sub_av-rowMeans(de_sub_av[,(c(1,2))])
hmp = heatmap(mat, Rowv=NA, Colv=NA)
mat = mat[rev(hmp$rowInd),]
hmp = heatmap(mat, Rowv=NA, Colv=NA)
```

[ ]:
```
pheatmap(mat,  cluster_rows=FALSE, cluster_cols=FALSE)
```

[ ]:
```
for_plot = data.frame(matrix(ncol = 2, nrow = length(colSums(de_sub))))
colnames(for_plot) = c("genotype", "value")
for_plot$value= colSums(de_sub)
for_plot$genotype = substr(names(colSums(de_sub)),1,4)
for_plot
ggplot(data = for_plot, aes(x=genotype, y = value)) +geom_point()
```
